## Supplementary Information for "Evolutionary signatures of human cancers revealed via genomic analysis of over 35,000 patients"

Supplementary Material

Diletta Fontana[1,+], Ilaria Crespiatico[1,+], Valentina Crippa[1,+], Federica Malighetti[1], Matteo Villa[1], Fabrizio Angaroni[2,3], Luca De Sano[2], Andrea Aroldi[1,4], Marco Antoniotti[2,5], Giulio Caravagna[6], Rocco Piazza[1], Alex Graudenzi[2,5,7,*], Luca Mologni[1], and Daniele Ramazzotti[1,*]

[1]Department of Medicine and Surgery, University of Milano-Bicocca, Monza, Italy
[2]Department of Informatics, Systems and Communication, University of Milano-Bicocca, Milan, Italy
[3]Center of Computational Biology, Human Technopole, Milano, Italy
[4]Hematology and Clinical Research Unit, San Gerardo Hospital, Monza, Italy
[5]Bicocca Bioinformatics, Biostatistics and Bioimaging Centre – B4, Milan, Italy
[6]Department of Mathematics and Geosciences, University of Trieste, Trieste, Italy
[7]Institute of Molecular Bioimaging and Physiology, Consiglio Nazionale delle Ricerche (IBFM-CNR), Segrate, Milan, Italy
[+]These authors contributed equally and should be regarded as co-first authors.

**3 Results on simulations** **14**

**4 Results on cancer data** **29**

**5 ASCETIC models of Myeloid Malignancies and Early-stage Non-small cell lung cancer** **103**

**6 Validation of ASCETIC models on unseen datasets** **105**

32 Evolutionary Signatures for Skin Cutaneous Melanoma (Pan-Cancer Atlas) . 46
33 Survival analysis for Skin Cutaneous Melanoma (Pan-Cancer Atlas) . . . . . 47
34 Oncoprint for Skin Cutaneous Melanoma (Pan-Cancer Atlas) . . . . . . . . . 48
35 Evolutionary Signatures for Bladder Urothelial (MSK-MET) . . . . . . . . . 49
36 Survival analysis for Bladder Urothelial (MSK-MET) . . . . . . . . . . . . . 50
37 Oncoprint for Bladder Urothelial (MSK-MET) . . . . . . . . . . . . . . . . . 51
38 Evolutionary Signatures for Breast Ductal (HR+HER2-, MSK-MET) . . . . 52
39 Survival analysis for Breast Ductal (HR+HER2-, MSK-MET) . . . . . . . . 53
40 Oncoprint for Breast Ductal (HR+HER2-, MSK-MET) . . . . . . . . . . . . 54
41 Evolutionary Signatures for Breast Ductal (Triple Negative, MSK-MET) . . 55
42 Survival analysis for Breast Ductal (Triple Negative, MSK-MET) . . . . . . 56
43 Oncoprint for Breast Ductal (Triple Negative, MSK-MET) . . . . . . . . . . 57
44 Evolutionary Signatures for Breast Lobular (HR+, MSK-MET) . . . . . . . 58
45 Survival analysis for Breast Lobular (HR+, MSK-MET) . . . . . . . . . . . 59
46 Oncoprint for Breast Lobular (HR+, MSK-MET) . . . . . . . . . . . . . . . 60
47 Evolutionary Signatures for Cholangio Intrahepatic (MSK-MET) . . . . . . . 61
48 Survival analysis for Cholangio Intrahepatic (MSK-MET) . . . . . . . . . . . 62
49 Oncoprint for Cholangio Intrahepatic (MSK-MET) . . . . . . . . . . . . . . 63
50 Evolutionary Signatures for Colorectal (MSS, MSK-MET) . . . . . . . . . . 64
51 Survival analysis for Colorectal (MSS, MSK-MET) . . . . . . . . . . . . . . 65
52 Oncoprint for Colorectal (MSS, MSK-MET) . . . . . . . . . . . . . . . . . . 66
53 Evolutionary Signatures for Head and Neck Squamous (MSK-MET) . . . . . 67
54 Survival analysis for Head and Neck Squamous (MSK-MET) . . . . . . . . . 68
55 Oncoprint for Head and Neck Squamous (MSK-MET) . . . . . . . . . . . . . 69
56 Evolutionary Signatures for Hepatocellular Carcinoma (MSK-MET) . . . . . 70
57 Survival analysis for Hepatocellular Carcinoma (MSK-MET) . . . . . . . . . 71
58 Oncoprint for Hepatocellular Carcinoma (MSK-MET) . . . . . . . . . . . . . 72
59 Evolutionary Signatures for Melanoma Cutaneous (MSK-MET) . . . . . . . 73
60 Survival analysis for Melanoma Cutaneous (MSK-MET) . . . . . . . . . . . 74
61 Oncoprint for Melanoma Cutaneous (MSK-MET) . . . . . . . . . . . . . . . 75
62 Evolutionary Signatures for Melanoma Uveal (MSK-MET) . . . . . . . . . . 76
63 Survival analysis for Melanoma Uveal (MSK-MET) . . . . . . . . . . . . . . 77
64 Oncoprint for Melanoma Uveal (MSK-MET) . . . . . . . . . . . . . . . . . . 78

65 Evolutionary Signatures for Pancreatic Neuroendocrine (MSK-MET) . . . . 79
66 Survival analysis for Pancreatic Neuroendocrine (MSK-MET) . . . . . . . . 80
67 Oncoprint for Pancreatic Neuroendocrine (MSK-MET) . . . . . . . . . . . . 81
68 Evolutionary Signatures for Renal Clear Cell (MSK-MET) . . . . . . . . . . 82
69 Survival analysis for Renal Clear Cell (MSK-MET) . . . . . . . . . . . . . . 83
70 Oncoprint for Renal Clear Cell (MSK-MET) . . . . . . . . . . . . . . . . . . 84
71 Evolutionary Signatures for Small Bowel Cancer (MSK-MET) . . . . . . . . 85
72 Survival analysis for Small Bowel Cancer (MSK-MET) . . . . . . . . . . . . 86
73 Oncoprint for Small Bowel Cancer (MSK-MET) . . . . . . . . . . . . . . . . 87
74 Evolutionary Signatures for Small-Cell Lung Cancer (MSK-MET) . . . . . . 88
75 Survival analysis for Small-Cell Lung Cancer (MSK-MET) . . . . . . . . . . 89
76 Oncoprint for Small-Cell Lung Cancer (MSK-MET) . . . . . . . . . . . . . . 90
77 Evolutionary Signatures for Thyroid Papillary (MSK-MET) . . . . . . . . . 91
78 Survival analysis for Thyroid Papillary (MSK-MET) . . . . . . . . . . . . . 92
79 Oncoprint for Thyroid Papillary (MSK-MET) . . . . . . . . . . . . . . . . . 93
80 Evolutionary Signatures for Thyroid Poorly Differentiated (MSK-MET) . . . 94
81 Survival analysis for Thyroid Poorly Differentiated (MSK-MET) . . . . . . . 95
82 Oncoprint for Thyroid Poorly Differentiated (MSK-MET) . . . . . . . . . . . 96
83 Evolutionary Signatures for Uterine Endometrioid (MSK-MET) . . . . . . . 97
84 Survival analysis for Uterine Endometrioid (MSK-MET) . . . . . . . . . . . 98
85 Oncoprint for Uterine Endometrioid (MSK-MET) . . . . . . . . . . . . . . . 99
86 Evolutionary Signatures for Uterine Hypermutated (MSK-MET) . . . . . . . 100
87 Survival analysis for Uterine Hypermutated (MSK-MET) . . . . . . . . . . . 101
88 Oncoprint for Uterine Hypermutated (MSK-MET) . . . . . . . . . . . . . . . 102
89 Survival analysis for Diffuse Glioma - Subtypes Validation (GLASS) . . . . . 107
90 Oncoprint for Diffuse Glioma - Subtypes Validation (GLASS) . . . . . . . . 108
91 Evolutionary Signatures for Diffuse Glioma - Subtypes Validation (GLASS) . 109
92 Simulations considering the Diffuse Glioma dataset from the GLASS study . 110
93 Survival analysis for Diffuse Glioma - Subtypes Validation (MSK) . . . . . . 111
94 Oncoprint for Diffuse Glioma - Subtypes Validation (MSK) . . . . . . . . . . 112
95 Evolutionary Signatures for Diffuse Glioma - Subtypes Validation (MSK) . . 113
96 Survival analysis for Diffuse Glioma - Subtypes Validation (TCGA) . . . . . 114
97 Oncoprint for Diffuse Glioma - Subtypes Validation (TCGA) . . . . . . . . . 115
98 Evolutionary Signatures for Diffuse Glioma - Subtypes Validation (TCGA) . 116
99 Survival analysis for Acute Myeloid Leukemia - Subtypes Validation (Pan-Cancer Atlas) . . . . . . . . . . . . . . . . . . . . . . . . . . . . . . . . . 117
100 Oncoprint for Acute Myeloid Leukemia - Subtypes Validation (Pan-Cancer Atlas) . . . . . . . . . . . . . . . . . . . . . . . . . . . . . . . . . . . . 118

101 Survival analysis for Lung Adenocarcinoma - Subtypes Validation (Pan-Cancer Atlas) . . . . . . . . . . 119
102 Oncoprint for Lung Adenocarcinoma - Subtypes Validation (Pan-Cancer Atlas) 120
103 Survival analysis for Lung Squamous Cell Carcinoma - Subtypes Validation (Pan-Cancer Atlas) . . . . . . . . . . 121
104 Oncoprint for Lung Squamous Cell Carcinoma - Subtypes Validation (Pan-Cancer Atlas) . . . . . . . . . . 122
105 Survival analysis for Prostate Adenocarcinoma - Subtypes Validation (Pan-Cancer Atlas) . . . . . . . . . . 123
106 Oncoprint for Prostate Adenocarcinoma - Subtypes Validation (Pan-Cancer Atlas) . . . . . . . . . . 124
107 Survival analysis for Metastatic Prostate Cancer - Subtypes Validation (SU2C) 125
108 Oncoprint for Metastatic Prostate Cancer - Subtypes Validation (SU2C) . . 126

### List of Tables

1 Models of Myeloid Malignancies and Early-stage Non-small cell lung cancer . 104
2 Validation of ASCETIC model of Acute Myeloid Leukemia evolution . . . . . 106

### 1 Modelling time evolution in cancer

In this Section we provide the mathematical formulation of the ASCETIC framework presented in the main text. We will give a brief description of the input data, the basic definitions, and the details of the algorithmic procedure.

#### 1.1 Input genomic data

Timed data reporting follow-ups of distinct cancer patients during the progression would be the ideal input to directly observe cancer progression through time. Unfortunately, these kind of data are nowadays scarcely available. The available genomics data are mostly of 3 types: *(i) single-cell sequencing*, *(ii) bulk sequencing* NGS data from multiple biopsies of the same tumor, *(iii) NGS sequencing* of a single biopsy per tumor.

Given any of these data inputs on multiple patients, it is straightforward to obtain the cross-sectional dataset $\mathbf{D}$ input of our inference task, by first selecting a set of genomic alterations $V$ and, then, converting for each tumor the detected alterations into a binary matrix where 1 indicates that the alteration was observed in the patient and 0 that it was not observed. Formally, given $n$ somatic alteration and $m$ samples, we consider as input a *cross-sectional* dataset $\mathbf{D}$ of size $m \times n$, where an element of such matrix $d_{i,j}$ is equal to 1 if the j-th somatic alteration is observed in the i-th sample and 0 otherwise. Furthermore, we assume the $n$ alterations to have been selected as possible *candidate drivers*, e.g., as discussed in [1], and, thus, $n \ll m$.

Doing this, as for all the preprocessing steps that follow, caution must be paid when the data inputs are of types ($i$) and ($ii$): these inputs include multiple samples (single cells or bulks) per patient, which describe the temporal evolution of the tumor. Our aim is to combine observation from multiple patients in order to derive overall probabilistic patterns among the patients at the ensemble-level depicting relations of selective advantage. To guarantee the emergence of such relations, we propose to keep the number of samples per patient to similar numbers as, in the case of large imbalance, the statistical estimators may become biased, as the signals from the patients with bigger number of samples may prevail.

#### 1.2 The evolution of cancer as a graph

Given the matrix $\mathbf{D}$ described in the previous Section, we want to obtain a graphical representation of the evolution of the somatic alterations accumulating during the disease progression for a given cancer. To model cancer progression at the individual-level, different models were developed, see for example [2]. In this work we will rely on directed acyclic graphs (DAGs) [1].

In detail, given a sample $\phi$ (i.e., a row of $\mathbf{D}$) in which we observe the set of genomic alterations $V_\phi$ (e.g., a somatic point mutation in *TP53*), the DAG representing the time evolution of a single tumor is specified by $G_\phi = (V_\phi, A_\phi, t_{A_\phi})$, where $V_\phi = \{v \in n \mid\in d_{\phi,v} = 1\}$ are the nodes of the graph, $A_\phi$ is a set of directed arcs representing the parental relation between alterations (i.e., connecting earlier alterations to later ones), and $t_{A_phi} = \{t : A_\phi \to \mathbb{R}^+ \mid t(A_{i,j} = t_j - t_i)\}$ is a function that specifies the length of each arc representing the time between the two mutation. We notice that $G_\phi$ is guaranteed to contain no cycles as a consequence of time irreversibility, i.e., we have a total time order between all the observation for any given patient.

Suppose now to consider a set of patients, namely $\phi = 1 \ldots, m$, and for each of them let us define its *DAG* $G_\phi$ representing the evolution of the relative tumor. If we add to each of these DAGs a wild type genotype (i.e., a node that represents a sample with no alterations at time 0), it is possible to define the union graph between the $m$ samples $\mathbb{G}_\mathbb{P} = (\mathbb{V}_\mathbb{P}, \mathbb{A}_\mathbb{P})$, where, $\mathbb{V}_\mathbb{P}$ is the union of all nodes of the individual DAGs (i.e., $\mathbb{V}_\mathbb{P} = \bigcup_{\phi=1}^{m} V_\phi$), and $\mathbb{A}_\mathbb{P}$ is the union of the edges of the individual DAGs (i.e., $\mathbb{A}_\mathbb{P} = \bigcup_{\phi=1}^{m} A_\phi)$). Obliviously, this graph contains all the alterations $v$ and the arcs present in at least one of the $G_\phi$. We observe that, while the single patients are represented by a DAG, i.e., no loops are possible due to time irreversibility, when we combine them into an ensemble of patients [1], cycles may occur because of inconsistencies in the orderings between patients. An example of what just described is depicted in Figure 1 where we show 3 individual-level *DAGs* and their union graph $\mathbb{G}_\mathbb{P}$ representing the time observations at the ensemble-level.

To build $\mathbb{G}_\mathbb{P}$ from real data, we rely on methods for the estimation of the progression of tumors at the individual-level. As a matter of fact, several approaches have been developed recently to reconstruct cancer mutational trees from single-cell data (see [3, 4, 5]) and bulk data (see [5, 6, 7]). Such mutational tree models provide a partial order set among the alterations given as input, which we can be interpreted as a set of time constraints. Thence, we extend these intuitions to the case in which only one NGS sample per patient is available and we adopt an estimation of the prevalence of alterations in a tumor (i.e., the *cancer cell fractions*, CCF) as a measure of time: in this case, more prevalent alterations (with high CCF) are assumed to be occurring earlier in the tumor development. To name one method, [8] can be adopted for this purpose. It goes without saying that when more samples are available (cases of single cells and bulk data), the time estimation is more reliable, but even in the latter case, albeit without a perfect imputation of tumor clonality, we might nevertheless expect valuable information to emerge among the different patients. In this latter case we can obtain $G_\phi$ for a given patient as the *total temporal ordering* derived from the CCF for the sample, i.e., we set an arc from earlier nodes toward all the reachable later ones. $\mathbb{G}_\mathbb{P}$ can be directly derived from all the inferred $G_\phi$.

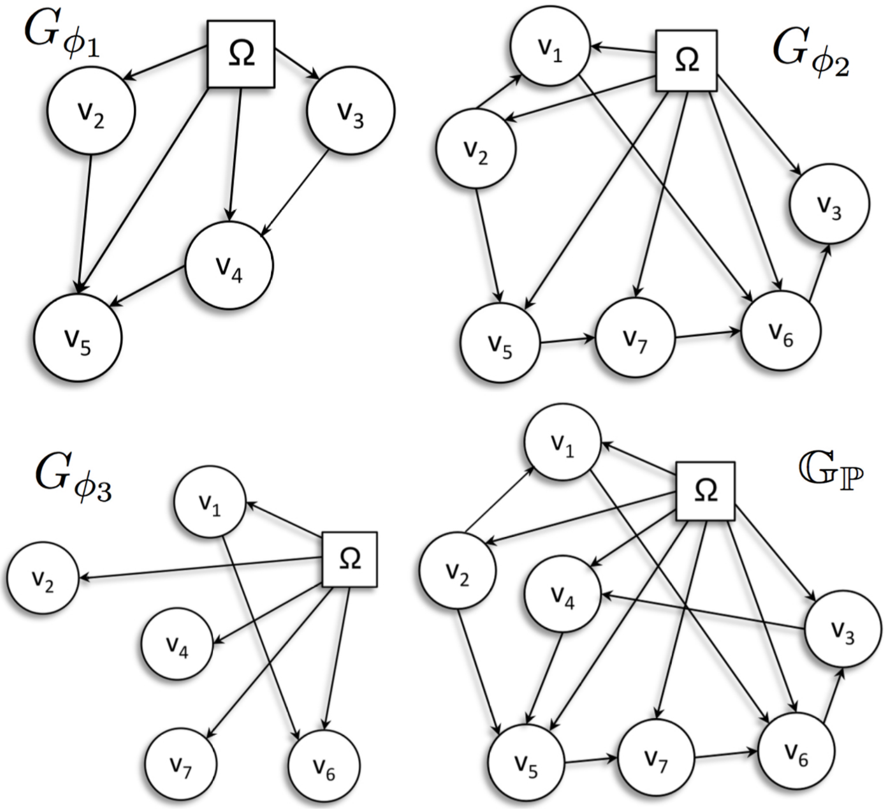

Figure 1: Example of ensemble-level temporal graph $\mathbb{G}_{\mathbb{P}}$ built from 3 individual-level *DAGs* $G_{\phi_1}$, $G_{\phi_2}$ and $G_{\phi_3}$. We recall that, while the individual-level *DAGs* cannot present cycles, their union graph may have some due to irregularities between the time orderings between the *DAGs*. This is shown in the figure where a loop among the nodes $V_3$, $V_4$, $V_5$, $V_6$ and $V_7$ arises.

#### 1.3 Irregularities in the time orderings

As outlined before, the ensemble-level temporal graph $\mathbb{G}_{\mathbb{P}}$ is a model of complex interpretation due to the presence of cycles and it cannot be directly used to depict a set of causal relations of selective advantage. Suppes Bayes Causal Networks (SBCN) [9, 10] have been proposed as a graphical representation capable of displaying causal relations among driver genes during cancer evolution.

Specifically, if we assume that a given genomic alteration is never lost, which is a reasonable assumption for mutations that provide a fitness advantage, then the cycles in the union graph $\mathbb{G}_{\mathbb{P}}$ could be considered as observed irregularities in the time orderings. By removing such irregularities we can transform the union graph $\mathbb{G}_{\mathbb{P}}$ into a SBCN. To this end, we need a proper measure to model irregularities in cancer.

In [11], Gupte and colleagues defined a measure of hierarchy within a directed graph and a score to quantify irregularities. Given a directed graph $G = (V, A)$, let us consider a ranking function $r : V \to \mathbb{N}$ for the nodes in $G$, such that $r(u) < r(v)$ expresses the fact that $u$ is "higher" in the hierarchy than $v$, i.e., the smaller $r(u)$ is, the more $u$ is an early event during tumor evolution. If $r(u) < r(v)$, then the arc $u \to v$ is expected and does not cause any "agony". Instead, if $r(u) \geq r(v)$ the arc $u \to v$ would cause agony because it would mean that $u$ has an incoming arc (an earlier event) from an alteration that occurred later in an other patient. Therefore, given a graph $G$ and a ranking $r$, the *agony* of each arc $(u, v)$ is defined as $\max\{r(u) - r(v) + 1, 0\}$, and the agony $a(G, r)$ of the whole graph for a given ranking $r$ is defined as the sum over all arcs:

$$a(G, r) = \sum_{(u,v) \in A} \max\{r(u) - r(v) + 1, 0\}.$$

In most cases (as in our case), the ranking $r$ is not explicitly provided. Thus, the objective becomes the one of finding a ranking (possibly non-unique) that minimizes the total agony of the graph. In this way, it is possible to compute the agony of any graph $G$ as:

$$a(G) = \min_{r} a(G, r).$$

This concept of hierarchy and agony could be directly applied to our problem. As a *DAG* implicitly induces a partial order over its nodes, it has always zero agony: the nodes of a *DAG* form a perfect hierarchy. For instance, in the *DAGs* $G_{\phi_1}, G_{\phi_2}$ and $G_{\phi_3}$ in Figure 1, it is sufficient to take the temporal ordering as ranking, i.e., $r(u) = t_u$ where $\langle \phi_i, u, t_u \rangle \in G_{\phi_i}$, in order to obtain agony equal to 0.

However, as already mentioned above, merging several *DAGs* to form ensemble-level temporal graph $\mathbb{G}_{\mathbb{P}}$ does not lead necessary to a *DAG* and agony can appear. The presence of any loop makes it impossible to find a ranking $r$ at 0 agony. In fact, any directed cycle containing $k$ arcs (and not sharing arcs with any other cycle) always generates agony equal to $k$ [11].

Although the number of possible rankings of a directed graph is exponential, Gupte and colleagues defined a polynomial-time algorithm for finding a ranking at minimum agony. They provided a linear-programming formulation and showed that (*i*) the dual problem has an optimal integral solution, and (*ii*) the optimal value obtained by maximizing the dual problem coincides with the minimum value of the primal. This finding led to an algorithm that decomposes the input graph $G$ into a *DAG* $D$ and a graph $H$ corresponding to the maximum (in terms of number of arcs) Eulerian subgraph[1] of $G$.

[1]An Eulerian graph is a graph in which the indegree of each node is equal to its outdegree.

Let $e$ be the number of arcs and $n$ the number of nodes of $G$, then the algorithm to compute such a decomposition takes $\mathcal{O}(e^2 n)$ time: it requires at each iteration to find a negative-weight cycle, which can be done by the Bellman-Ford algorithm [12] in $\mathcal{O}(en)$, while, in the worst case, the number of iterations is $e$. In a recent work, Tatti and colleagues [13] provided a faster algorithm for computing the agony of a directed graph. The algorithm has a theoretical bound of $\mathcal{O}(e^2)$ time, but it was empirically shown that such a bound is pessimistic and in practice the algorithm can scale to large graphs. In this work, we adopted this implementation.

### 2 Ensemble-level cancer progression

The semantic of SBCNs is grounded in the notion of probabilistic causality [14]. In fact, the relationships depicted in such networks encode two *necessary conditions* to claim causalities among pairs of observables, which, biologically, resemble the notion of selective advantage through cancer progression [9, 10, 15].

**Definition 1** (Probabilistic causation, [14]). *For any two events $u$ and $v$, occurring respectively at times $t_u$ and $t_v$, under the mild assumptions that $P(u), P(v) \in [0,1]$, the event $u$ is called a* prima facie cause *of $v$ if it occurs* before *and* raises the probability *of $u$, i.e.,*

$$\begin{aligned} \mathsf{TP}: &\quad t_u < t_v \\ \mathsf{PR}: &\quad P(v \mid u) > P(v \mid \overline{u}). \end{aligned}$$

The first condition of TP refers to the presence in the data of a temporal pattern where, if event $u$ is a cause of event $v$, it often occurs before its effect. This notion can be naturally reframed in terms of the existence of a temporal hierarchy among the two events as described in Section 1. Therefore, we can consistently assess the TP condition to be verified for any event $u$ toward its candidate effect $v$ when $r(u) < r(v)$.

The PR condition subsumes instead the presence of a statistically significant pattern of occurrence between pair of observables. In particular this adds a further meaning to the ranking between pair of events defined above: $u$ is a valid cause of $v$ if it occurs before and if a *significant pattern* is observed between the two events with the earlier occurrence of the cause raising the expectation of subsequently observing its effect.

In [15] the authors describe an efficient algorithm to infer SBCNs. The method first learns a poset of prima facie causes, i.e., the arcs in the network that verify Suppes' conditions and, hence, *may* be causal, and then infer the SBCN by maximum likelihood fit within this poset. This method is proved to be effective when all the parents of commod effects are conjunctive [15], but, although it may still approximate a good solution even if this assumption does not hold [9, 10, 15], to guarantee the inference of the true causal arcs, the input dataset needs to be *lifted* to add hypotheses of candidate parent sets which may not be conjunctively causing their common child [1, 15].

Unluckily, the definition of a good set of hypotheses is often hard, and the assumption of conjunctive causes is typically violated mostly because of cancer heterogeneity [10]. In this case, the statistical complications preventing a straightforward inference of the SBCN are due to the effect of the so-called *Simpson's paradox* [10]: in each observation we may observe a statistical trend, which yet may disappear or even revert when we consider the whole dataset. To overcome this hurdle, we extend Suppes' probabilistic causation in order to directly exploit the temporal information provided in the ensemble-level temporal graph $\mathbb{G}_{\mathbb{P}}$.

Specifically, for any pair of event $u$ and $v$, instead of estimating the prima facie conditions on the whole dataset, we limit our analysis to consider only the subset of the data where both $u$ and $v$ are observed together, i.e., when the Simpson's paradox is not expected to be happening. To this extend, we intuitively reframe our causal claims from "event $u$ is causing event $v$" to "the fact that event $u$ occurred before $v$ caused $u$ and $v$ to have been both observed together later on". This leads us to the formulation of an extended theory of probabilistic causality as follow.

**Definition 2** (Extended probabilistic causation)**.** *For any two events $u$ and $v$, occurring respectively at times $t_u$ and $t_v$, the event $u$ is called a* temporal prima facie cause *of $v$ if it occurs* before *and* raises the probability *of $u$ in terms of a significant temporal pattern as follow,*

$$\begin{aligned} &\mathsf{TP}: \quad r(u) < r(v) \\ &\mathsf{PR}: \quad P\left(u, v \mid t_u < t_v\right) > P\left(u, v \mid t_u \geq t_v\right). \end{aligned}$$

This intuitively aims at assessing if, when $u$ and $v$ both happened, $u$ consistently occurred first. If the two conditions hold, then we will say that $u$ is a candidate parent of $v$ and that the events may be involved in a relation of selective advantage with event $u$ occurring first and $v$ later on during tumor evolution.

#### 2.1 Agony-baSed Cancer EvoluTion InferenCe (ASCETIC)

Let us now consider a pair of events $u$ and $v$ for which we want to assess a claim for prima facie causality. We formulate this task as described in Definition 2 and, to do so, we consider now the two events, $\tau = [t_u < t_v]$ and $\rho = (u, v)$ as a proxy to estimate any causal relation among $u$ and $v$. To this end, we will say that $u$ is a prima facie cause of $v$ if $\tau$ is a prima facie cause of $\rho$. While the TP condition is directly assessable by means of the ranking metric we already discussed, PR needs further considerations.

PR for $\tau$ and $\rho$ implies [16]:

$$P(\tau, \rho) > P(\tau) \cdot P(\rho) \ .$$

This, expanding $\rho = (u, v)$, leads to

$$\begin{aligned}
&P(\tau, u, v) > P(\tau) \cdot P(u, v) \ , \\
&P(\tau, u, v) > [P(\tau, u, v) + P(\tau, \overline{u}, v) + P(\tau, u, \overline{v}) + P(\tau, \overline{u}, \overline{v})] \cdot P(u, v) \ .
\end{aligned}$$

We here aim at describing the timeline of each observable and, to do so, we assume that any considered event is occurring at a time $t \in [1, +\infty]$. Following this model, we can derive probabilistic relations involving our osservables. For instance, for $\tau$ and $\rho$ we can state:

$$\begin{cases}
P(\tau \mid u, v) = \alpha \in [0, 1], \\
P(\tau \mid \overline{u}, v) = 0, \\
P(\tau \mid u, \overline{v}) = 1, \\
P(\tau \mid \overline{u}, \overline{v}) = \beta = P(u, v) \cdot \alpha \in [0, 1],
\end{cases}$$

with the following interpretation. If we consider the subset of observations where both $u$ and $v$ are present, then the probability of $t_u < t_v$ is an (unknown) value $\alpha$ with values ranging in $[0, 1]$, which possibly depends on a causal relation between the two events. Similarly, if both $u$ and $v$ are yet to occur, the probability of $u$ to happen before $v$ when they both have occurred, still depends on $\alpha$. Furthermore, if $v$ occurred at a certain time in the past, but $u$ is yet to occur, then $t_u$ cannot be before $t_v$. On the contrary if $u$ occurred and $v$ did not, $t_u$ is before $t_v$ with probability 1.

Therefore, if we plug this probabilistic relations into Suppes' conditions and with some algebraic rearrangements, we get to the following.

$$\begin{aligned}
&\alpha \cdot P(u, v) > P(u, v) \cdot [\alpha \cdot P(u, v) + P(u, \overline{v}) + \beta \cdot P(\overline{u}, \overline{v})] \, , \\
&\alpha > \alpha \cdot P(u, v) + P(u) - P(u, v) + \beta \cdot [1 - P(u) - P(v) + P(u, v)] \, , \\
&\alpha > \alpha \cdot P(u, v) + P(u) - P(u, v) + \alpha \cdot P(u, v) \cdot [1 - P(u) - P(v) + P(u, v)] \, , \\
&\alpha - P(u) + P(u, v) > \alpha \cdot P(u, v) \, [P(\overline{u}) + P(\overline{v}) + P(u, v)] \, .
\end{aligned}$$

This leads us to,

$$\frac{\alpha - P(u) + P(u, v)}{\alpha \cdot P(u, v) \, [P(\overline{u}) + P(\overline{v}) + P(u, v)]} > 0 \, .$$

The denominator of this fraction is strictly greater than 0 if $\alpha > 0$, which leads us to the following condition:

**Algorithm 1** *Agony-baSed Cancer EvoluTion InferenCe* (ASCETIC)

1: **Input:** A *cross-sectional* dataset $\mathbb{D}$ of $n$ somatic alterations $V$ for $m$ distinct patients and an ensemble-level temporal graph $\mathbb{G}_{\mathbb{P}} = (\mathbb{V}_{\mathbb{P}}, \mathbb{A}_{\mathbb{P}})$ built from the $m$ patients.
2: [*Initialization*] Start from a complete graph $G$ connecting all the $n$ somatic alterations.
3: [*Minumum agony ranking*] Define a ranking $r$ over the $n$ somatic alterations at minimum agony from $\mathbb{G}_{\mathbb{P}}$, as done in [13]. Create from $G$, the minimum agony poset, a DAG $D_a$ by removing all the arcs from equal or higher ranked nodes to lower ranked ones.
4: [*Prima facie causal DAG*] Remove from $D_a$ all the arcs where the PR condition does not hold as for Definition 2. This leads us to the prima facie causal DAG $D_{pf}$.
5: [*Likelihood fit*] Filter out all spurious causes from $D_{pf}$ by likelihood fit with regularization over $\mathbb{D}$ to obtain the final progression model $\mathcal{D}$.
6: **Output:** the DAG $\mathcal{D}$.

$$\textit{PR:}\quad P\left(u, v \mid t_u < t_v\right) > P\left(u, v \mid t_u \geq t_v\right) \iff P\left(\tau \mid u, v\right) > P\left(u, \overline{v}\right).$$

We recall that such inequality is directly verifiable from the input data and considers only situations when both the observables are observed, hence, reducing the impact of Simpson's paradox, since our statistics are not performed on mixed populations. All the considerations above, allow us to define the Agony-baSed Cancer EvoluTion InferenCe (ASCETIC) framework to perform the inference of SBCNs (Algorithm 1).

The algorithm first creates a partially ordered set (poset) among the $n$ genomics alterations. This poset accounts for TP being computed from a ranking at minimum agony derived from the ensemble-level temporal graph $\mathbb{G}_{\mathbb{P}}$. Furthermore, it also accounts for temporal priority as defined above. It considers each pair of event $u$ and $v$ and estimates both $P\left(t_u < t_v \mid u, v\right)$ and $P\left(u, \overline{v}\right)$ from $\mathbb{D}$ and $\mathbb{G}_{\mathbb{P}}$. Once the poset is created, the final progression model is estimated by maximum likelihood.

If information about cell prevalence is available, e.g., when cancer cell fractions are provided, it is possible to also take this information into account in the estimation of the minimum agony ranking. In fact, intuitively, genes observed at very different cancer cell fractions, are more strongly indicators for a temporal order than genes with a close CCF. This might be done by weighting each observation in the agony computation (see [13]), but is left for future research.

### 3 Results on simulations

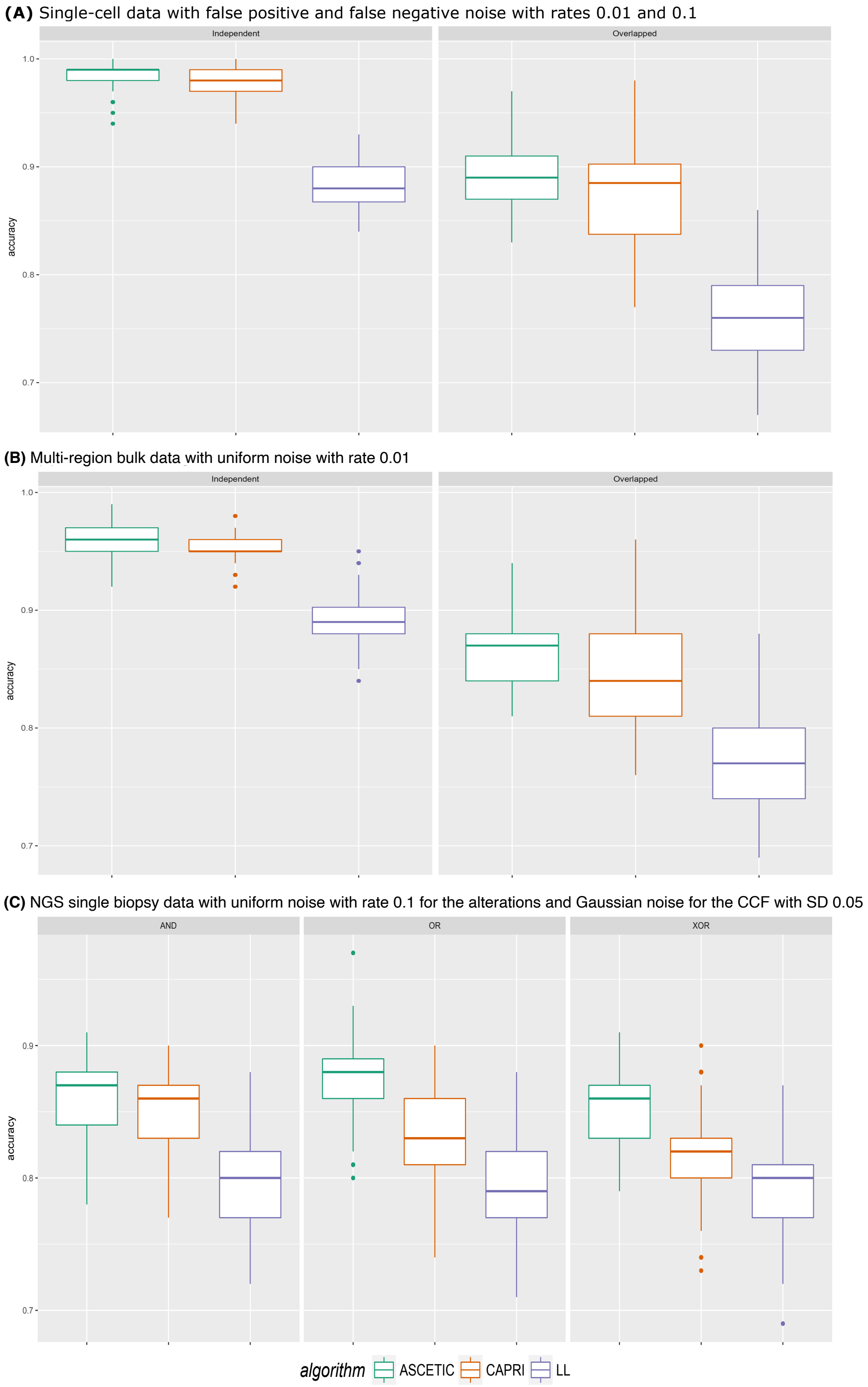

Figure 2: Results of simulations comparing the accuracy of ASCETIC, CAPRI algorithm [15] and the standard maximum likelihood fit approach for structure learning for a set of selected representative simulated scenarios. In panel $(A)$ we show simulations of single-cell data with false positive and false negative noise with rates 0.01 and 0.1. In panel $(B)$ we show results of simulations of multi-region bulk data with uniform noise with rate 0.01. Finally, in panel $(C)$ we show simulations of NGS single biopsy data with uniform noise with rate 0.1 for the alterations and Gaussian noise for the cancer cell fractions (CCF) with standard deviation SD 0.05.

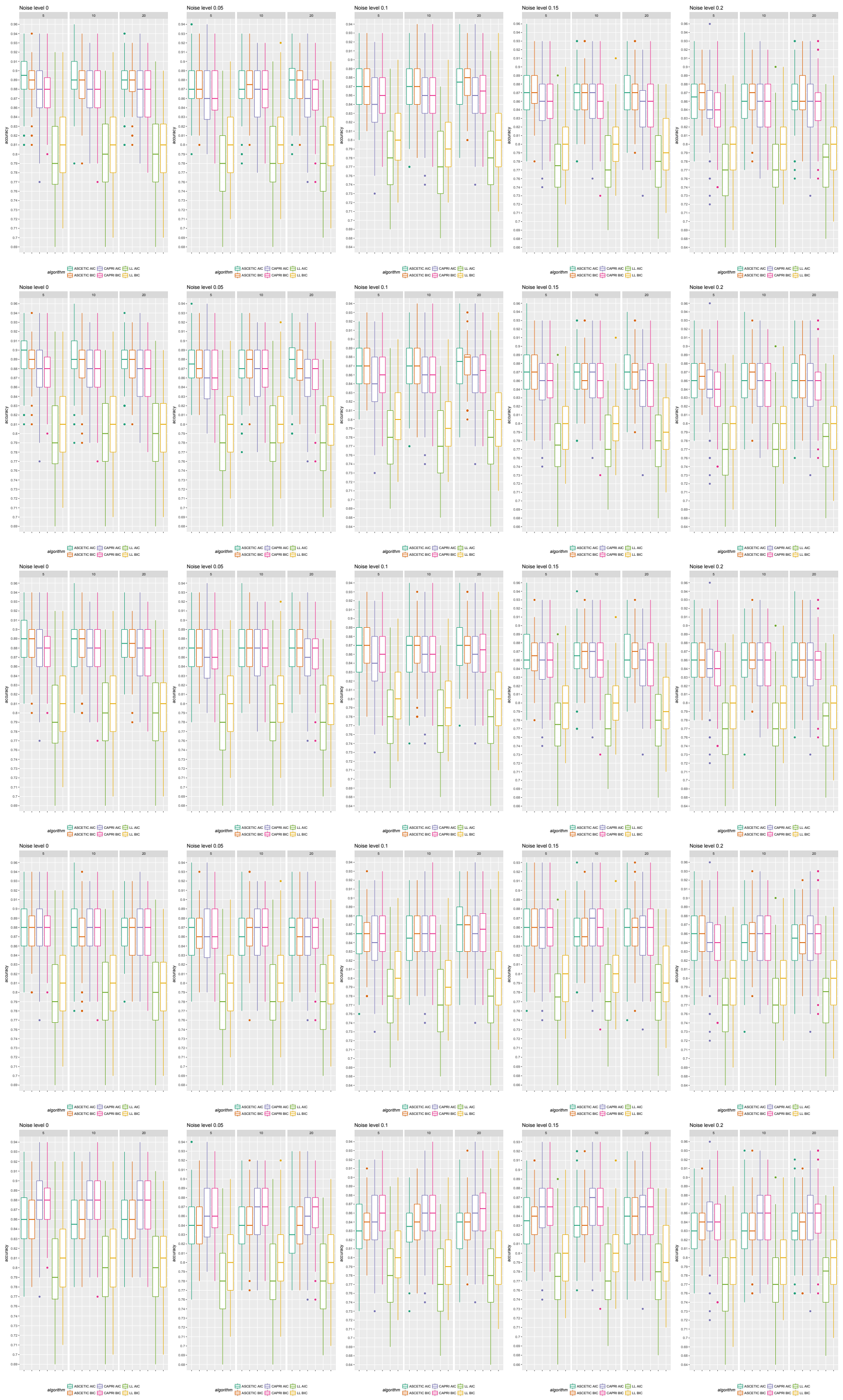

Figure 3: Results of the simulations for the agony method. Here we simulated NGS data from one biopsis with the AND parent sets. In each column we consider 5 noise levels, namely 0%, 5%, 10%, 15%, 20%, and datasets of 50, 100 and 200 patients. In each row we consider simulations of the cancer cell fractions at different noise levels, namely Gaussian samplig with standard deviation 0, 0.01, 0.05, 0.1and 0.2. ASCETIC is compared against CAPRI [15] and the standard likelihood fit with both AIC [17] and BIC [18] likelihood scores in terms of recall.

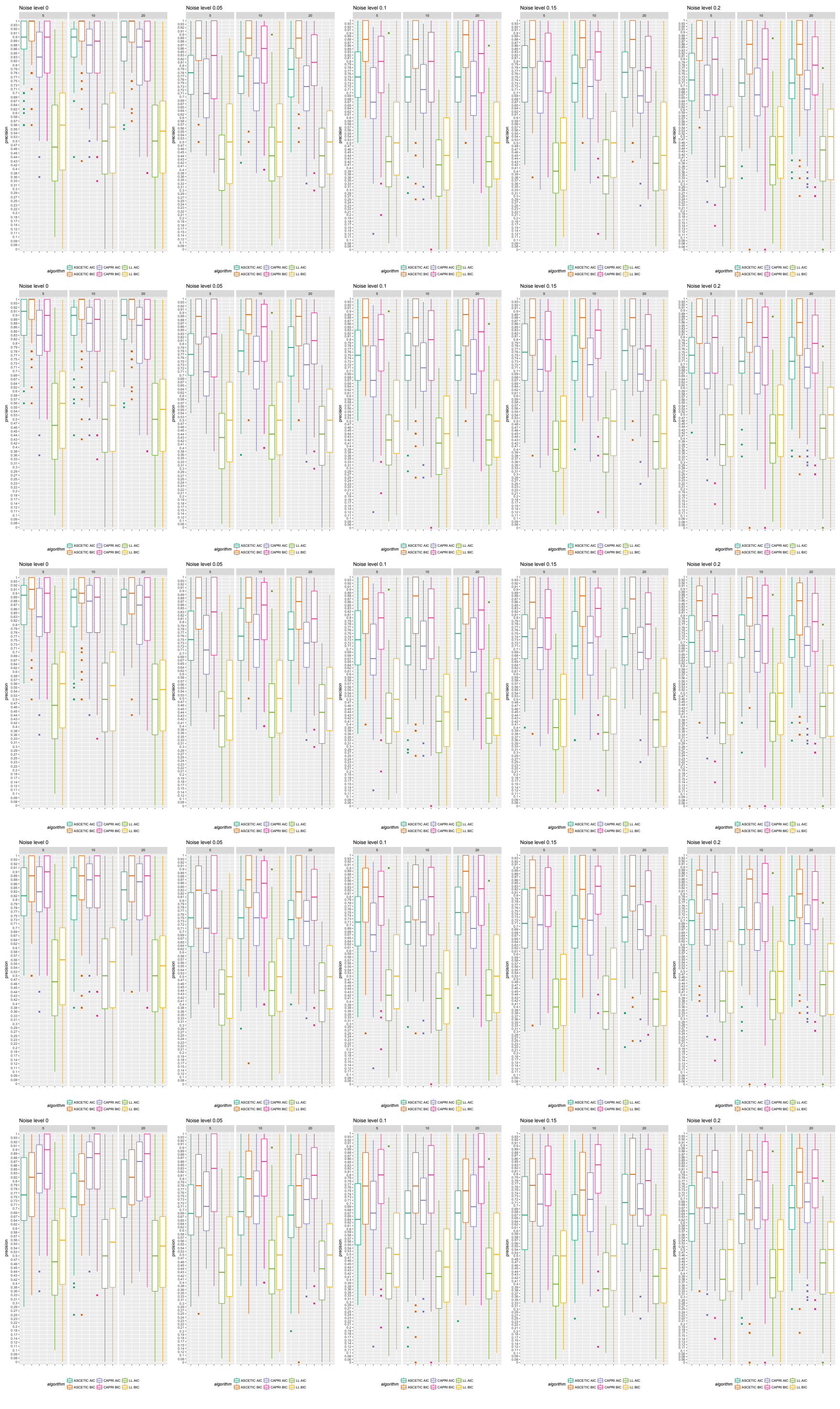

Figure 4: Results of the simulations for the agony method. Here we simulated NGS data from one biopsis with the AND parent sets. In each column we consider 5 noise levels, namely 0%, 5%, 10%, 15%, 20%, and datasets of 50, 100 and 200 patients. In each row we consider simulations of the cancer cell fractions at different noise levels, namely Gaussian samplig with standard deviation 0, 0.01, 0.05, 0.1and 0.2. ASCETIC is compared against CAPRI [15] and the standard likelihood fit with both AIC [17] and BIC [18] likelihood scores in terms of recall.

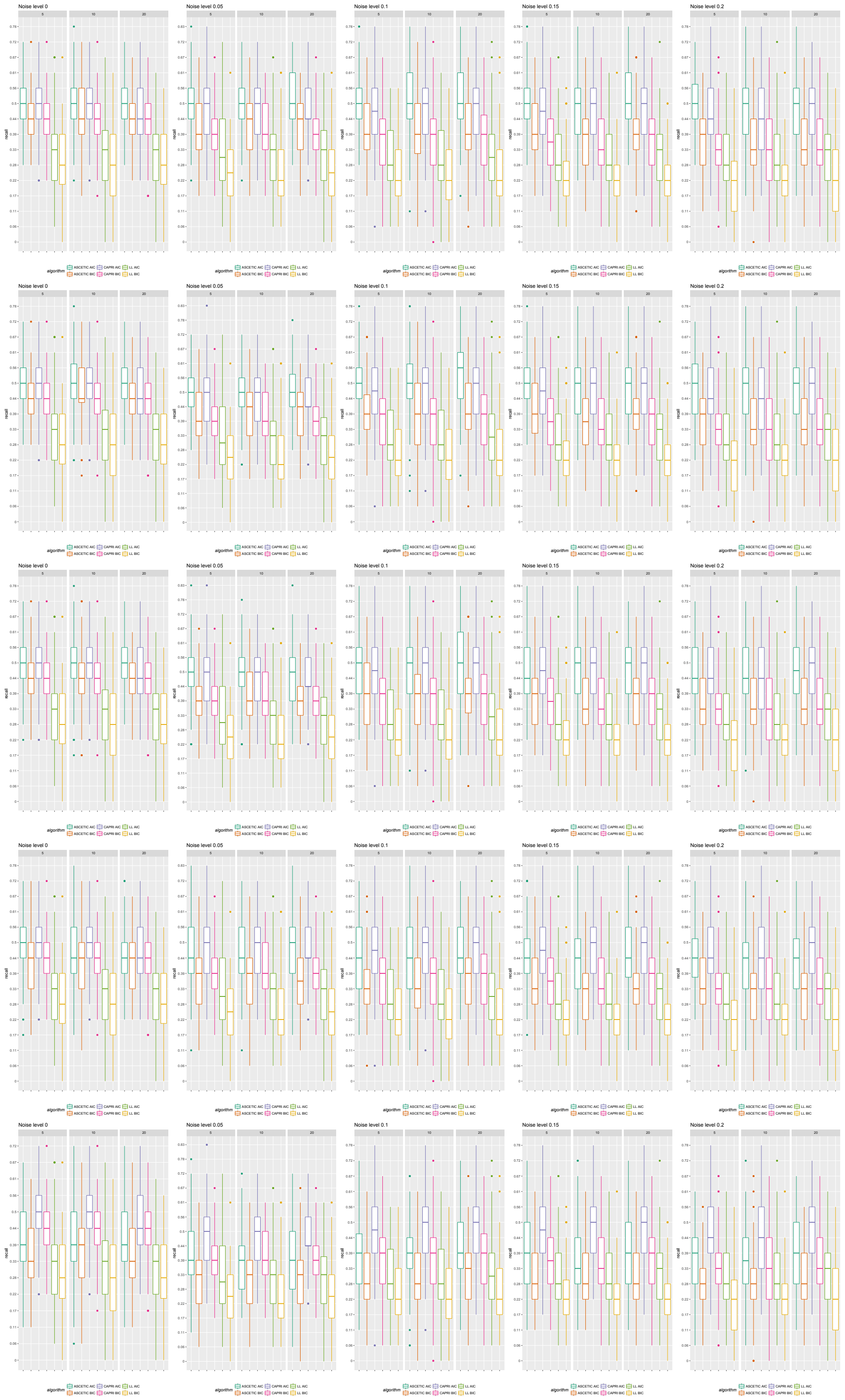

Figure 5: Results of the simulations for the agony method. Here we simulated NGS data from one biopsis with the AND parent sets. In each column we consider 5 noise levels, namely 0%, 5%, 10%, 15%, 20%, and datasets of 50, 100 and 200 patients. In each row we consider simulations of the cancer cell fractions at different noise levels, namely Gaussian samplig with standard deviation 0, 0.01, 0.05, 0.1and 0.2. ASCETIC is compared against CAPRI [15] and the standard likelihood fit with both AIC [17] and BIC [18] likelihood scores in terms of recall.

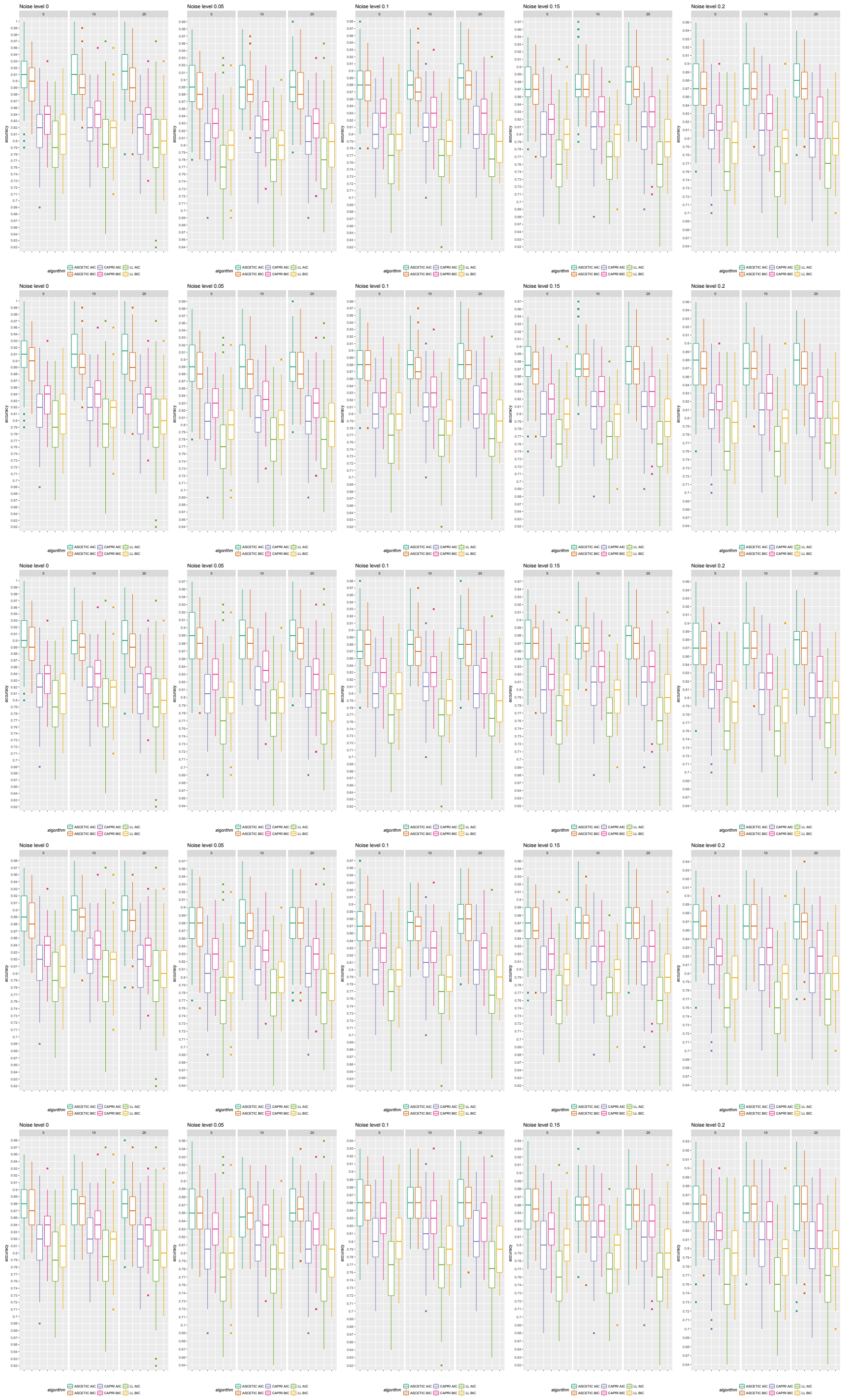

Figure 6: Results of the simulations for the agony method. Here we simulated NGS data from one biopsis with the OR parent sets. In each column we consider 5 noise levels, namely 0%, 5%, 10%, 15%, 20%, and datasets of 50, 100 and 200 patients. In each row we consider simulations of the cancer cell fractions at different noise levels, namely Gaussian samplig with standard deviation 0, 0.01, 0.05, 0.1and 0.2. ASCETIC is compared against CAPRI [15] and the standard likelihood fit with both AIC [17] and BIC [18] likelihood scores in terms of accuracy.

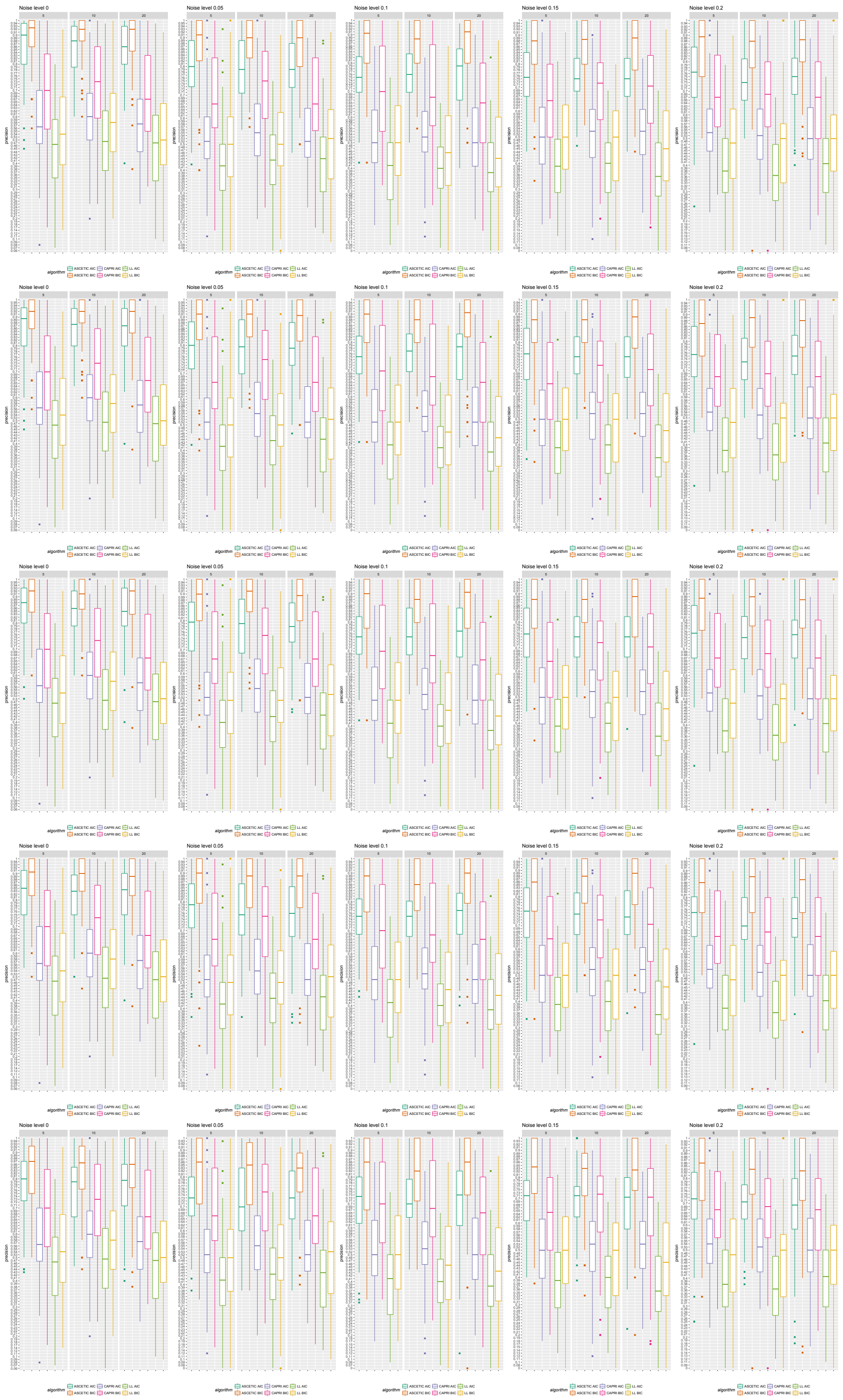

Figure 7: Results of the simulations for the agony method. Here we simulated NGS data from one biopsis with the OR parent sets. In each column we consider 5 noise levels, namely 0%, 5%, 10%, 15%, 20%, and datasets of 50, 100 and 200 patients. In each row we consider simulations of the cancer cell fractions at different noise levels, namely Gaussian samplig with standard deviation 0, 0.01, 0.05, 0.1and 0.2. ASCETIC is compared against CAPRI [15] and the standard likelihood fit with both AIC [17] and BIC [18] likelihood scores in terms of precision.

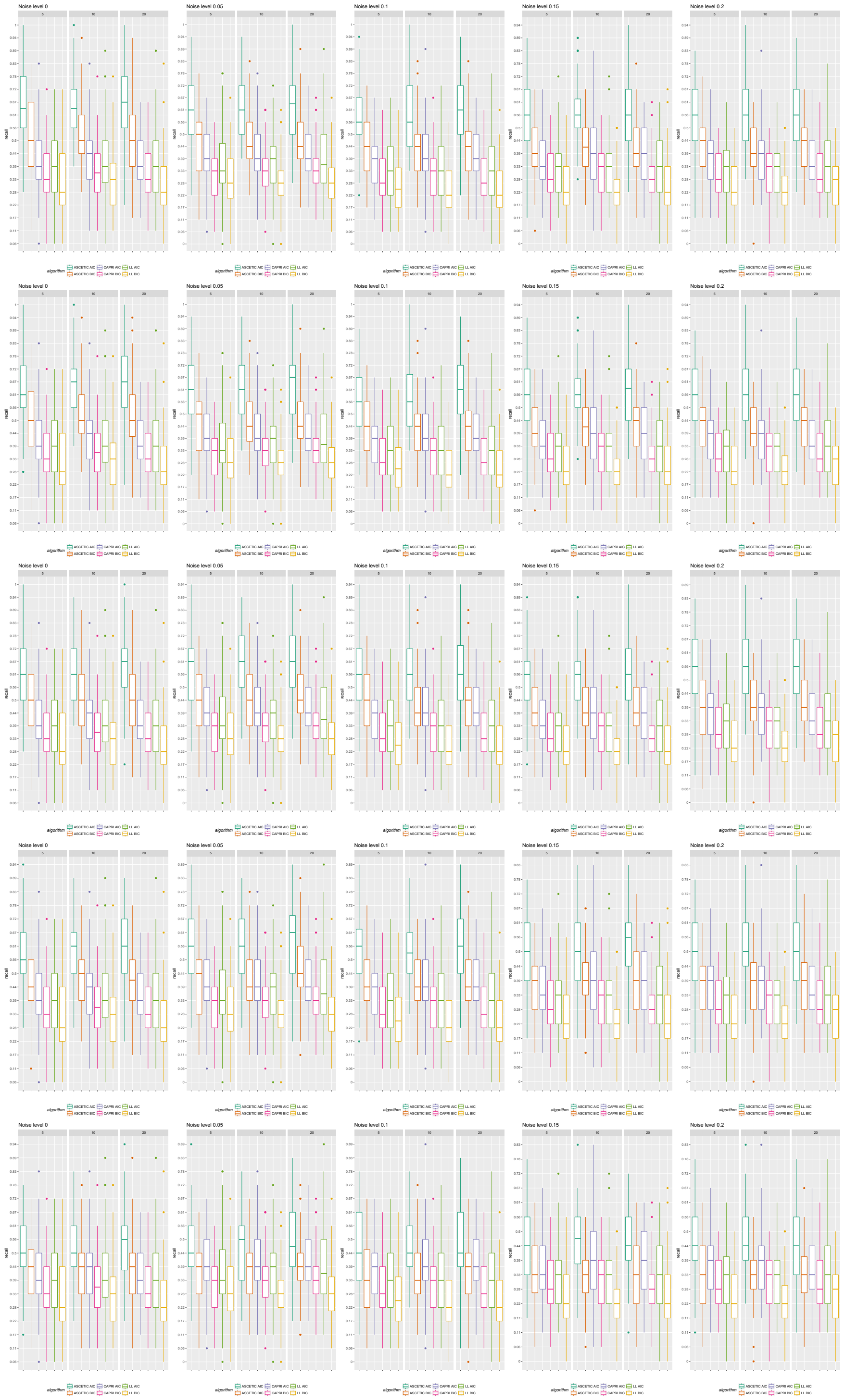

Figure 8: Results of the simulations for the agony method. Here we simulated NGS data from one biopsis with the OR parent sets. In each column we consider 5 noise levels, namely 0%, 5%, 10%, 15%, 20%, and datasets of 50, 100 and 200 patients. In each row we consider simulations of the cancer cell fractions at different noise levels, namely Gaussian samplig with standard deviation 0, 0.01, 0.05, 0.1and 0.2. ASCETIC is compared against CAPRI [15] and the standard likelihood fit with both AIC [17] and BIC [18] likelihood scores in terms of recall.

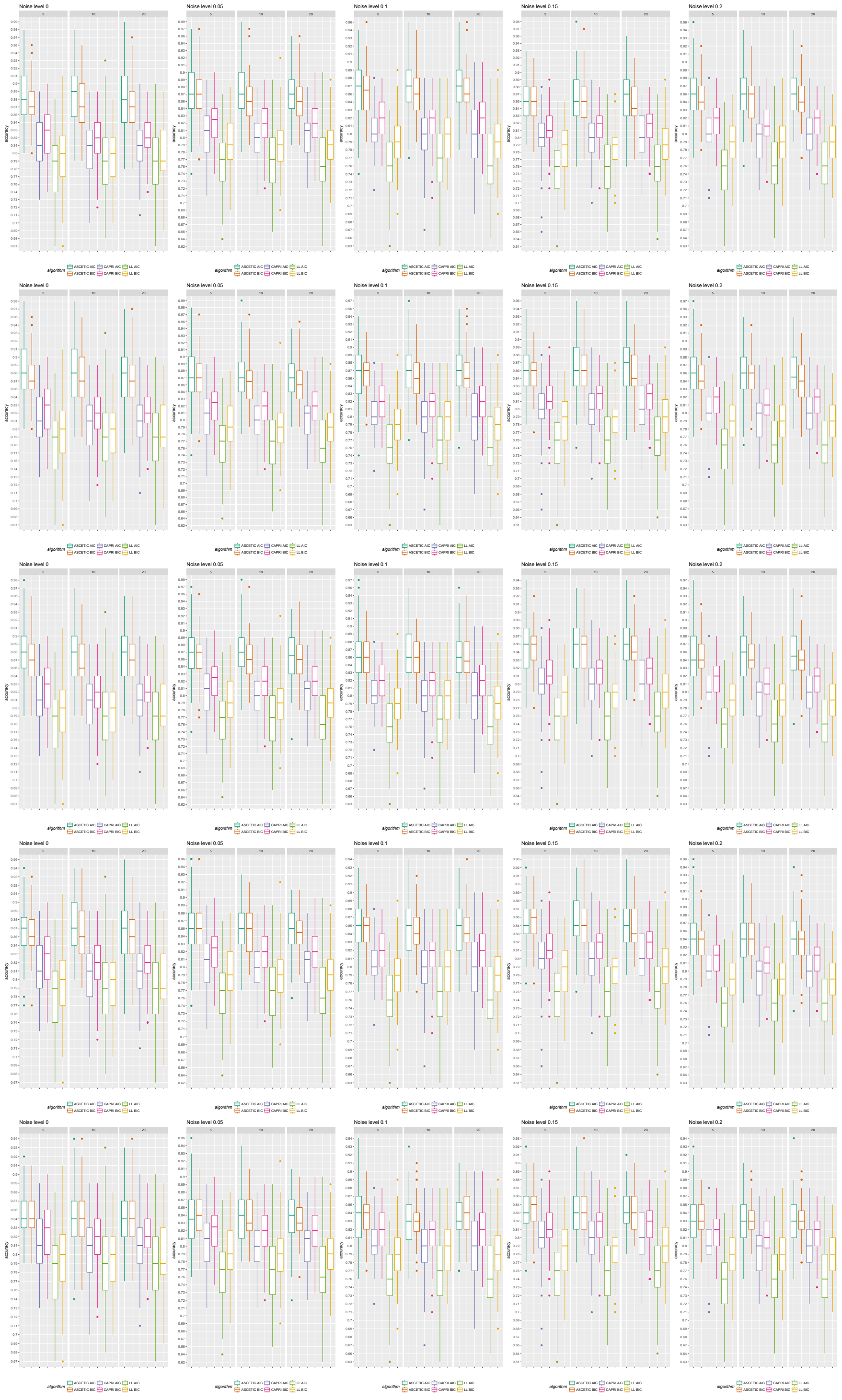

Figure 9: Results of the simulations for the agony method Here we simulated NGS data from one biopsis with the XOR parent sets. In each column we consider 5 noise levels, namely 0%, 5%, 10%, 15%, 20%, and datasets of 50, 100 and 200 patients. In each row we consider simulations of the cancer cell fractions at different noise levels, namely Gaussian samplig with standard deviation 0, 0.01, 0.05, 0.1and 0.2. ASCETIC is compared against CAPRI [15] and the standard likelihood fit with both AIC [17] and BIC [18] likelihood scores in terms of accuracy.

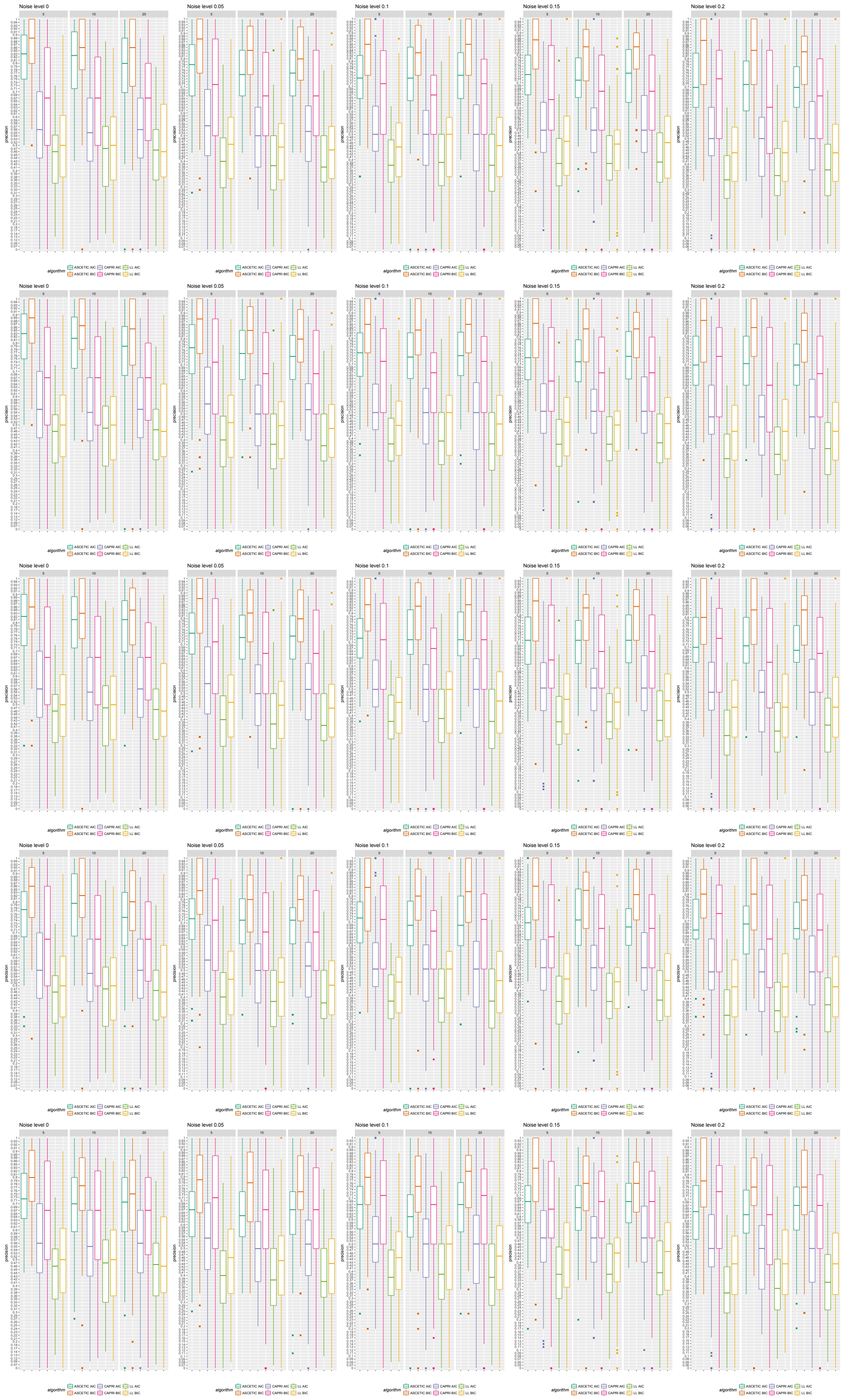

Figure 10: Results of the simulations for the agony method. Here we simulated NGS data from one biopsis with the XOR parent sets. In each column we consider 5 noise levels, namely 0%, 5%, 10%, 15%, 20%, and datasets of 50, 100 and 200 patients. In each row we consider simulations of the cancer cell fractions at different noise levels, namely Gaussian samplig with standard deviation 0, 0.01, 0.05, 0.1and 0.2. ASCETIC is compared against CAPRI [15] and the standard likelihood fit with both AIC [17] and BIC [18] likelihood scores in terms of precision.

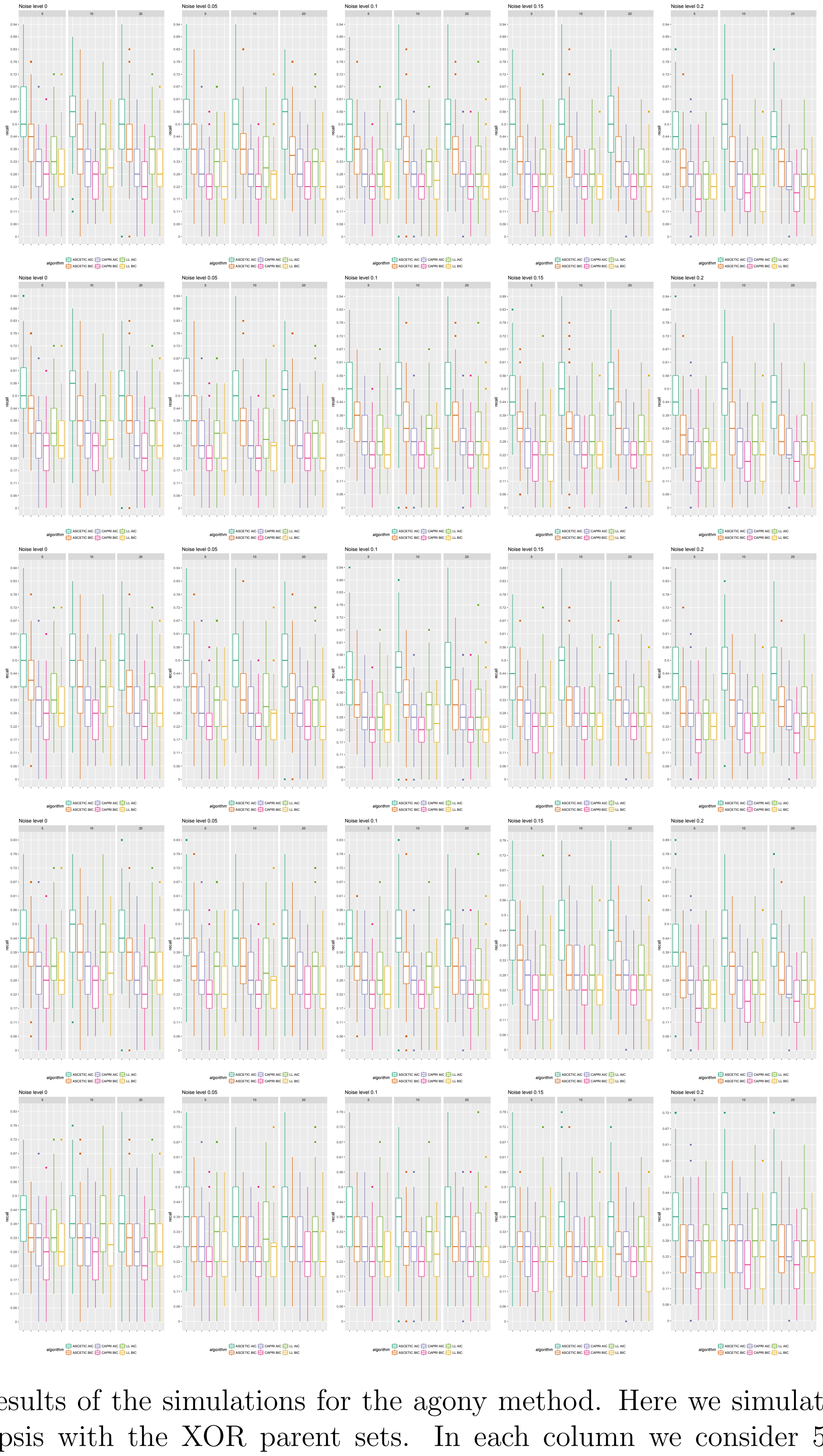

Figure 11: Results of the simulations for the agony method. Here we simulated NGS data from one biopsis with the XOR parent sets. In each column we consider 5 noise levels, namely 0%, 5%, 10%, 15%, 20%, and datasets of 50, 100 and 200 patients. In each row we consider simulations of the cancer cell fractions at different noise levels, namely Gaussian samplig with standard deviation 0, 0.01, 0.05, 0.1and 0.2. ASCETIC is compared against CAPRI [15] and the standard likelihood fit with both AIC [17] and BIC [18] likelihood scores in terms of recall.

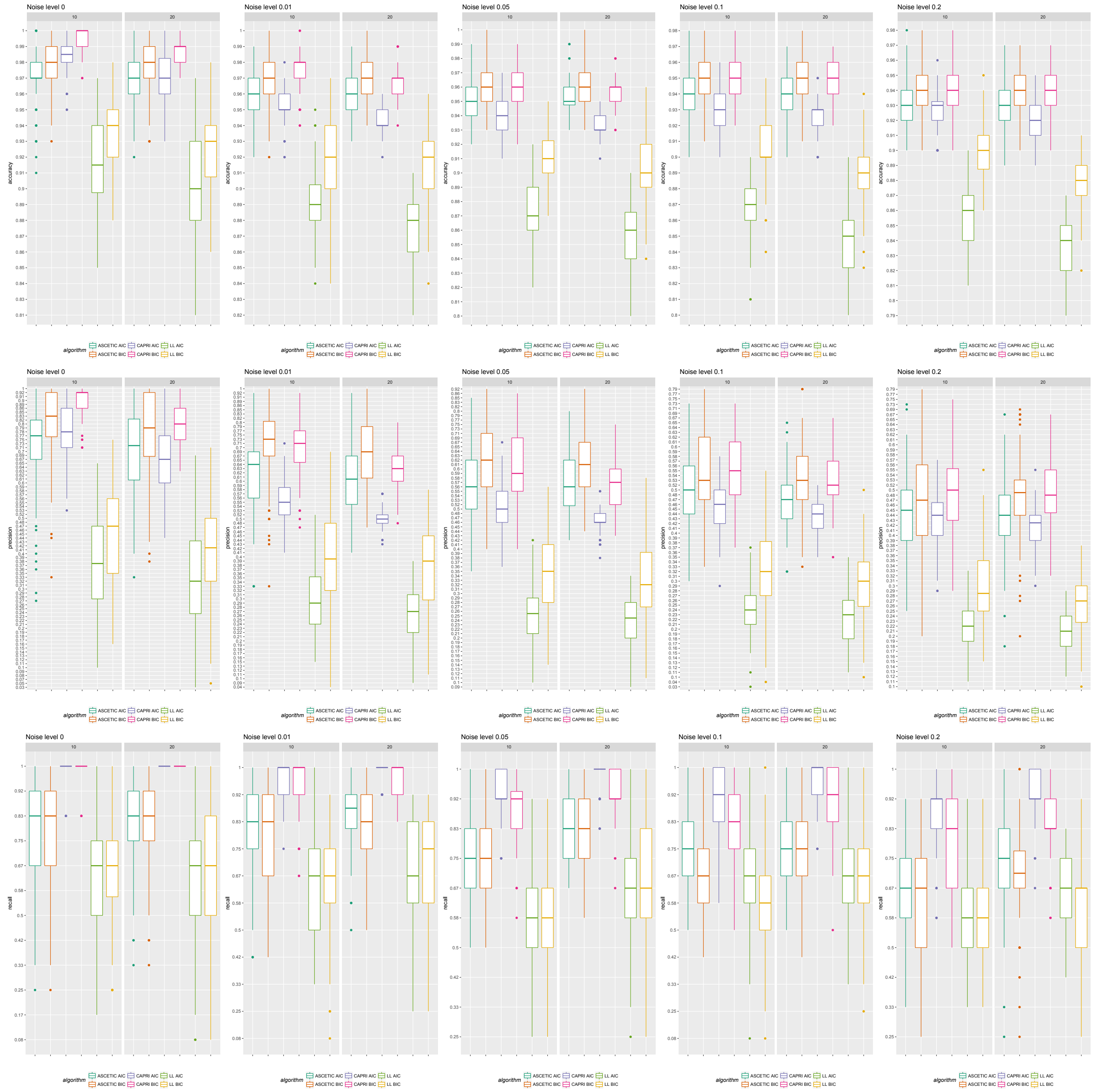

Figure 12: Results of the simulations for the agony method. Here we simulated NGS data from multiple biopses with 3 independent progressions and a total of 15 mutations. In each column we consider 5 noise levels, namely 0%, 1%, 5%, 10%, 20%, and datasets of 10 or 20 patients with 10 biopses each. In each row we show the results for different matrics, specifically for accuracy, precision and recall. ASCETIC is compared against CAPRI [15] and the standard likelihood fit with both AIC [17] and BIC [18] likelihood scores in terms of recall.

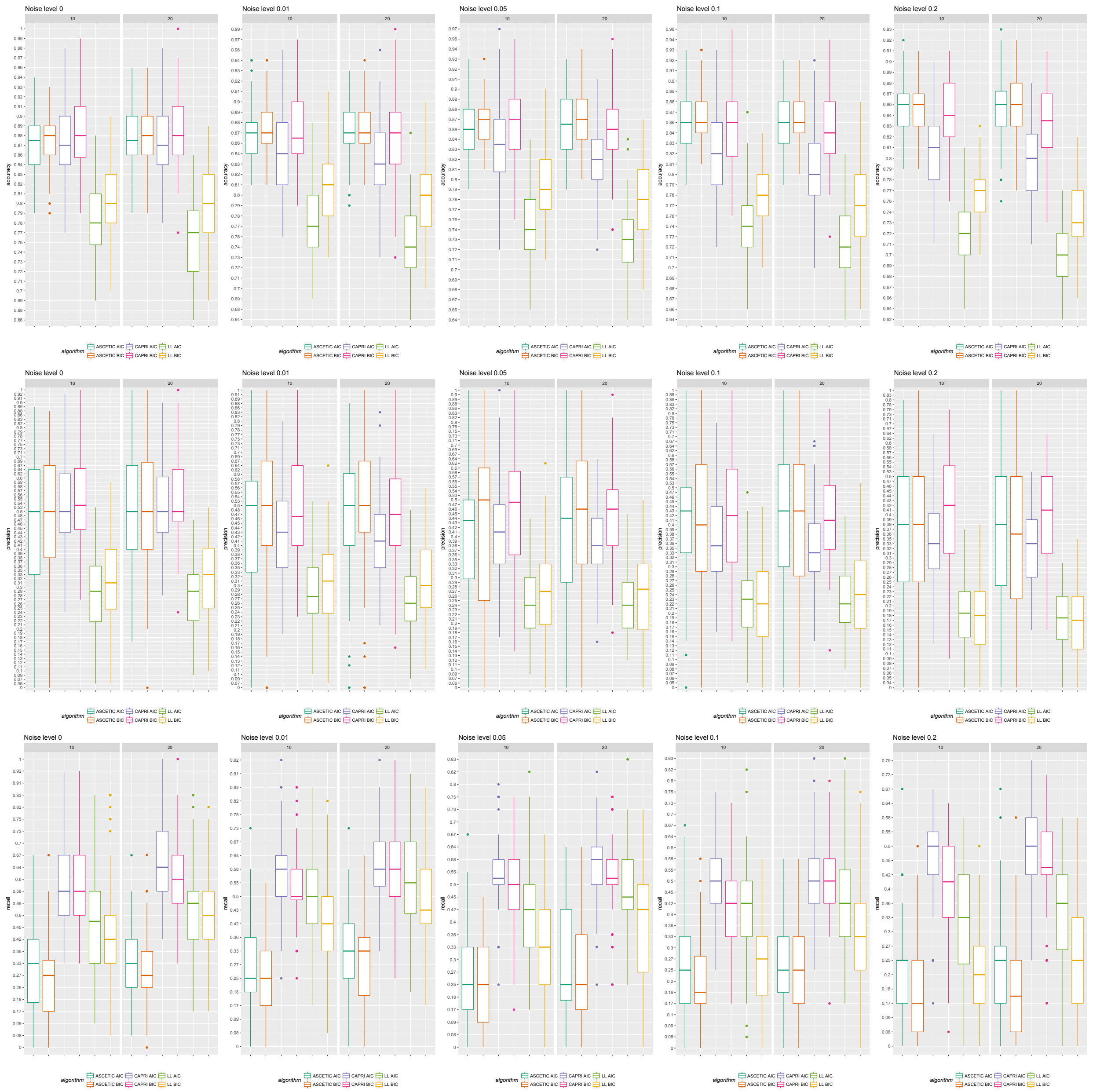

Figure 13: Results of the simulations for the agony method. Here we simulated NGS data from multiple biopses with 3 overlapping progressions of 5 mutations each for a total of 10 mutations. In each column we consider 5 noise levels, namely 0%, 1%, 5%, 10%, 20%, and datasets of 10 or 20 patients with 10 biopses each. In each row we show the results for different matrics, specifically for accuracy, precision and recall. ASCETIC is compared against CAPRI [15] and the standard likelihood fit with both AIC [17] and BIC [18] likelihood scores in terms of recall.

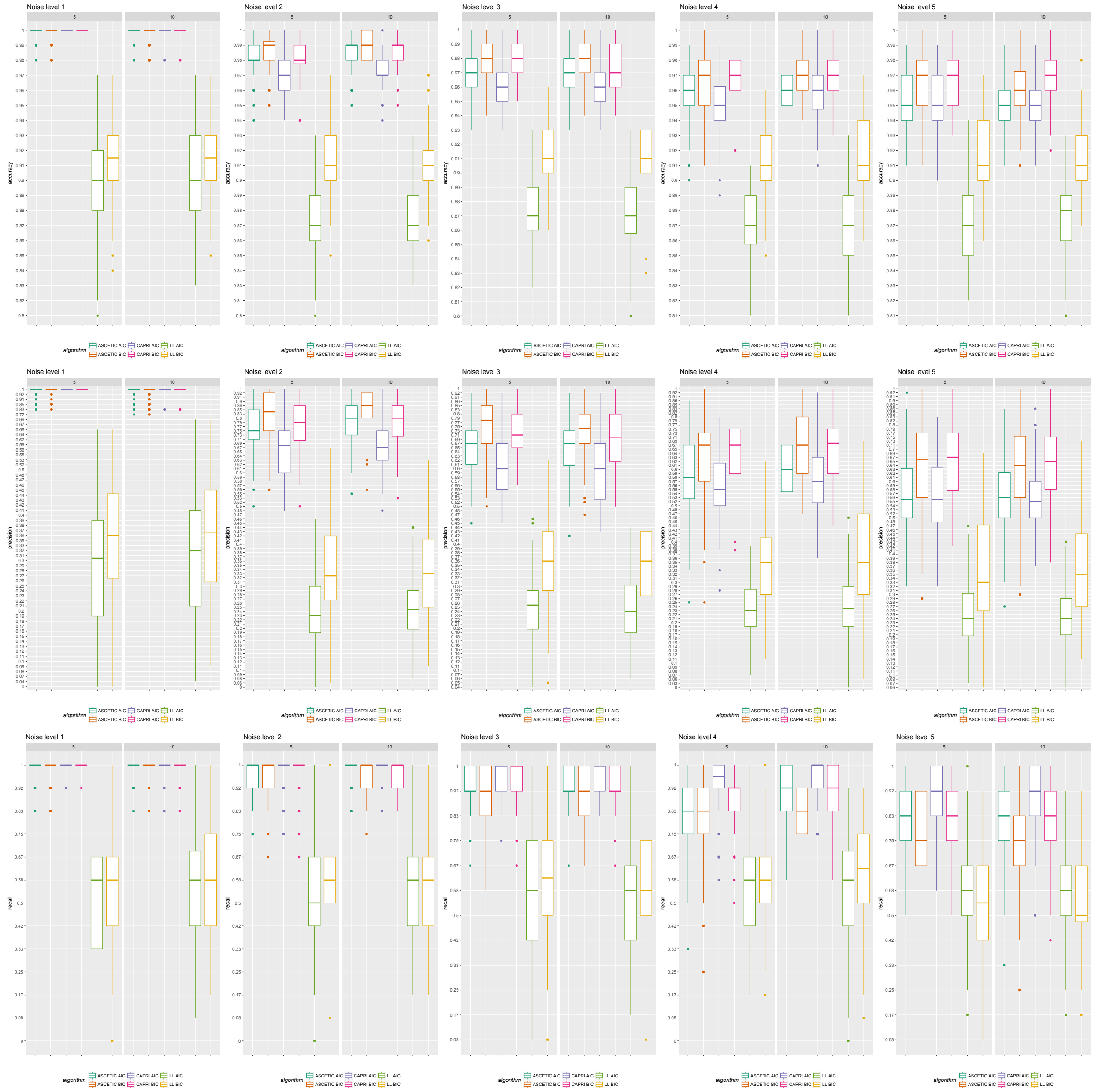

Figure 14: Results of the simulations for the agony method. Here we simulated data from single cell with 3 independent progressions and a total of 15 mutations. In each column we consider 5 unbalanced noise levels for false positive and false negative (allele dropout) errors respectively of 0%, 1%, 2%, 3% and 4% and 0%, 10%, 20%, 30% and 40%. Moreover, we considered datasets of 5 or 10 patients with 25 cells each. In each row we show the results for different matrics, specifically for accuracy, precision and recall. ASCETIC is compared against CAPRI [15] and the standard likelihood fit with both AIC [17] and BIC [18] likelihood scores in terms of recall.

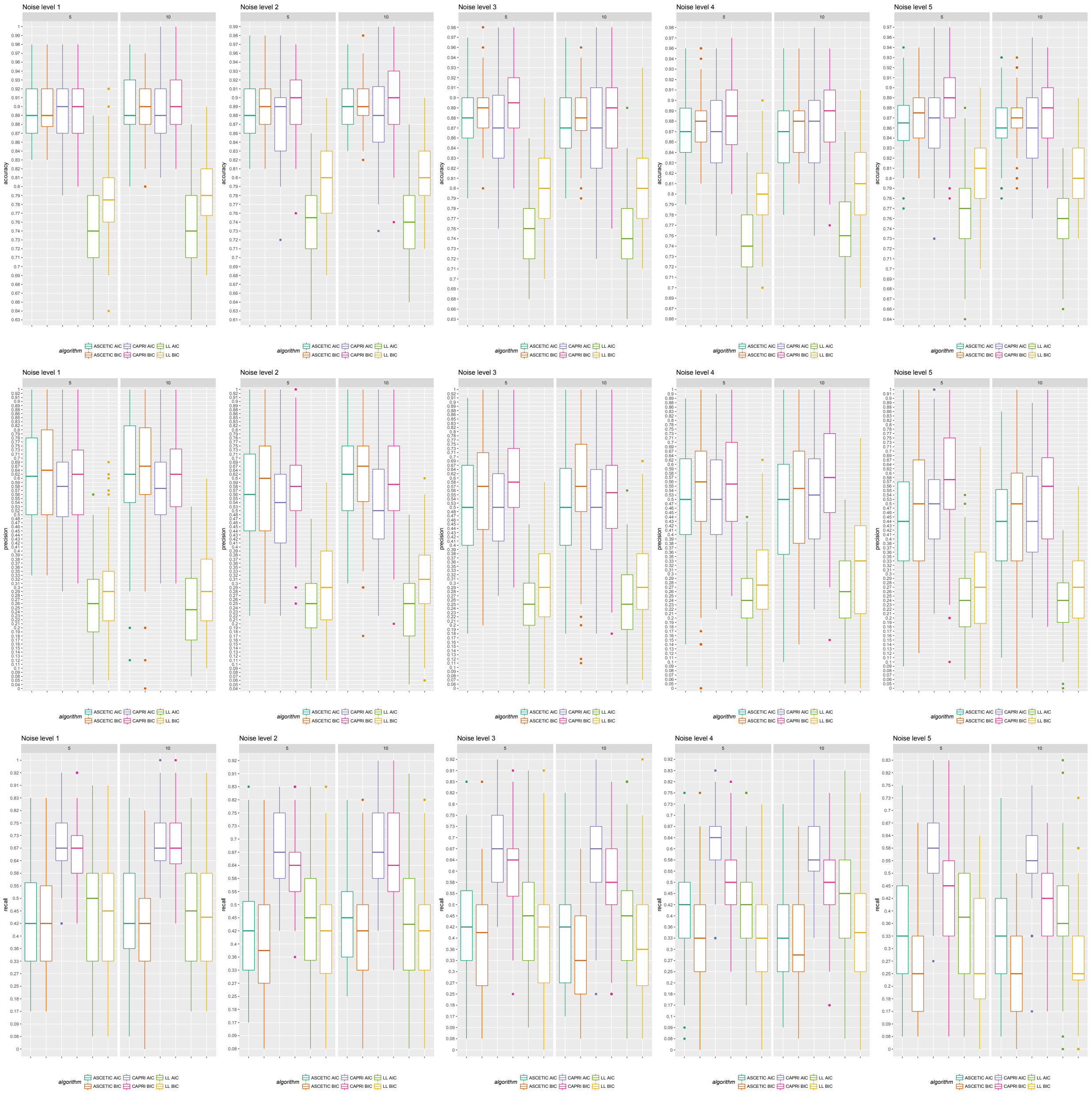

Figure 15: Results of the simulations for the agony method. Here we simulated data from single cell with 3 overlapping progressions and a total of 10 mutations. In each column we consider 5 unbalanced noise levels for false positive and false negative (allele dropout) errors respectively of 0%, 1%, 2%, 3% and 4% and 0%, 10%, 20%, 30% and 40%. Moreover, we considered datasets of 5 or 10 patients with 25 cells each. In each row we show the results for different matrics, specifically for accuracy, precision and recall. ASCETIC is compared against CAPRI [15] and the standard likelihood fit with both AIC [17] and BIC [18] likelihood scores in terms of recall.

### 4 Results on cancer data

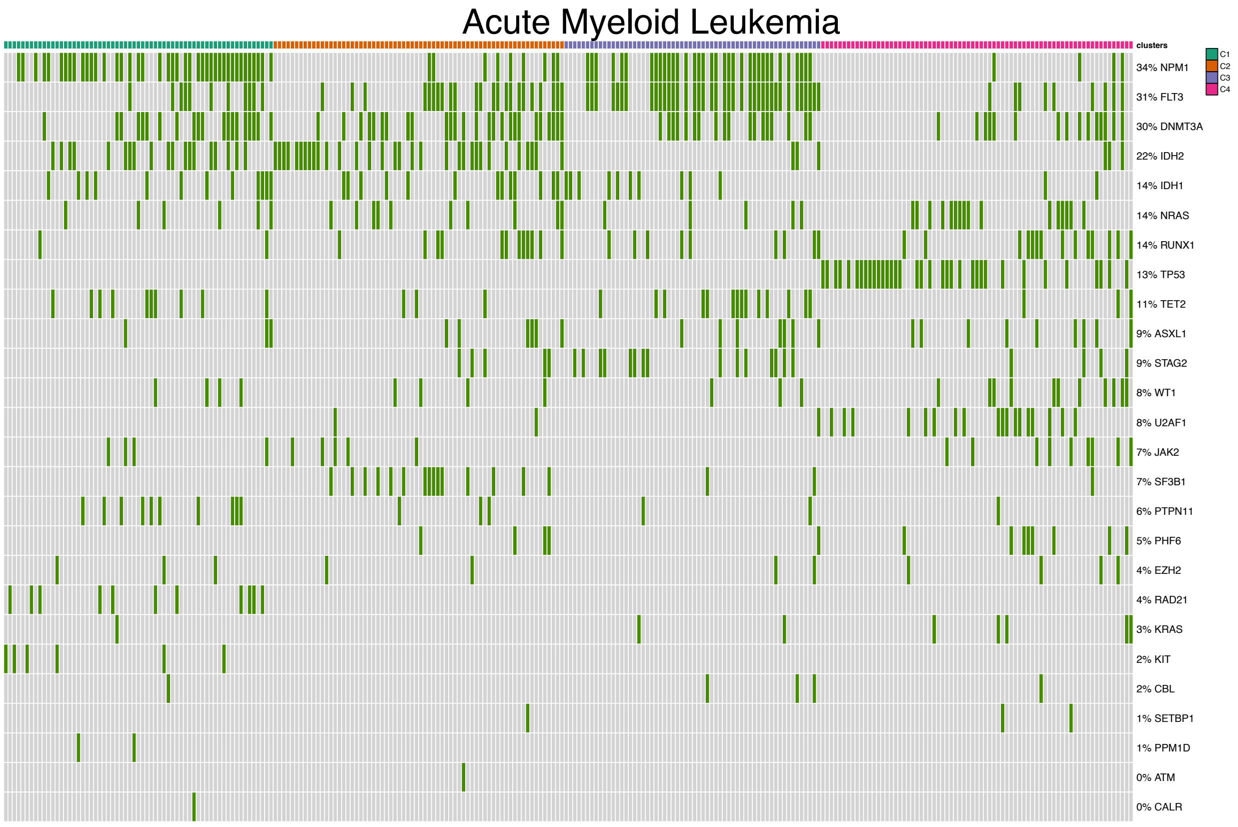

Figure 16: Mutational profile for Acute Myeloid Leukemia [19].

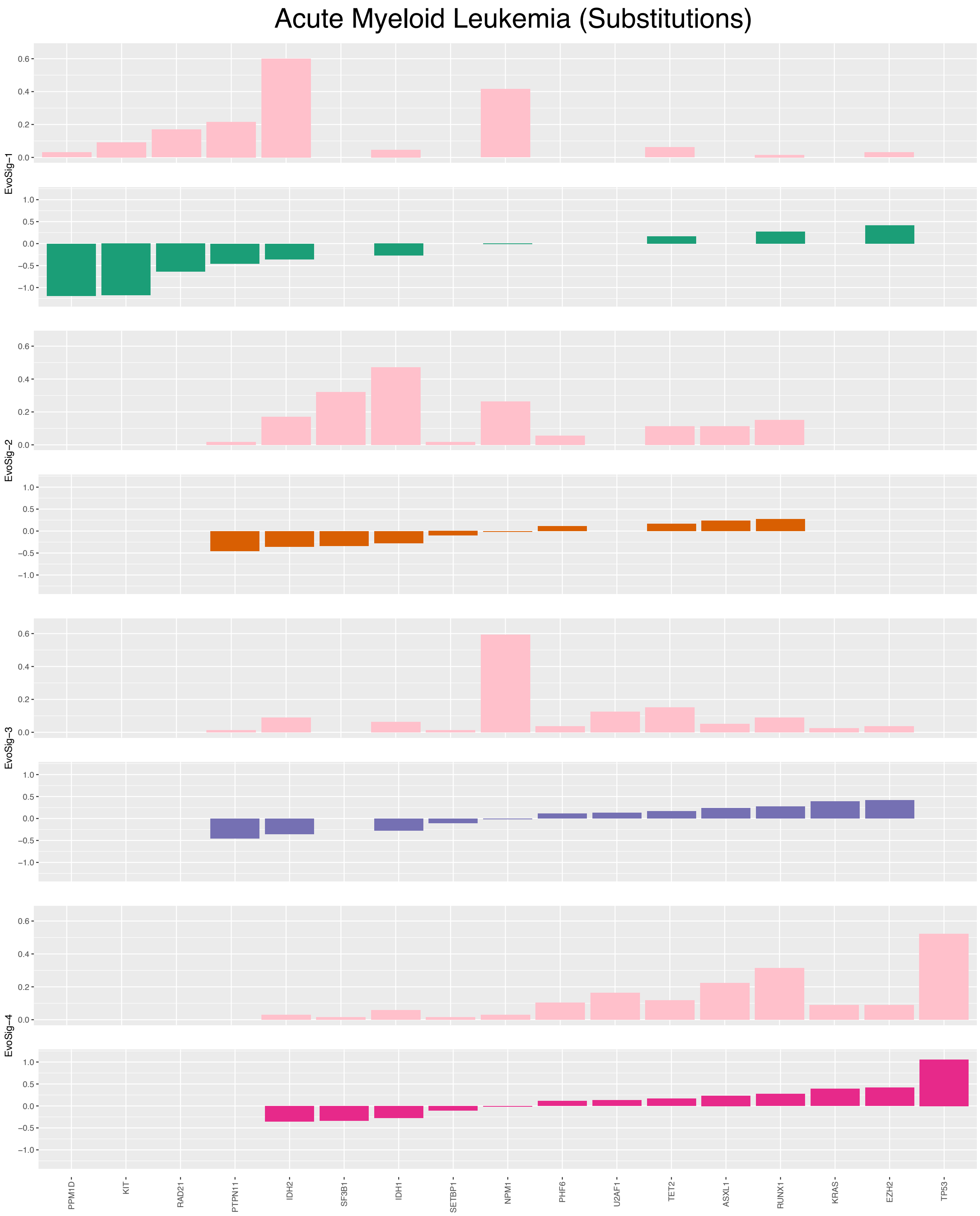

Figure 17: Association of substitutions to prognosis in Acute Myeloid Leukemia [19].

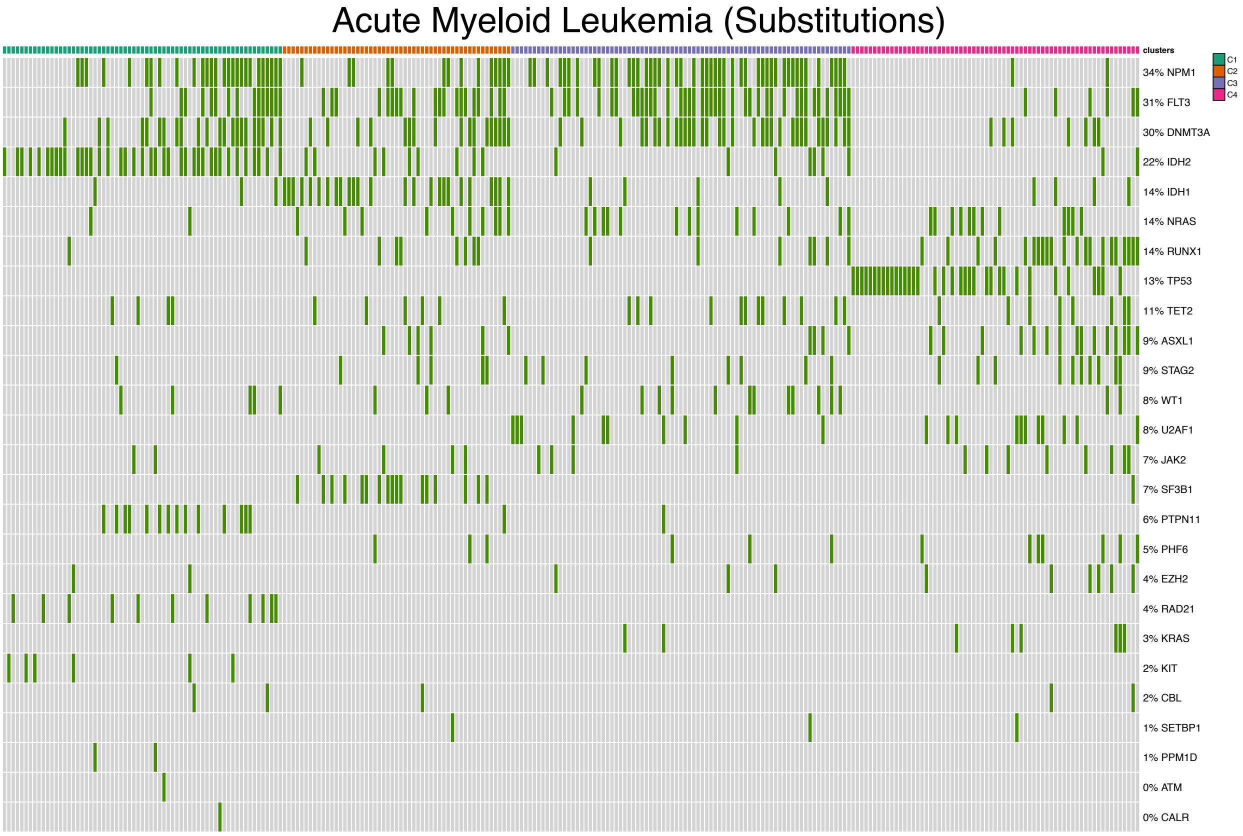

Figure 18: Mutational profile for Acute Myeloid Leukemia [19] considering substitutions.

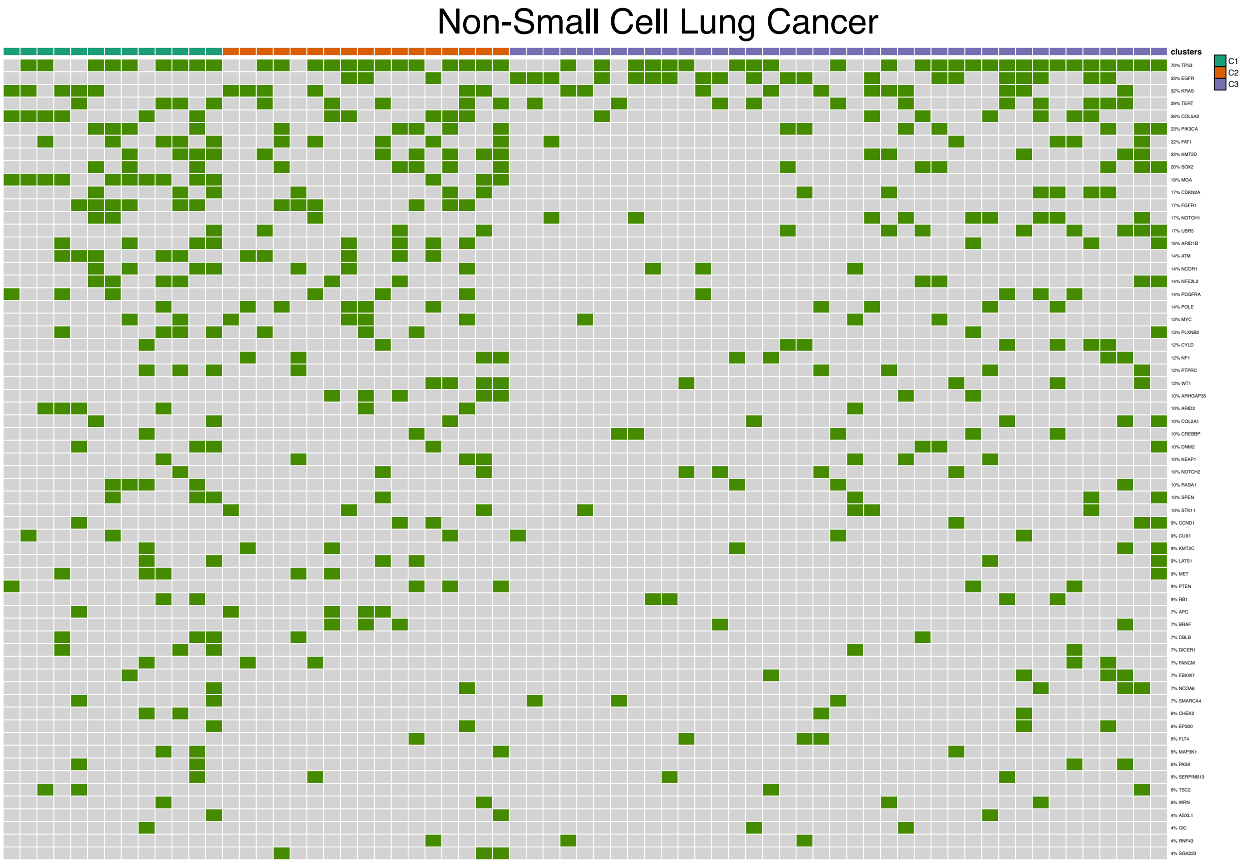

Figure 19: Oncoprint for Early-stage Non-Small Lung Cancer (TRACERx) [20].

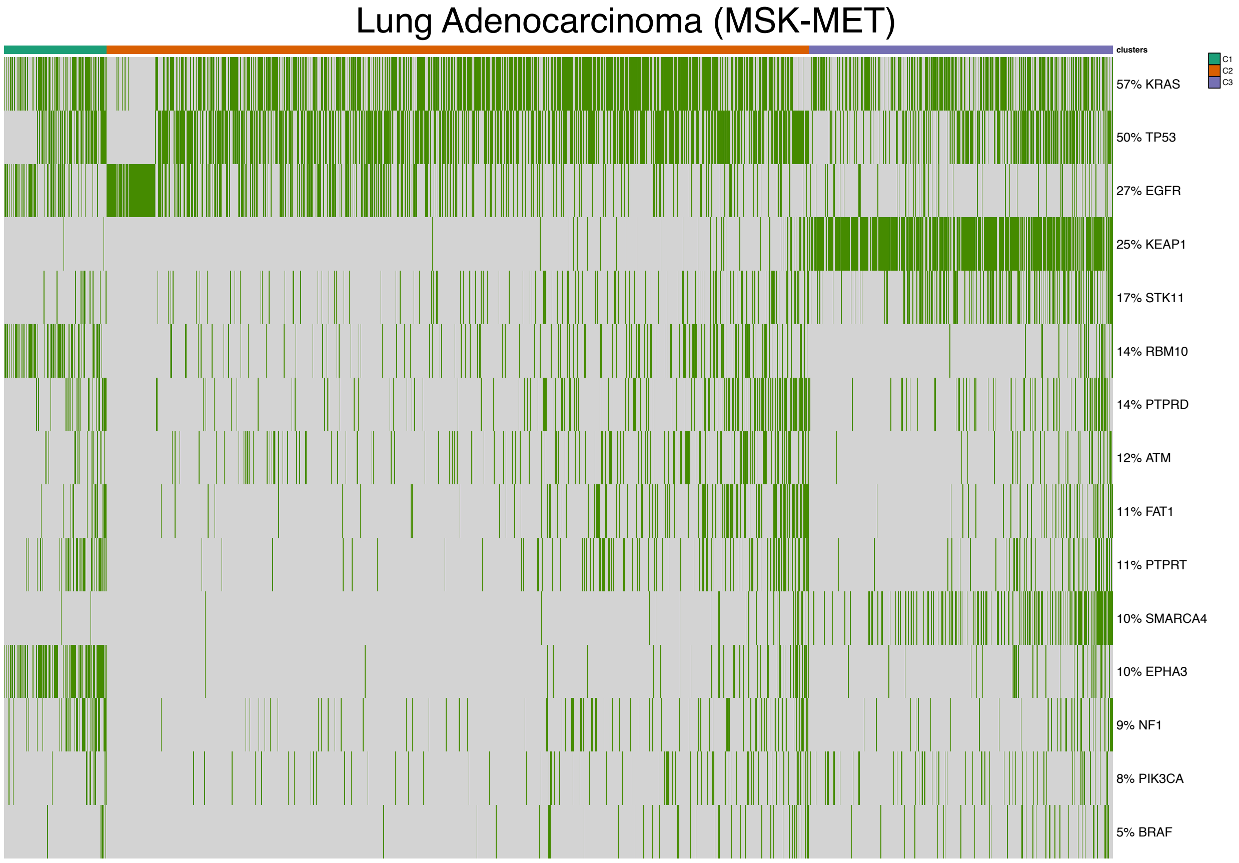

Figure 20: Oncoprint for Lung Adenocarcinoma (MSK-MET) [21].

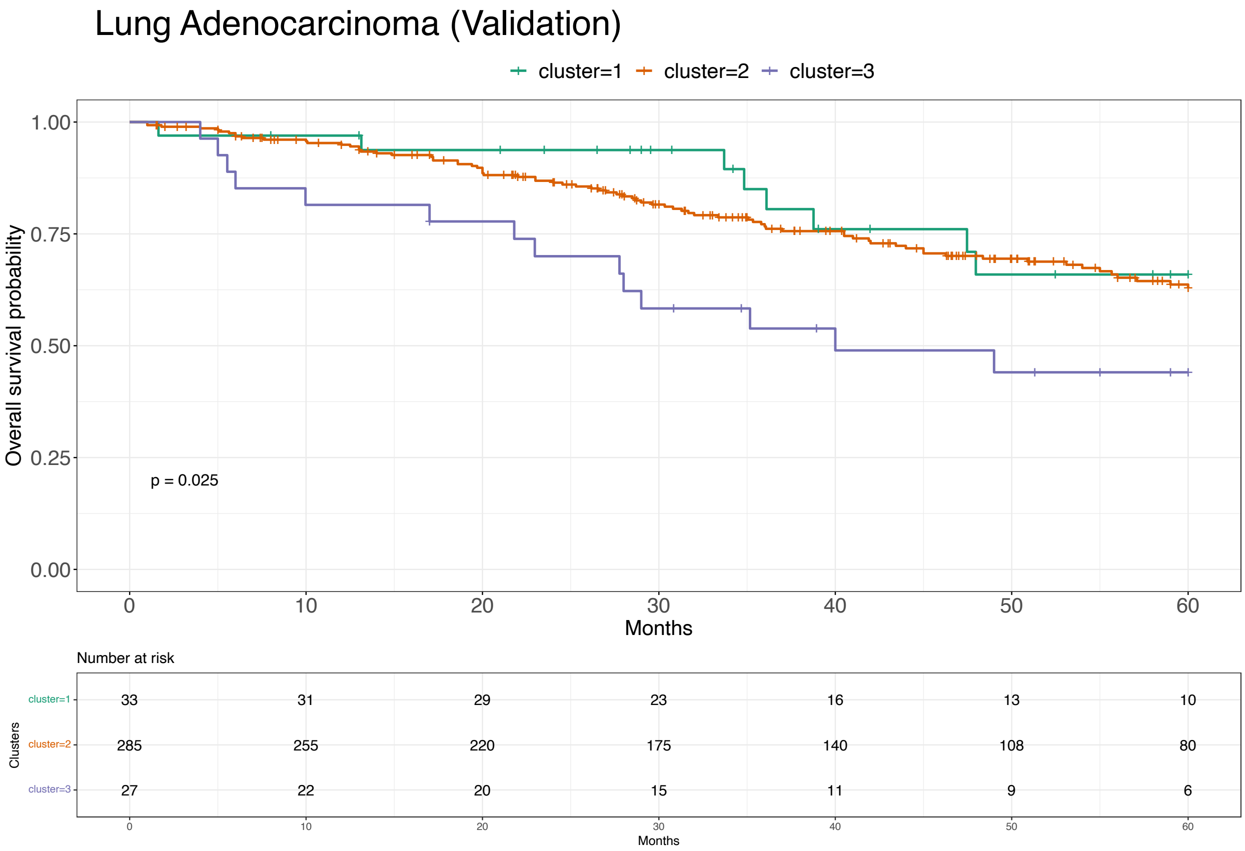

Figure 21: Survival analysis for Lung Adenocarcinoma (Validation) [22].

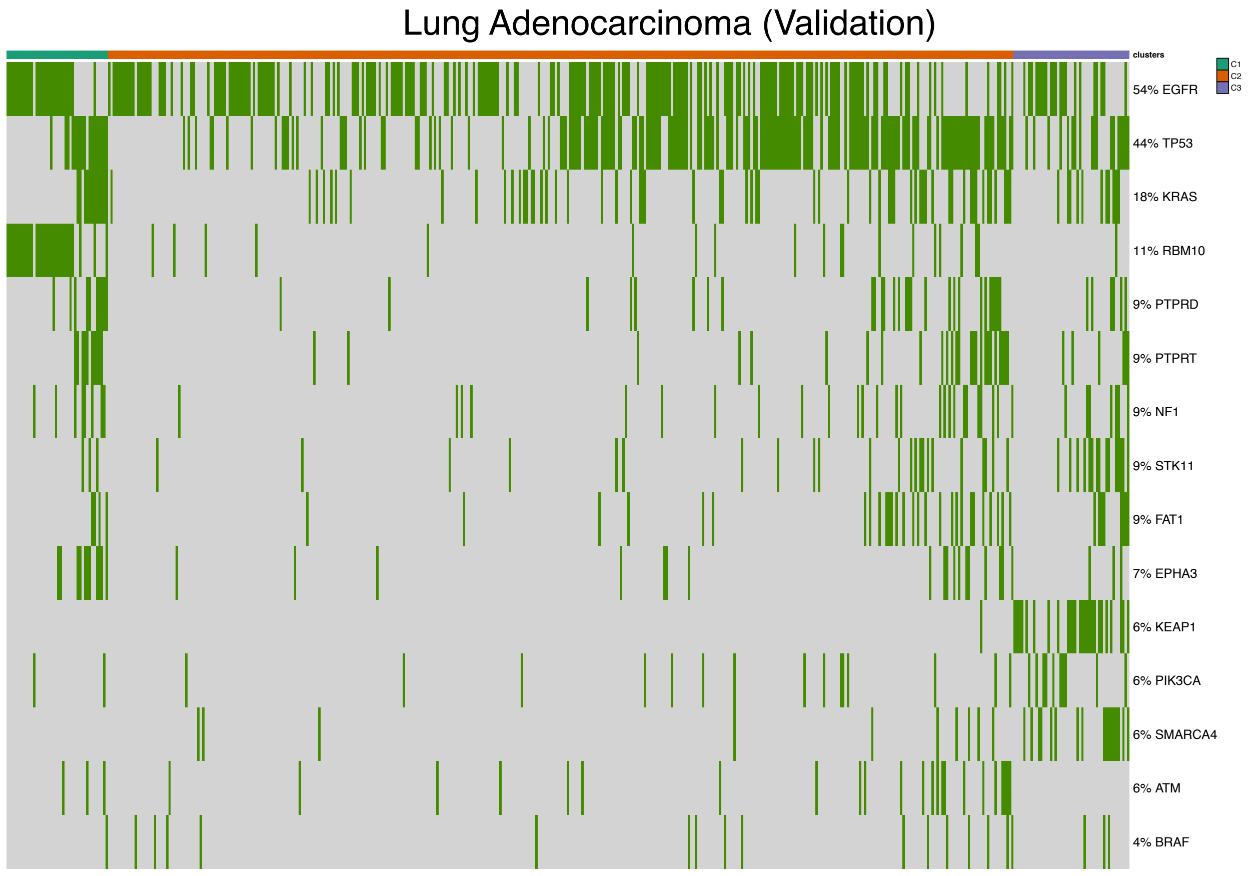

Figure 22: Mutational profile for Lung Adenocarcinoma (Validation) [22].

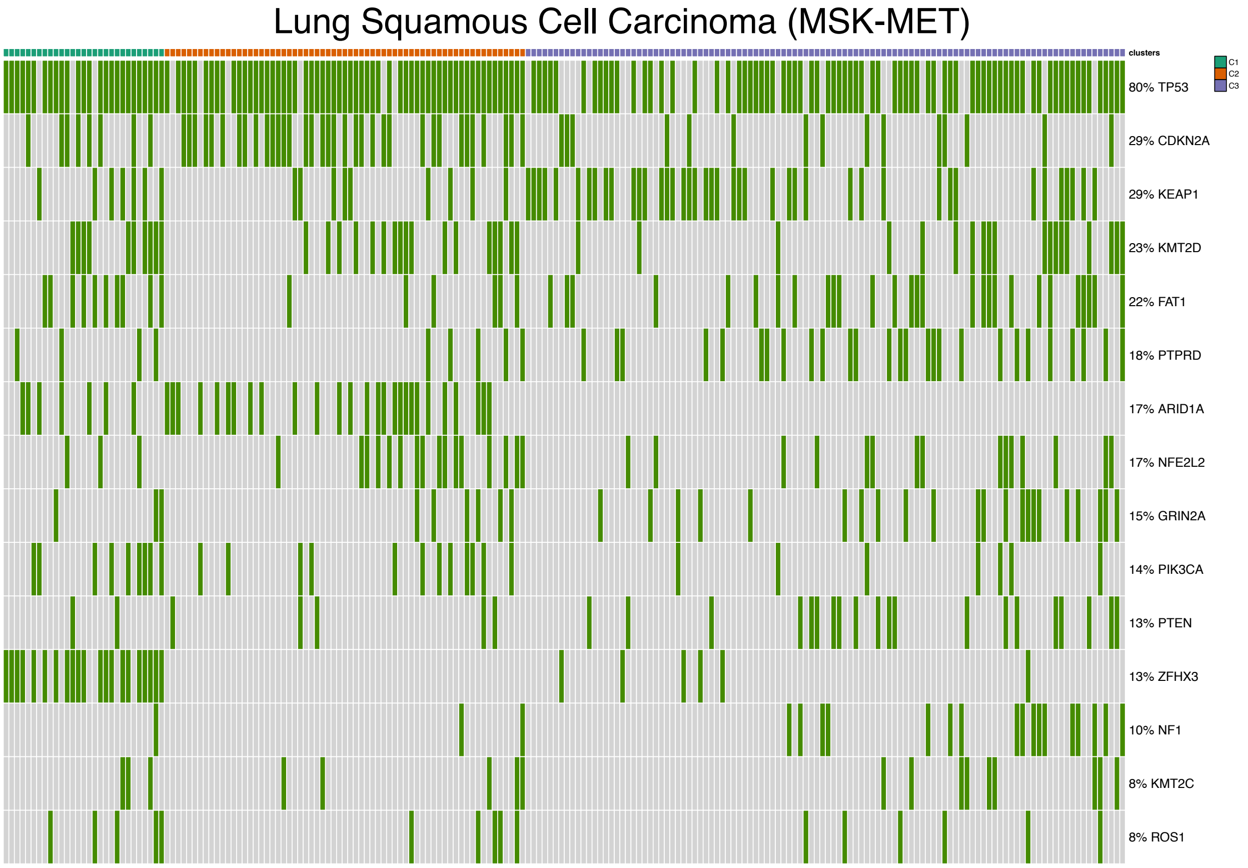

Figure 23: Oncoprint for Lung Squamous Cell Carcinoma (MSK-MET) [21].

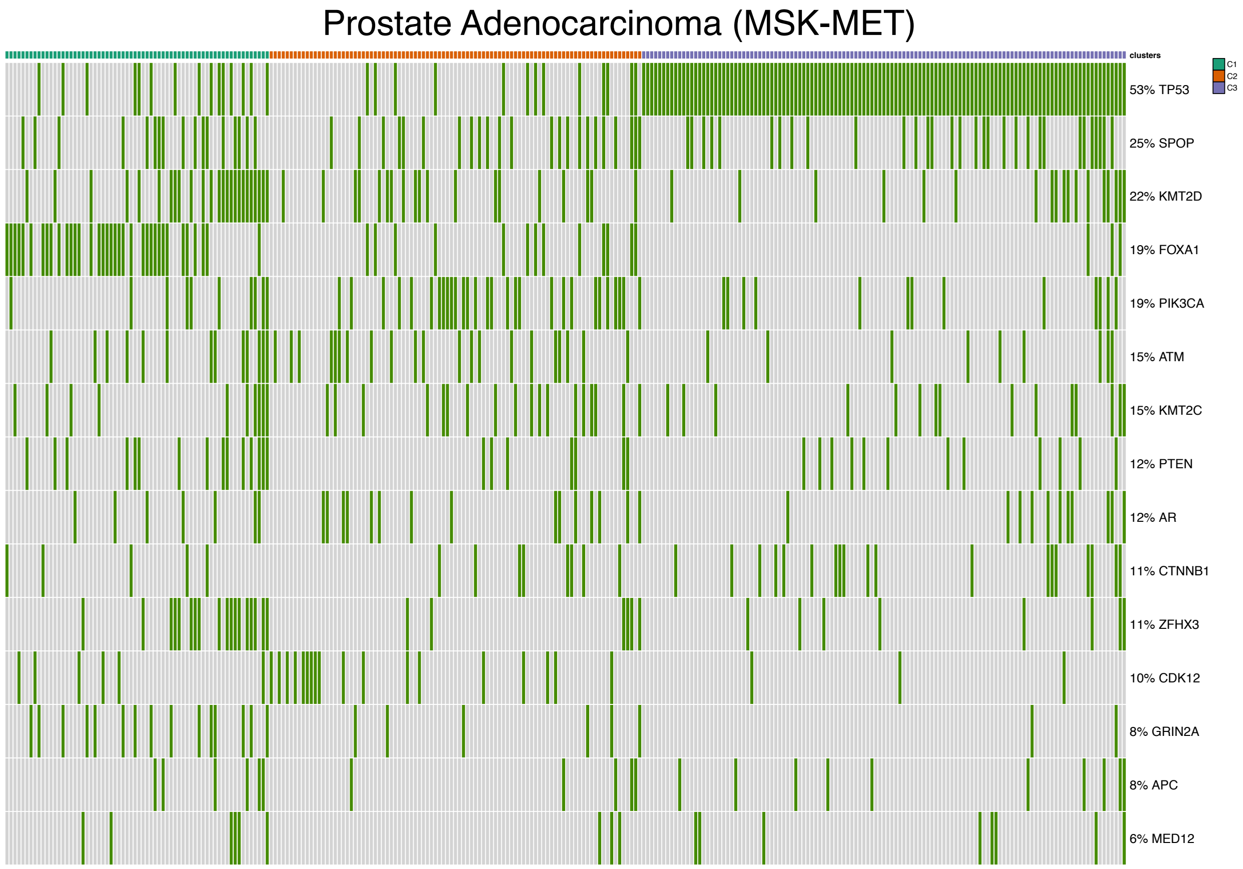

Figure 24: Oncoprint for Prostate Adenocarcinoma (MSK-MET) [21].

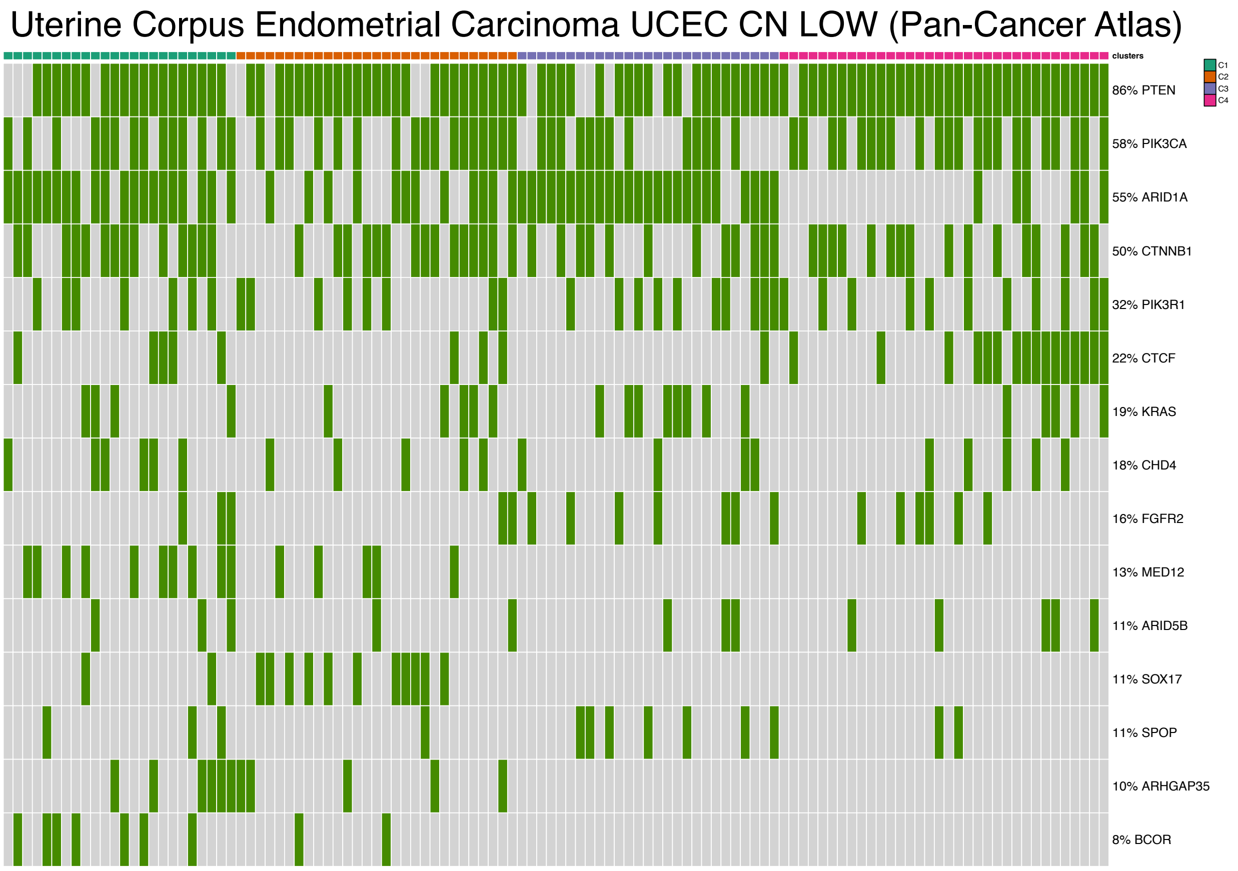

Figure 25: Oncoprint for Uterine Corpus Endometrial Carcinoma (UCEC CN LOW, Pan-Cancer Atlas) [23].

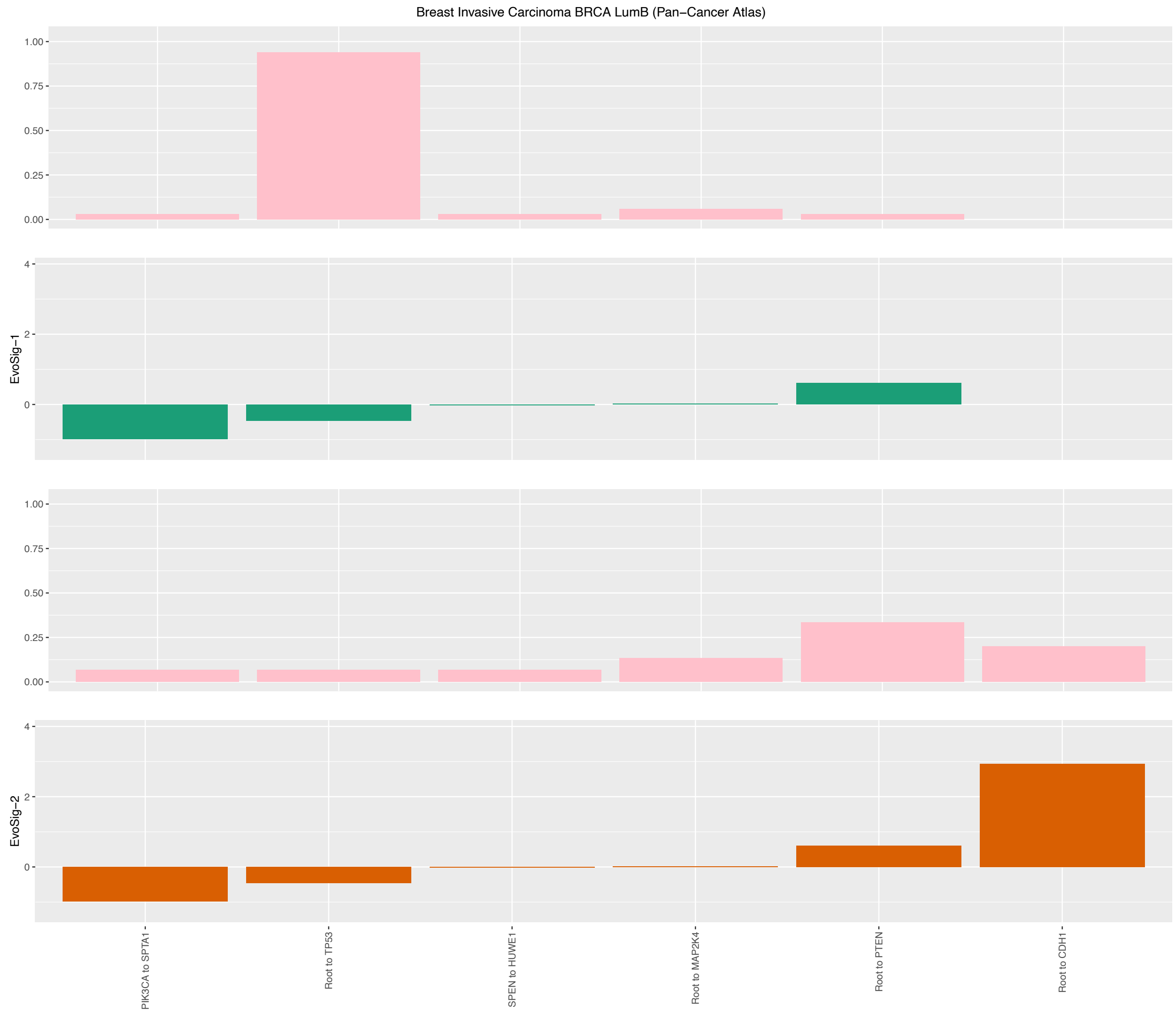

Figure 26: Evolutionary Signatures for Breast Invasive Carcinoma (Luminal-B, Pan-Cancer Atlas) [23].

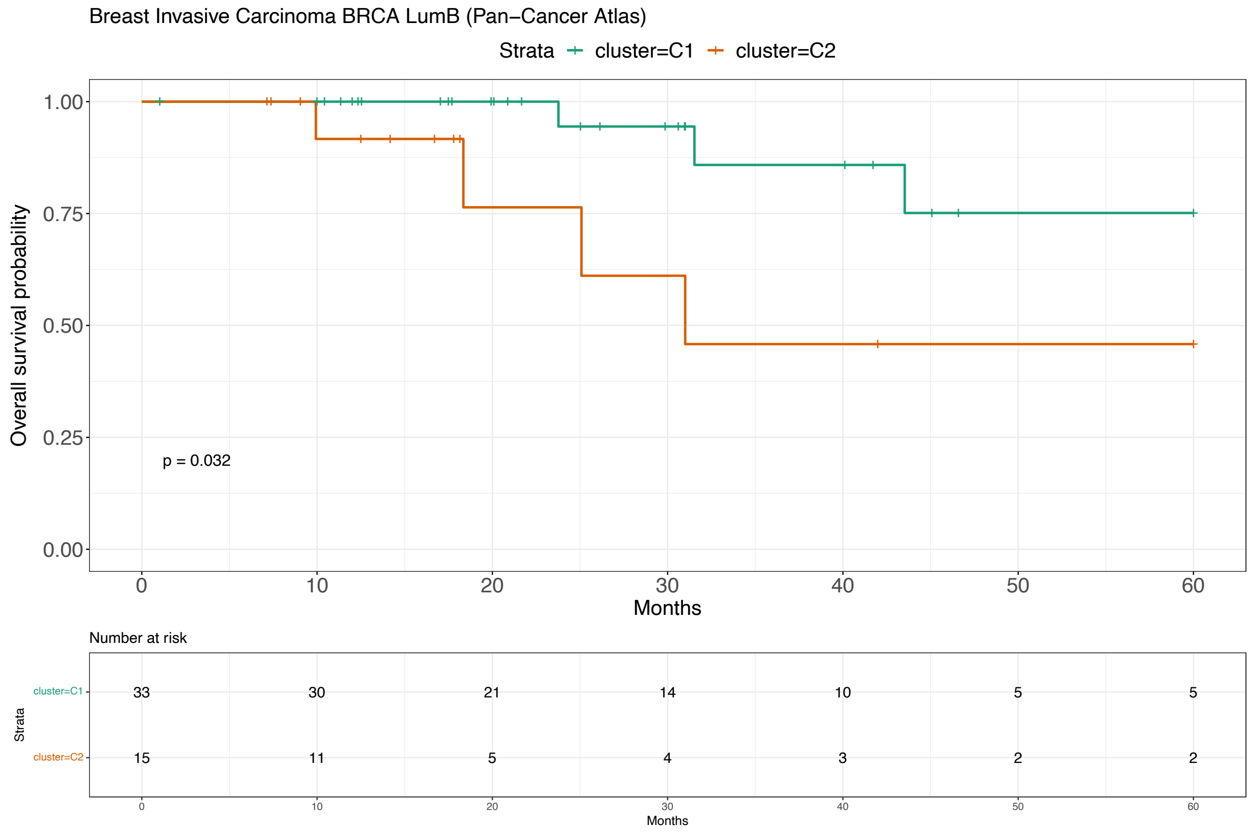

Figure 27: Survival analysis for Breast Invasive Carcinoma (Luminal-B, Pan-Cancer Atlas) [23].

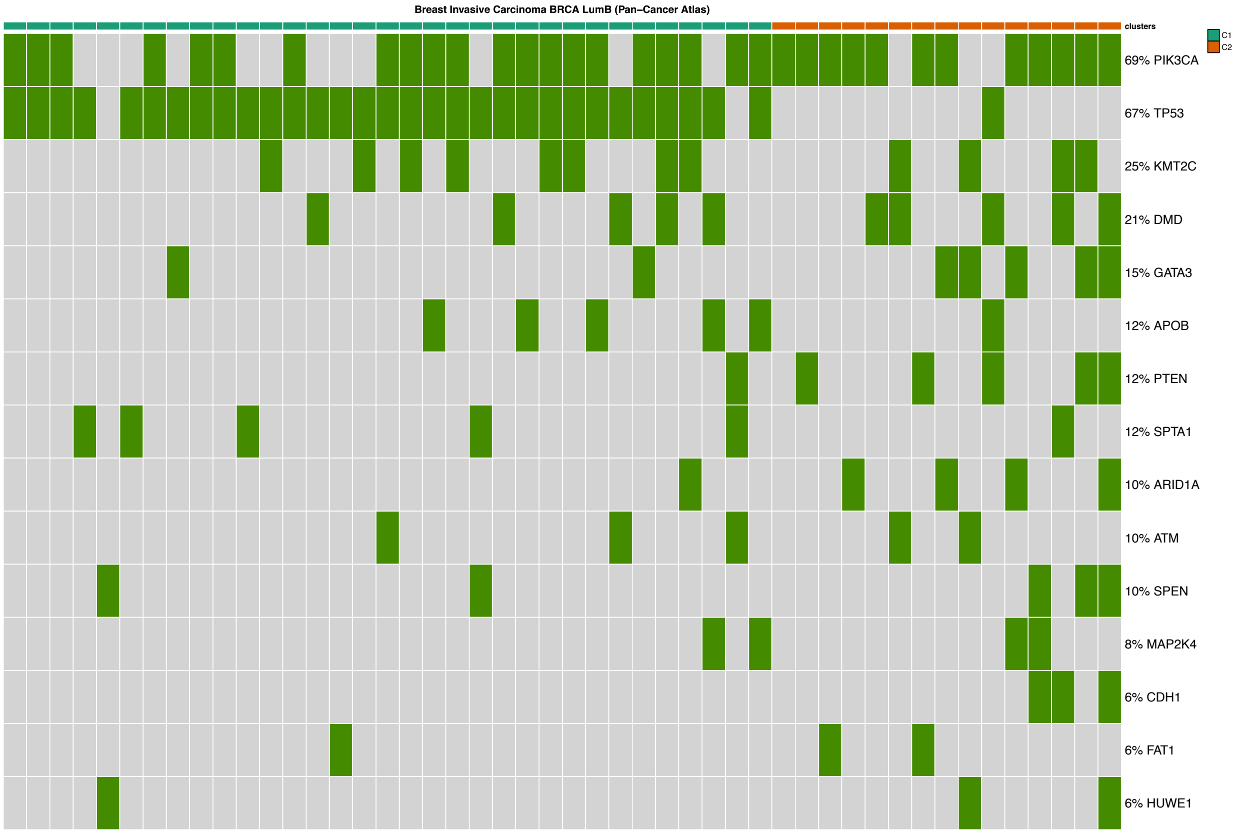

Figure 28: Mutational profile for Breast Invasive Carcinoma (Luminal-B, Pan-Cancer Atlas) [23].

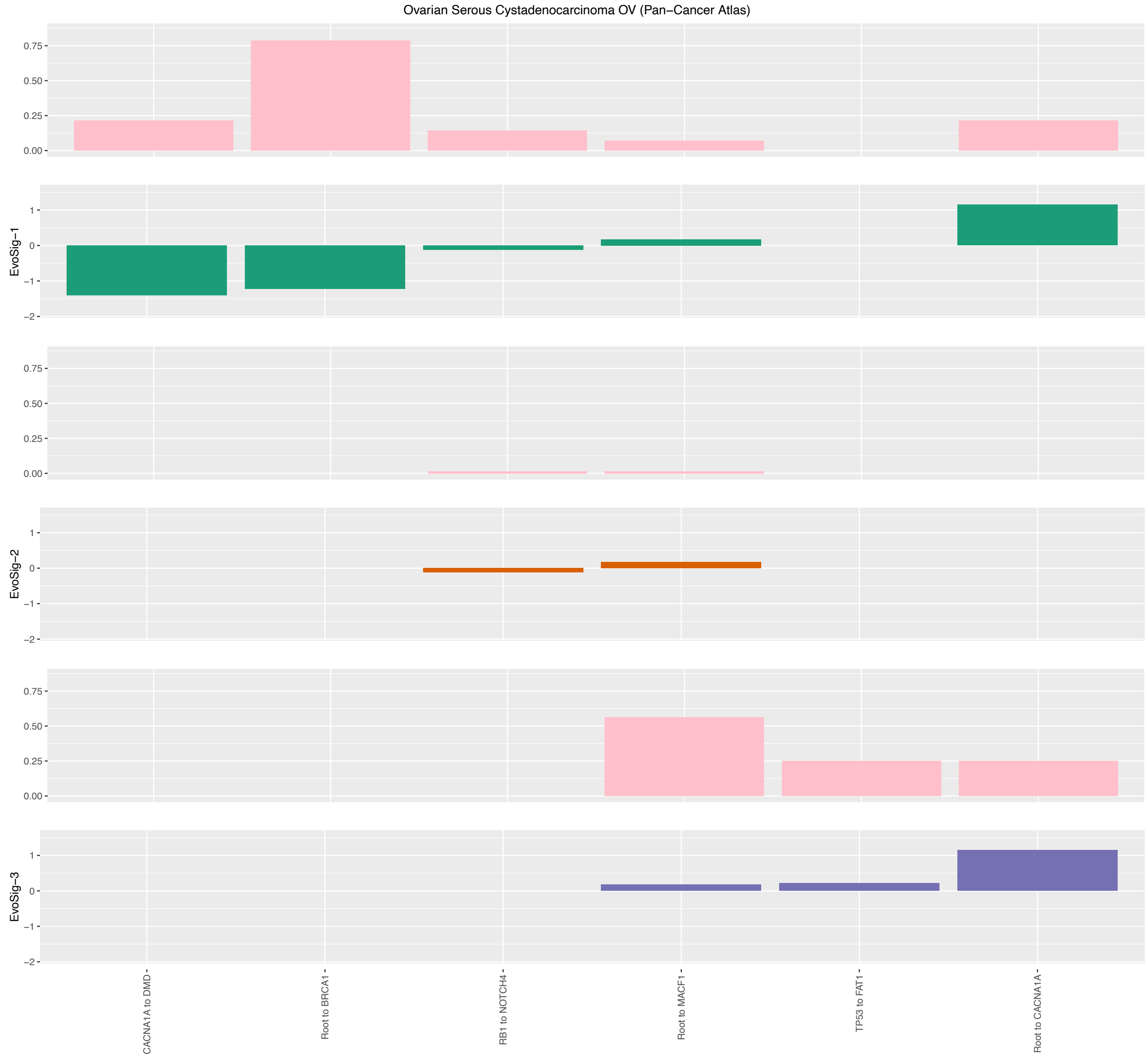

Figure 29: Evolutionary Signatures for Ovarian Serous Cystadenocarcinoma (Pan-Cancer Atlas) [23].

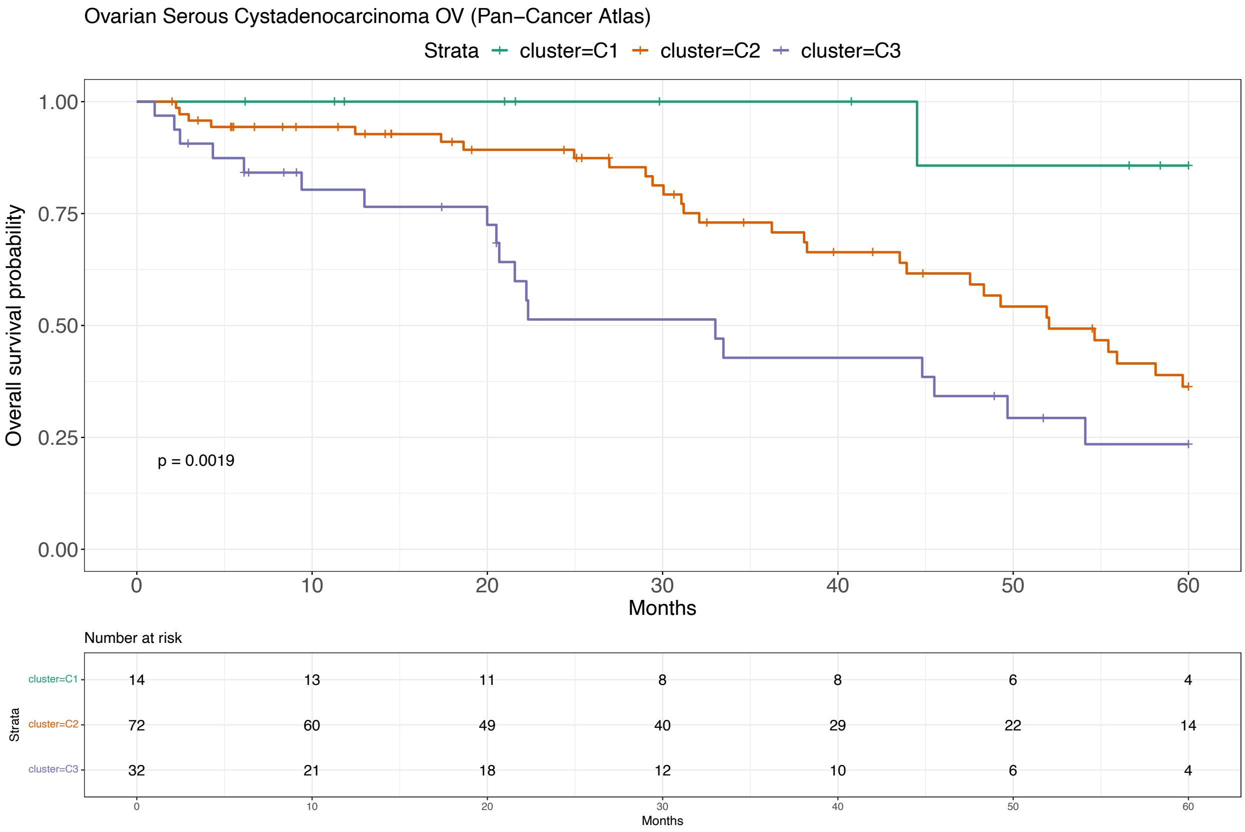

Figure 30: Survival analysis for Ovarian Serous Cystadenocarcinoma (Pan-Cancer Atlas) [23].

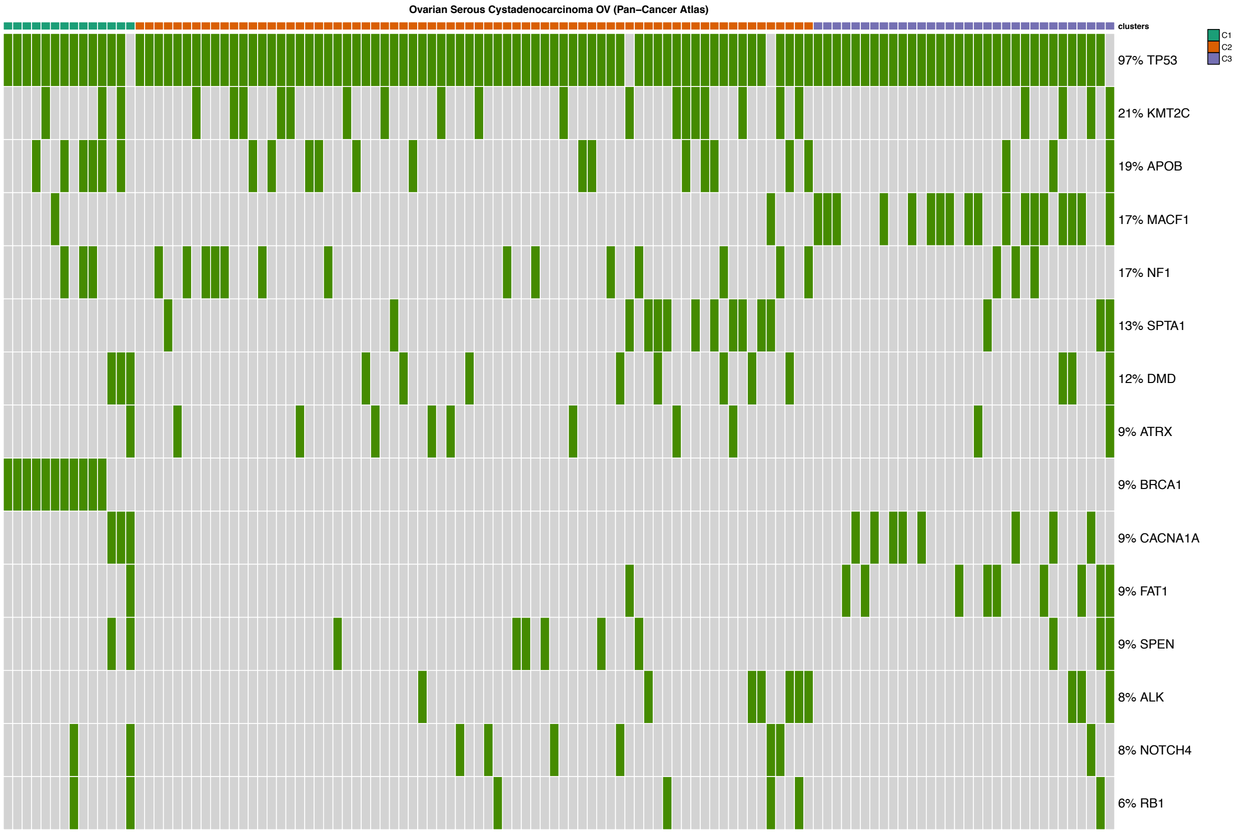

Figure 31: Mutational profile for Ovarian Serous Cystadenocarcinoma (Pan-Cancer Atlas) [23].

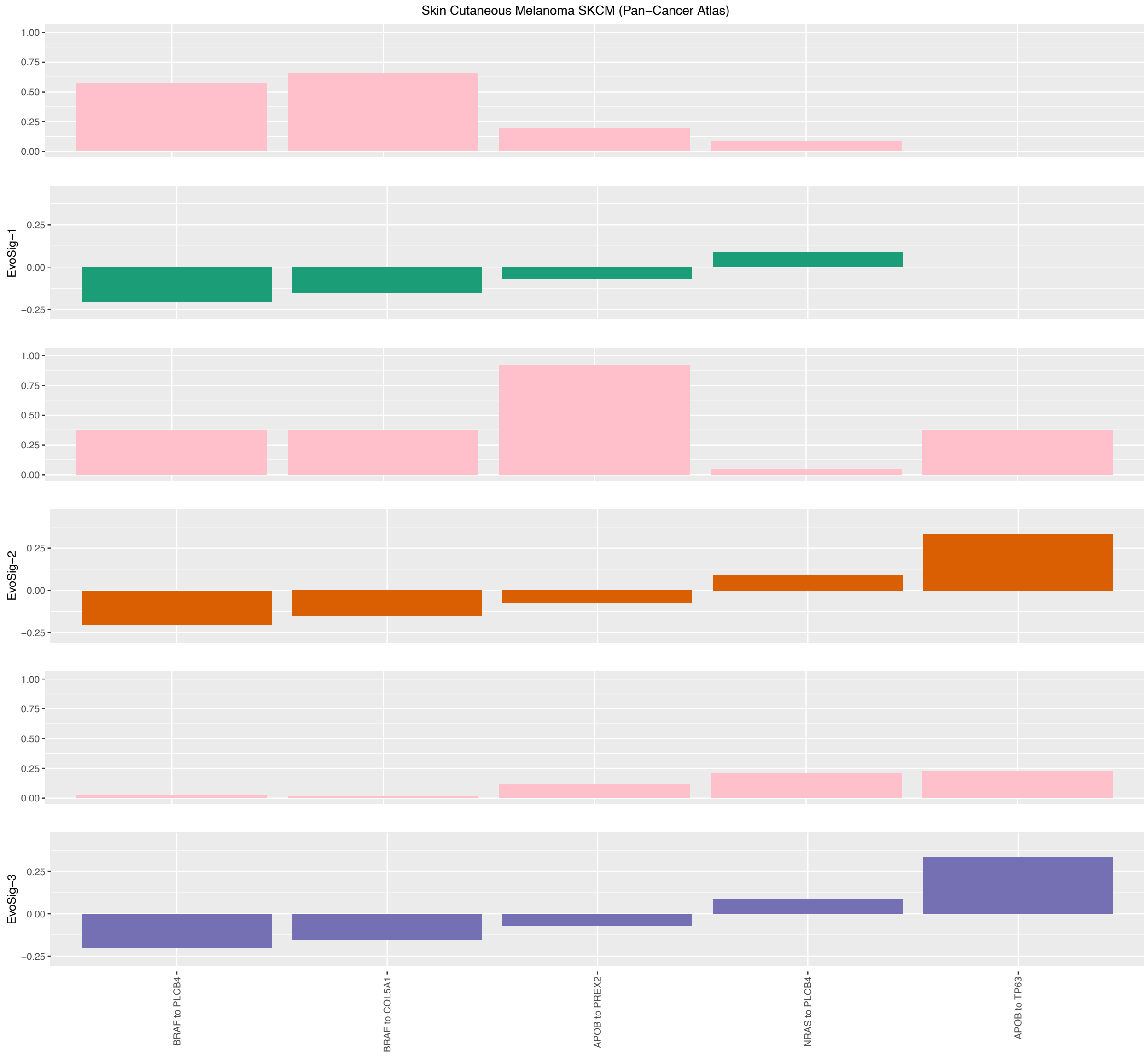

Figure 32: Evolutionary Signatures for Skin Cutaneous Melanoma (Pan-Cancer Atlas) [23].

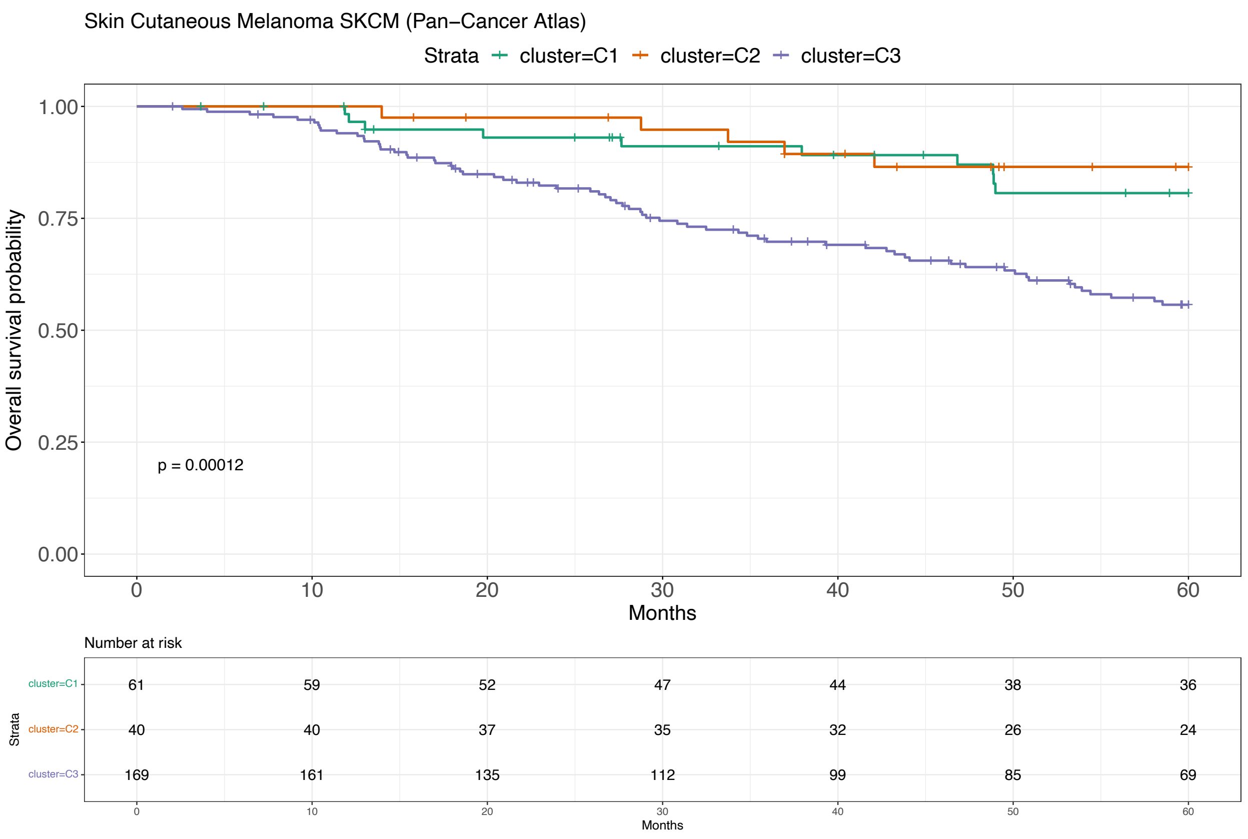

Figure 33: Survival analysis for Skin Cutaneous Melanoma (Pan-Cancer Atlas) [23].

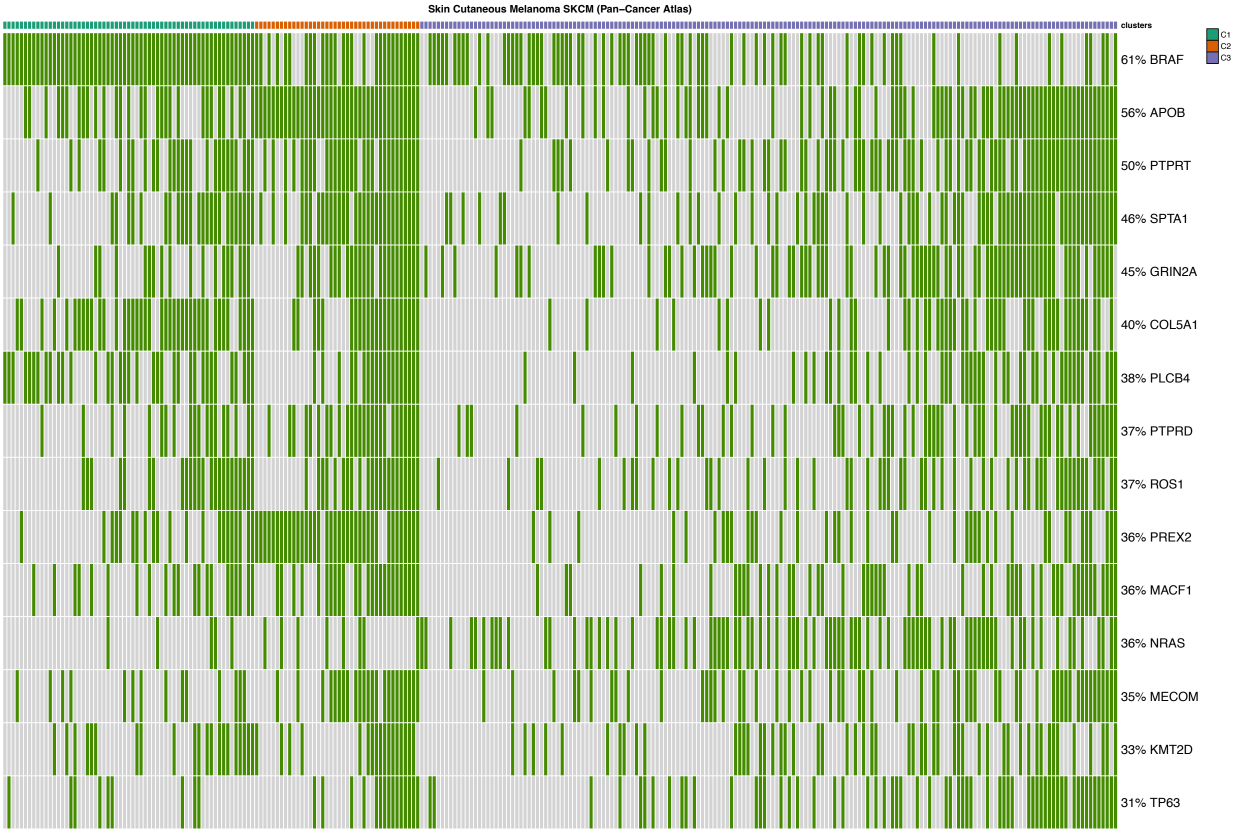

Figure 34: Mutational profile for Skin Cutaneous Melanoma (Pan-Cancer Atlas) [23].

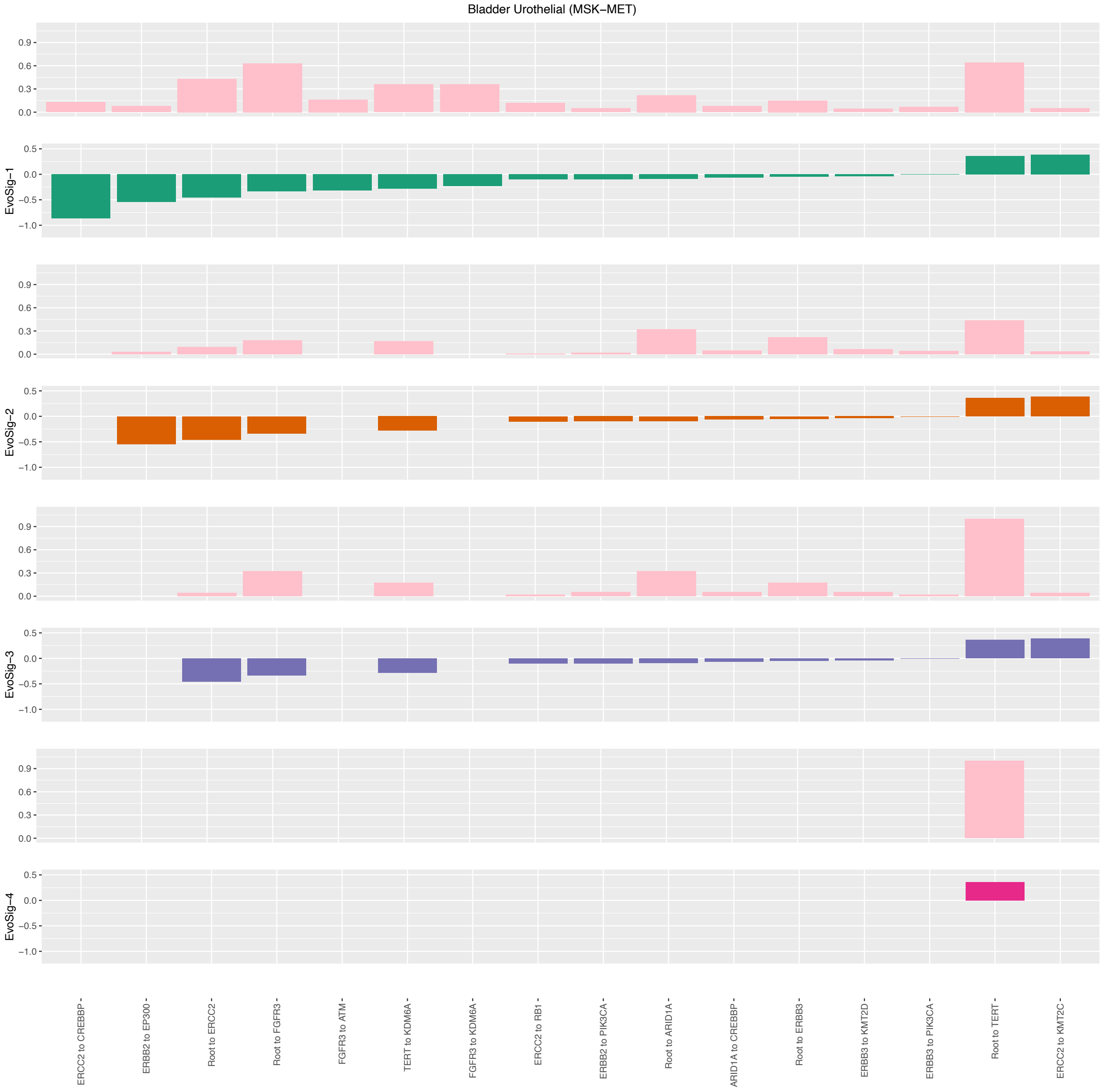

Figure 35: Evolutionary Signatures for Bladder Urothelial (MSK-MET) [21].

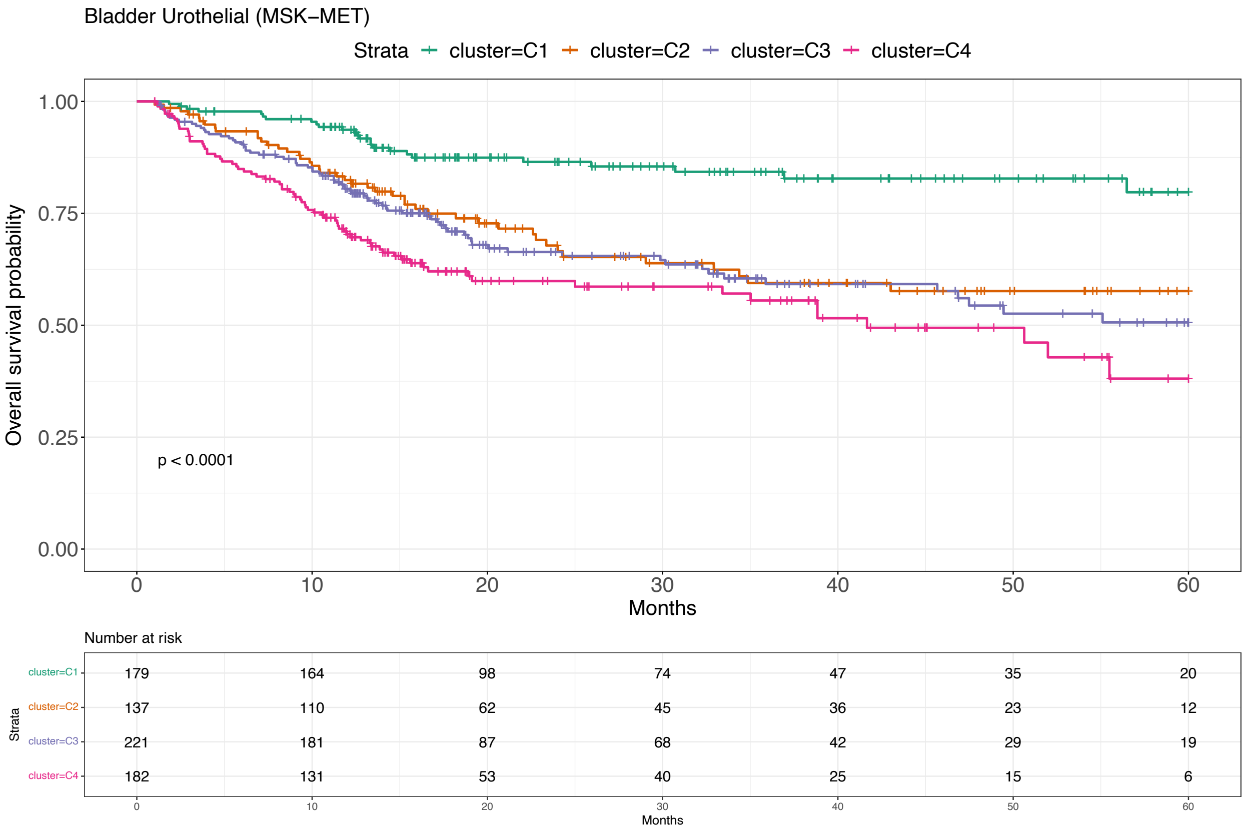

Figure 36: Survival analysis for Bladder Urothelial (MSK-MET) [21].

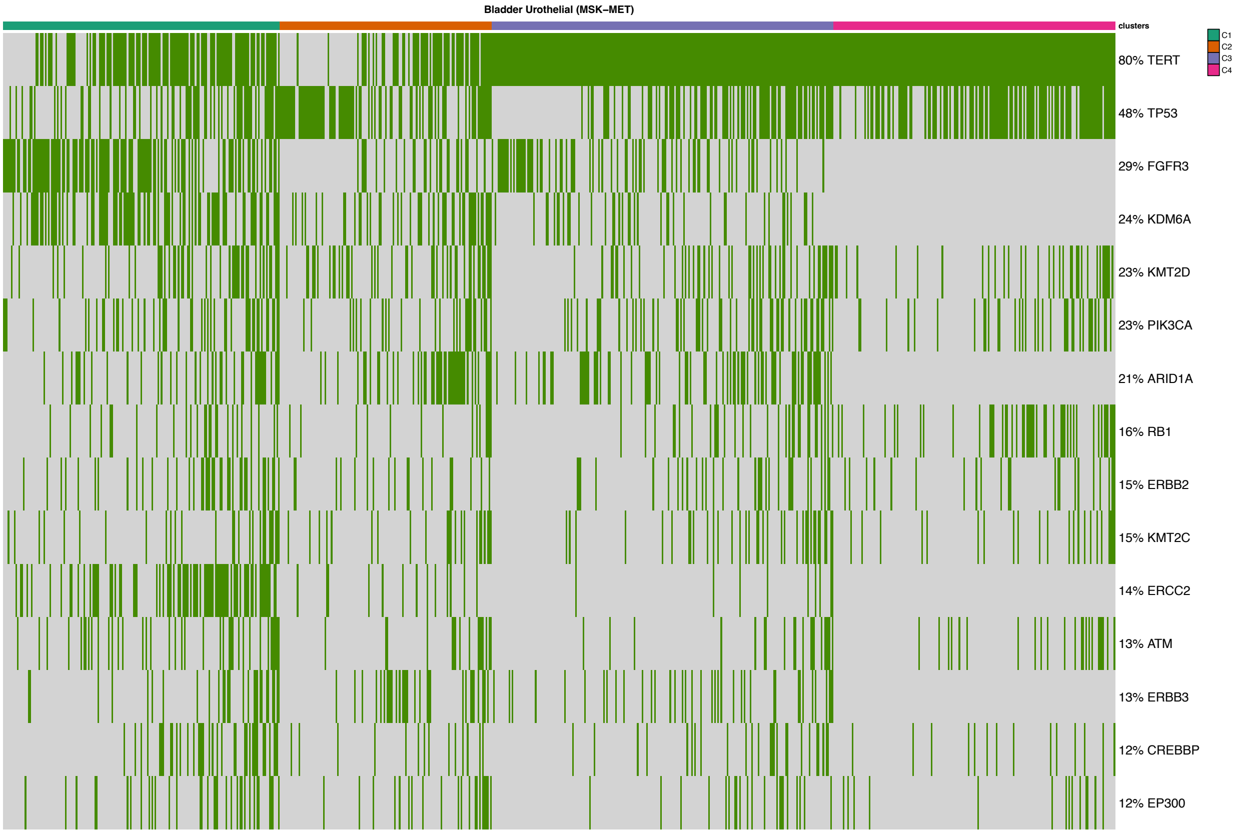

Figure 37: Mutational profile for Bladder Urothelial (MSK-MET) [21].

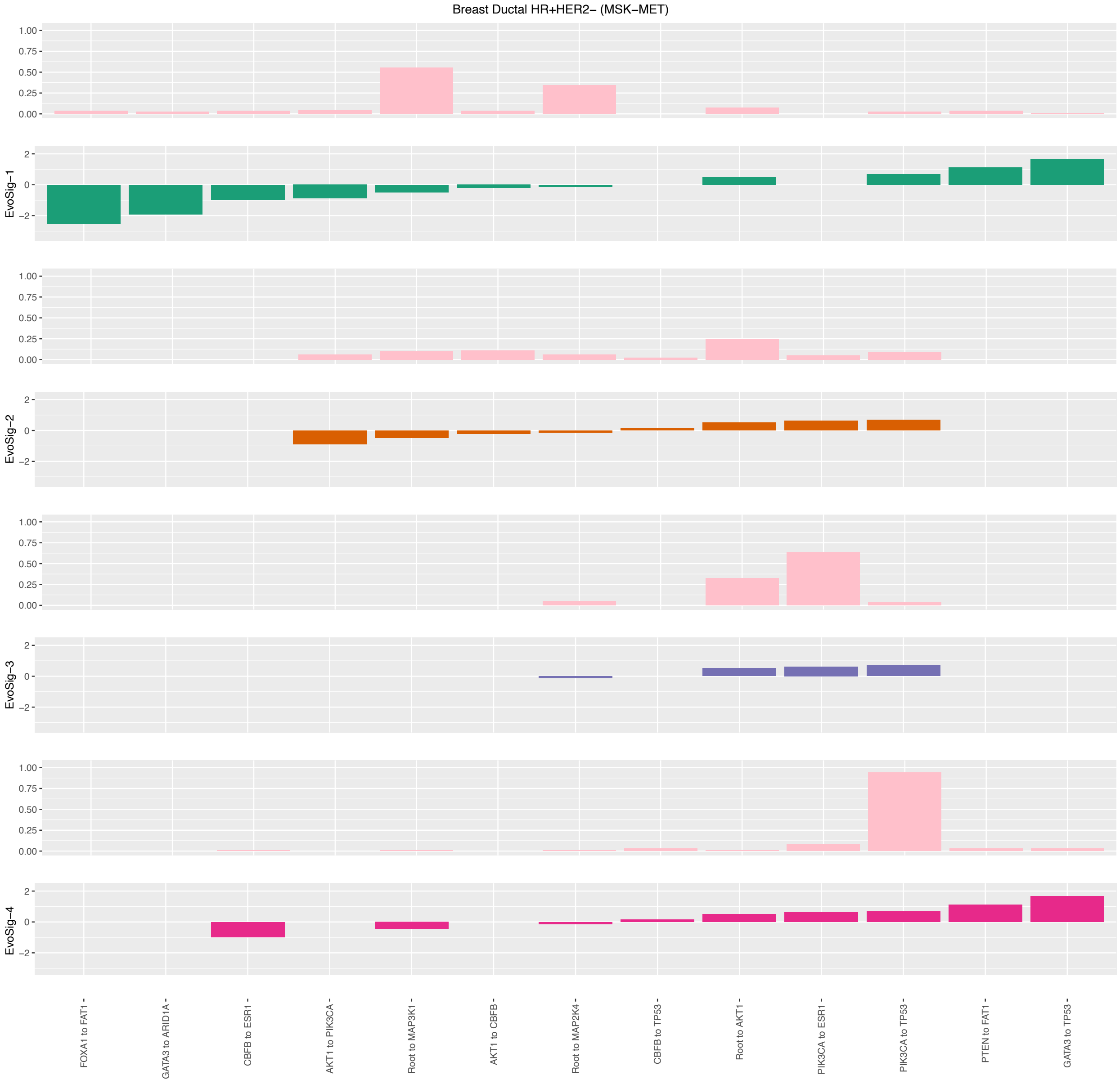

Figure 38: Evolutionary Signatures for Breast Ductal (HR+HER2-, MSK-MET) [21].

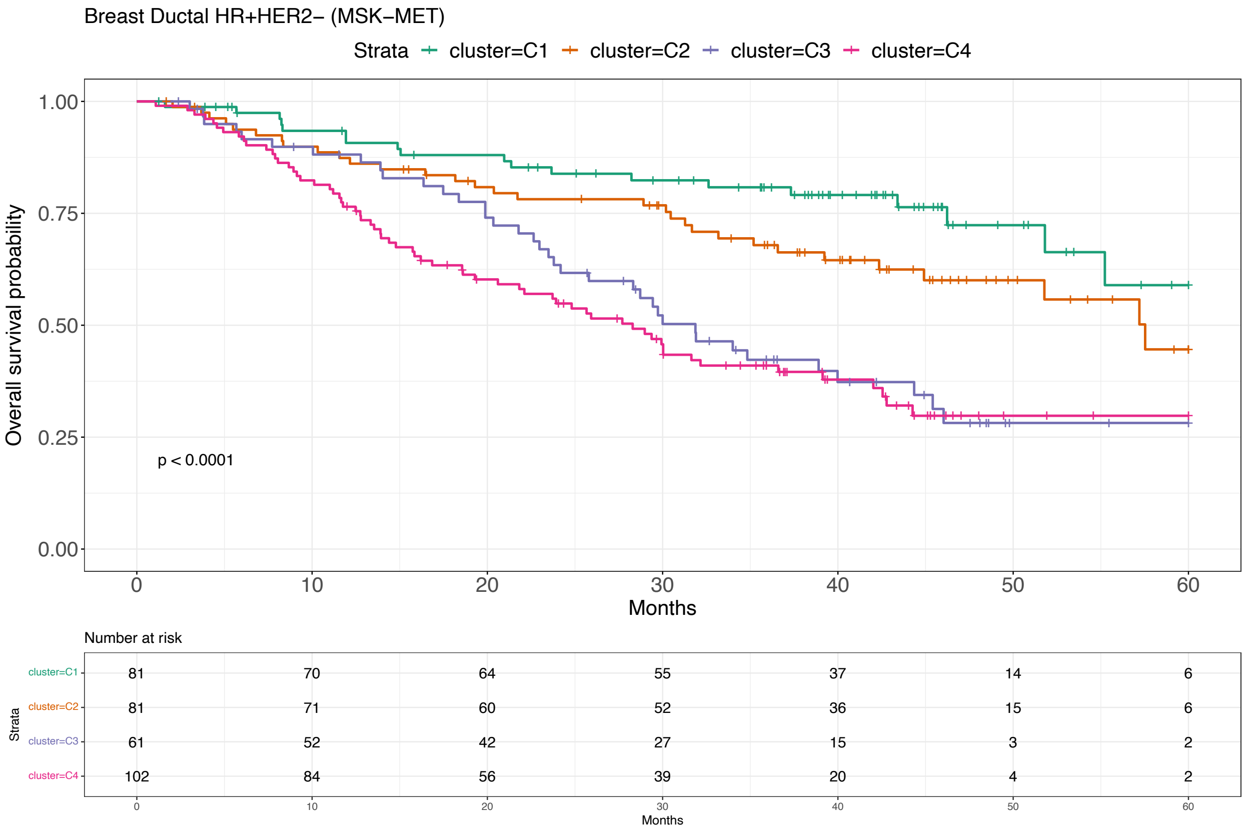

Figure 39: Survival analysis for Breast Ductal (HR+HER2-, MSK-MET) [21].

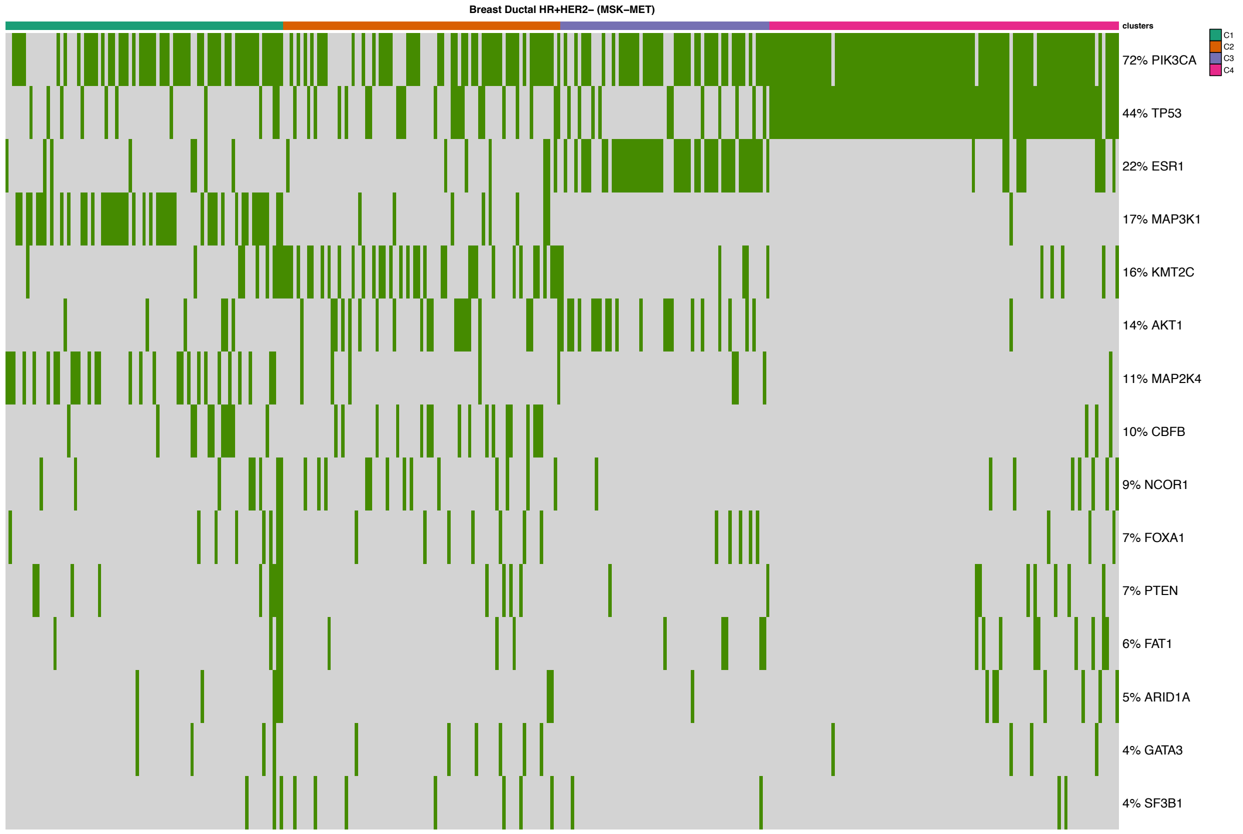

Figure 40: Mutational profile for Breast Ductal (HR+HER2-, MSK-MET) [21].

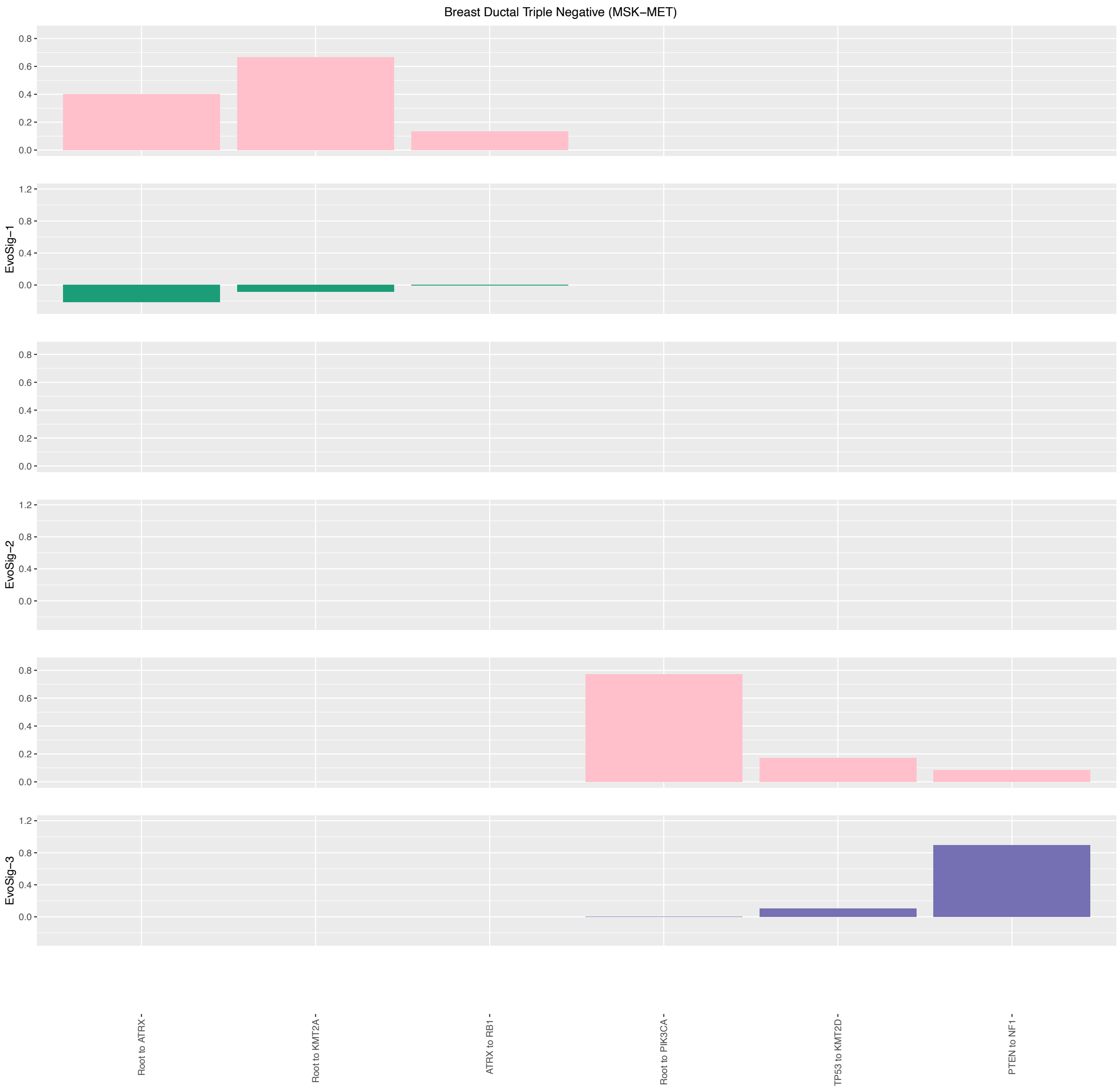

Figure 41: Evolutionary Signatures for Breast Ductal (Triple Negative, MSK-MET) [21].

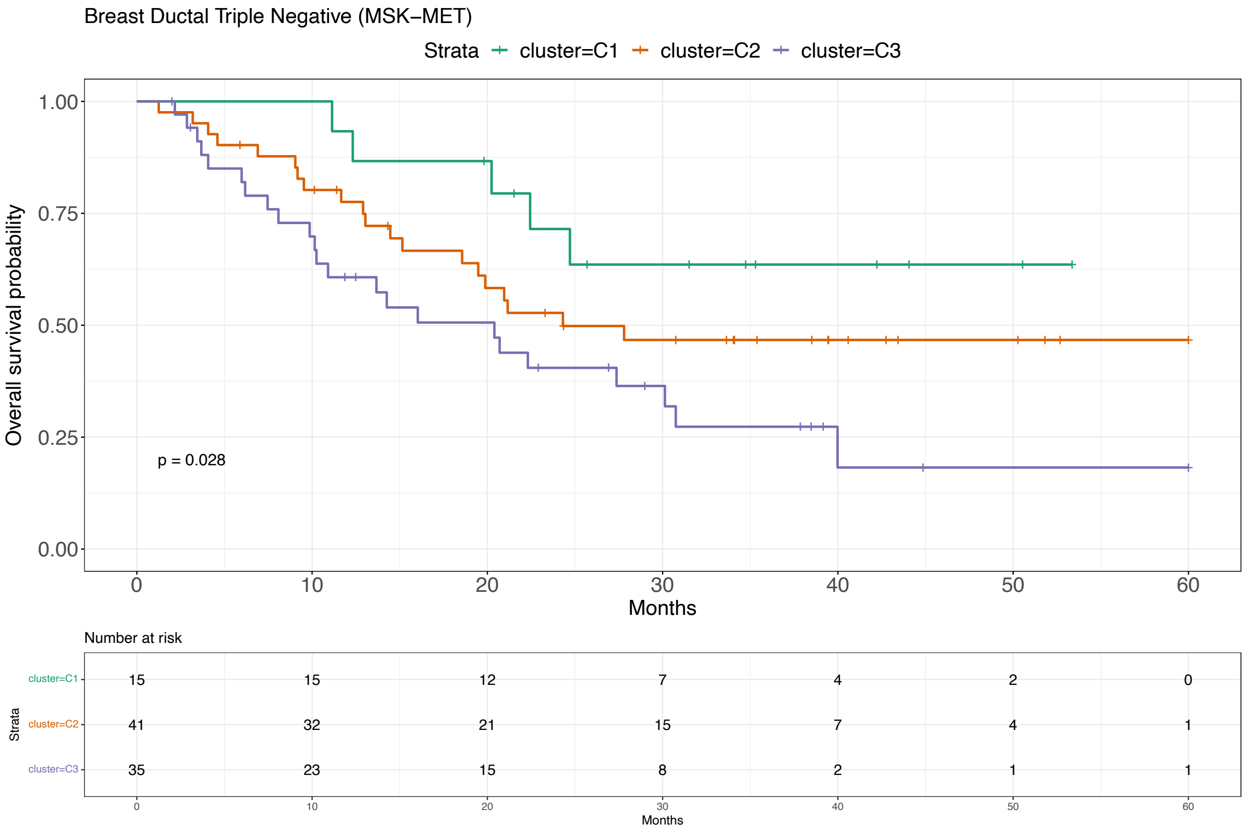

Figure 42: Survival analysis for Breast Ductal (Triple Negative, MSK-MET) [21].

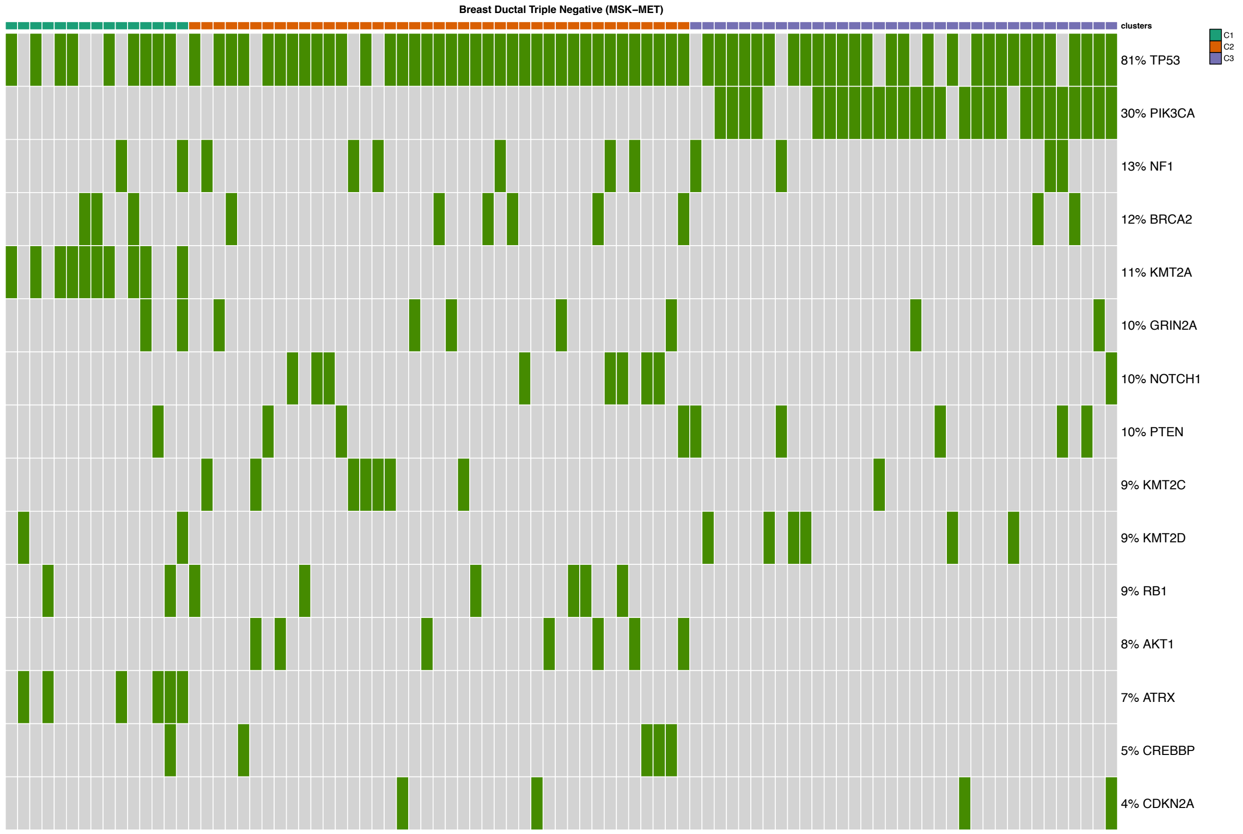

Figure 43: Mutational profile for Breast Ductal (Triple Negative, MSK-MET) [21].

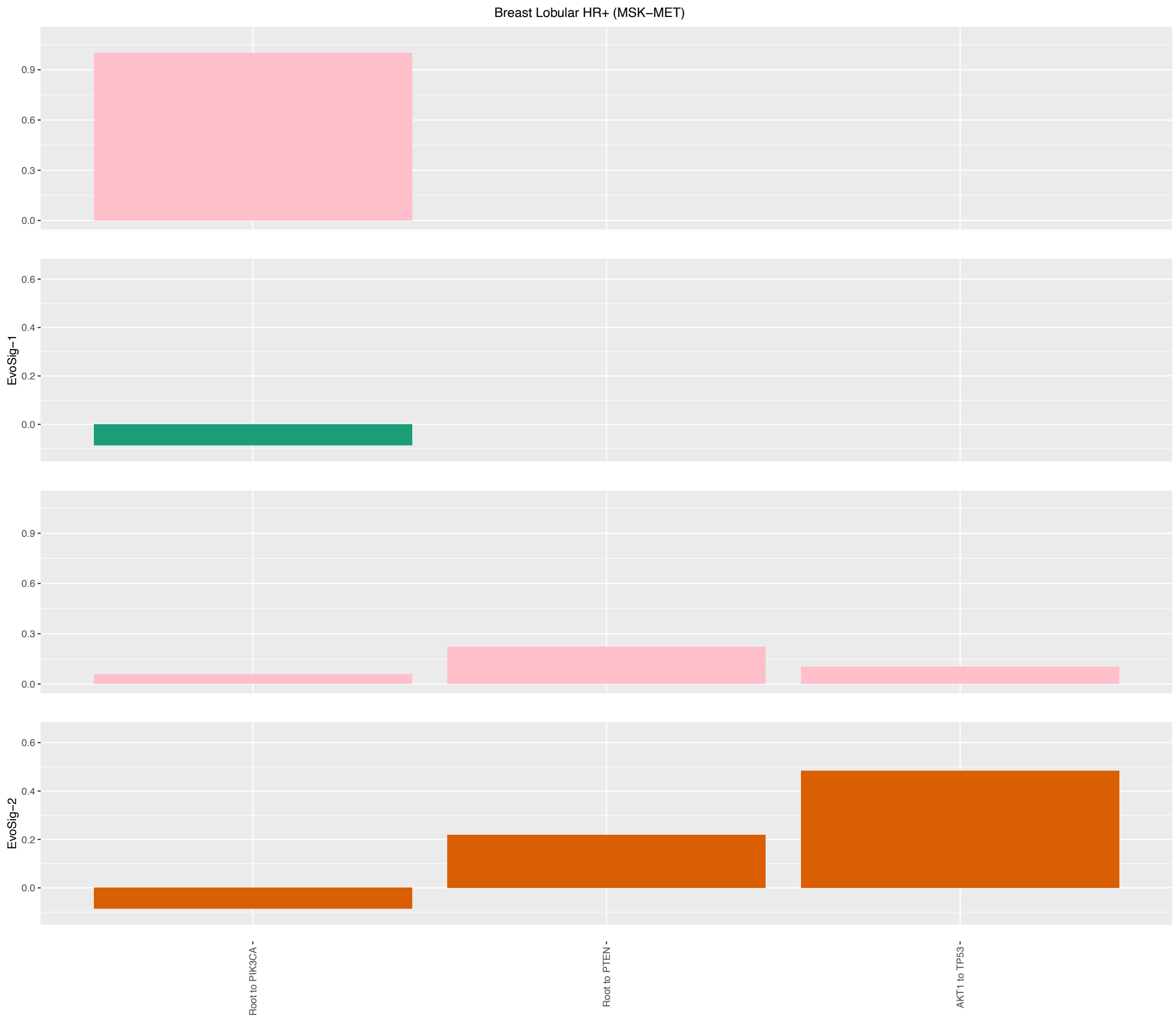

Figure 44: Evolutionary Signatures for Breast Lobular (HR+, MSK-MET) [21].

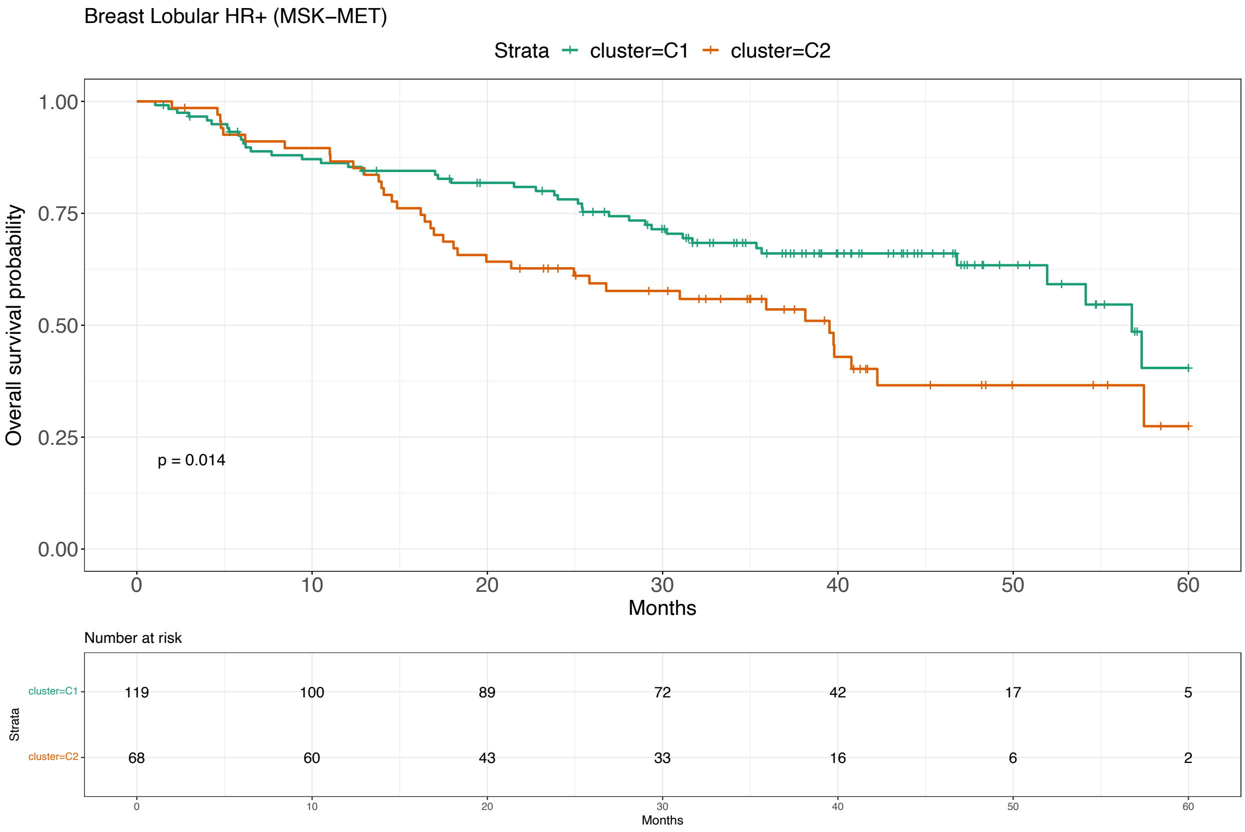

Figure 45: Survival analysis for Breast Lobular (HR+, MSK-MET) [21].

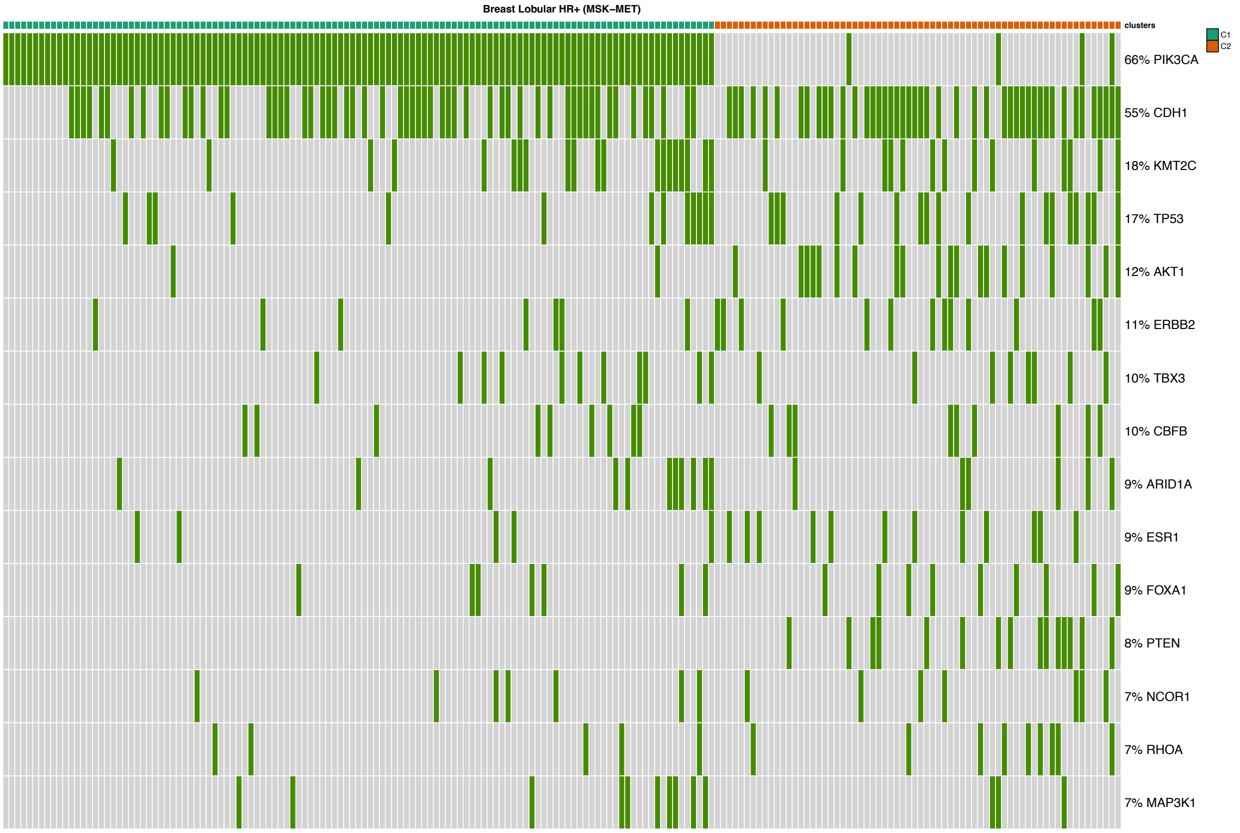

Figure 46: Mutational profile for Breast Lobular (HR+, MSK-MET) [21].

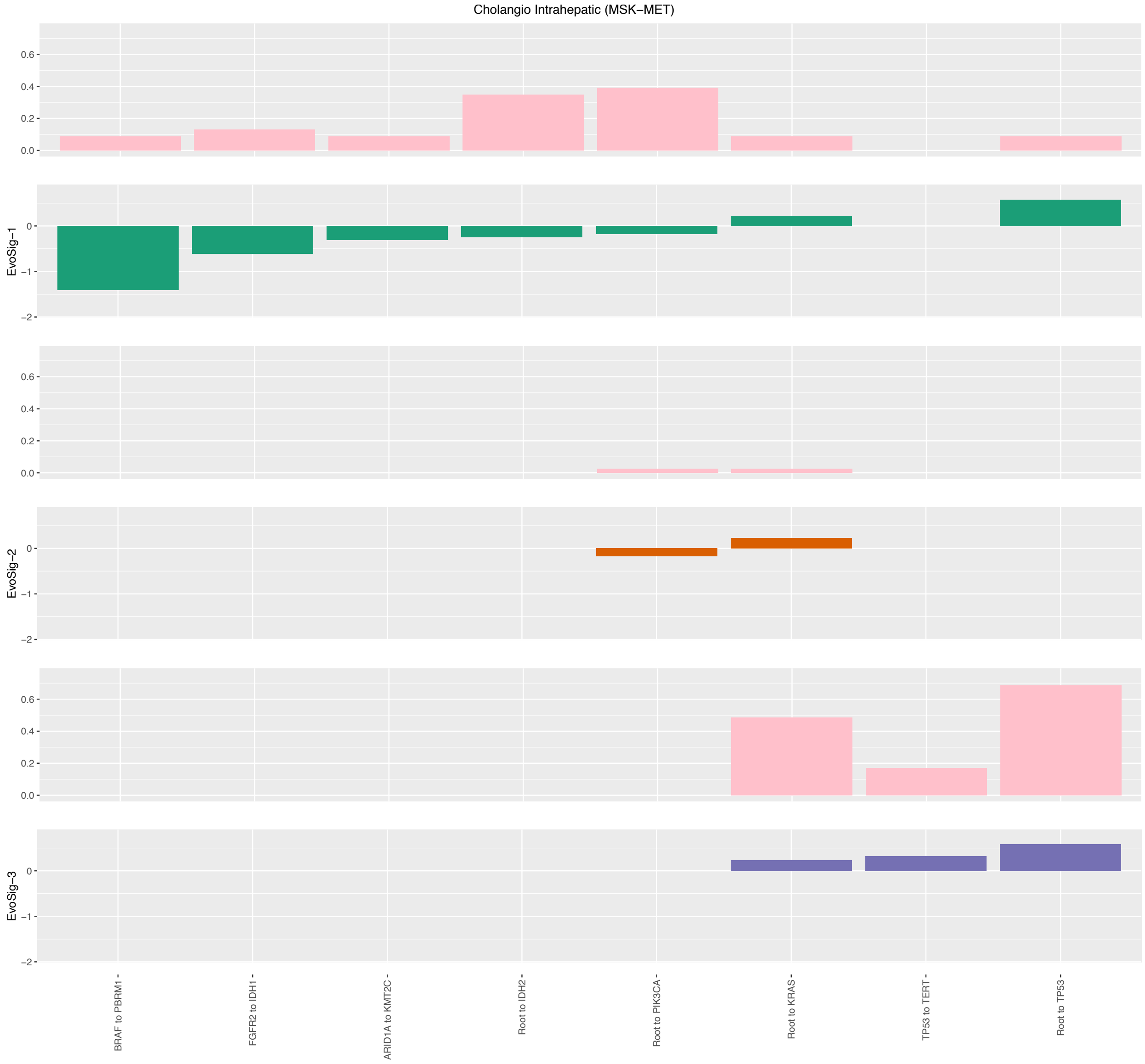

Figure 47: Evolutionary Signatures for Cholangio Intrahepatic (MSK-MET) [21].

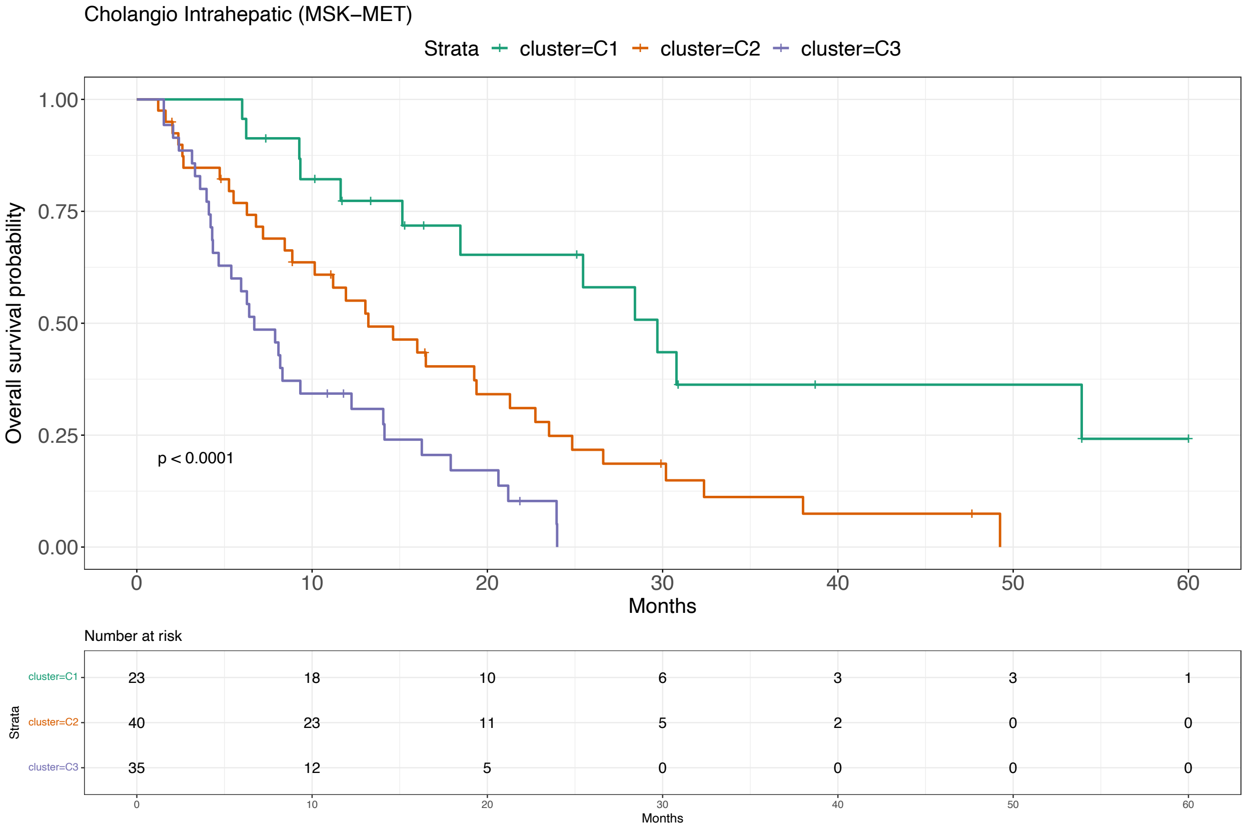

Figure 48: Survival analysis for Cholangio Intrahepatic (MSK-MET) [21].

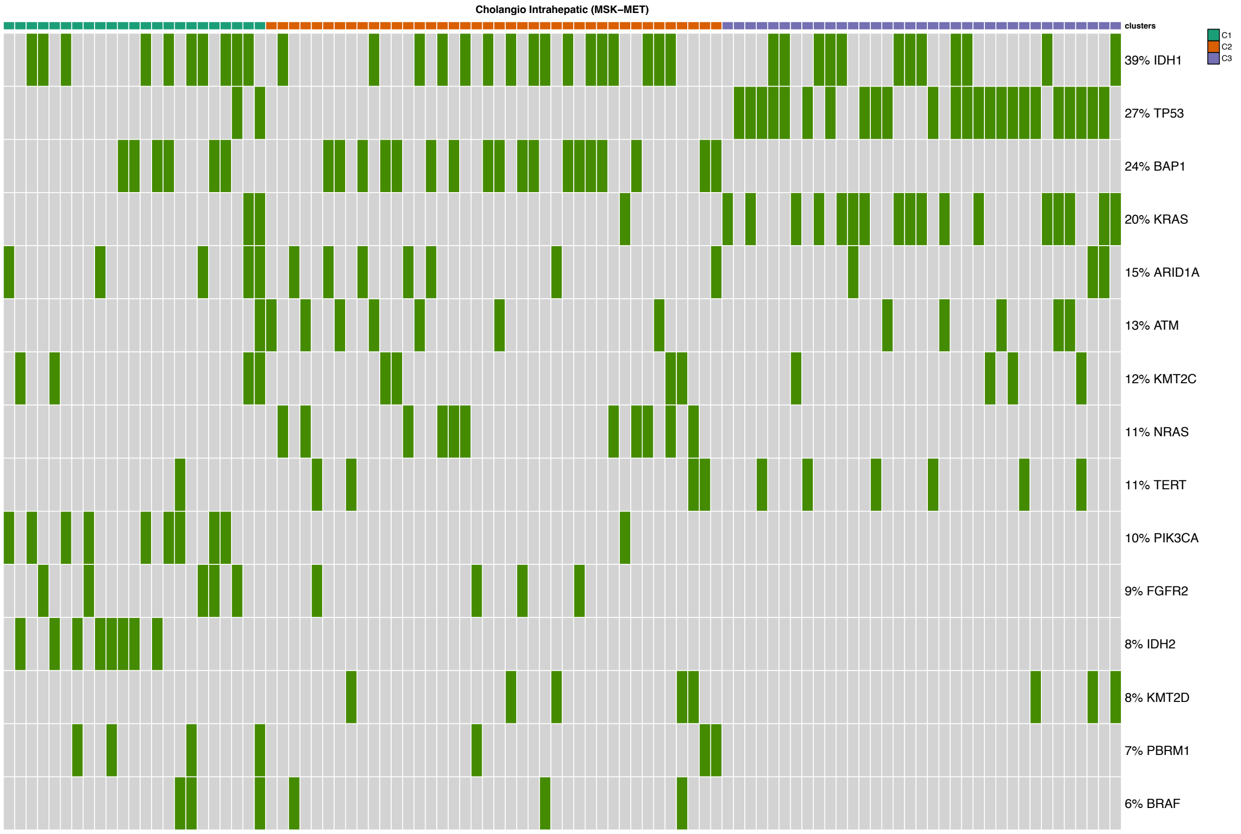

Figure 49: Mutational profile for Cholangio Intrahepatic (MSK-MET) [21].

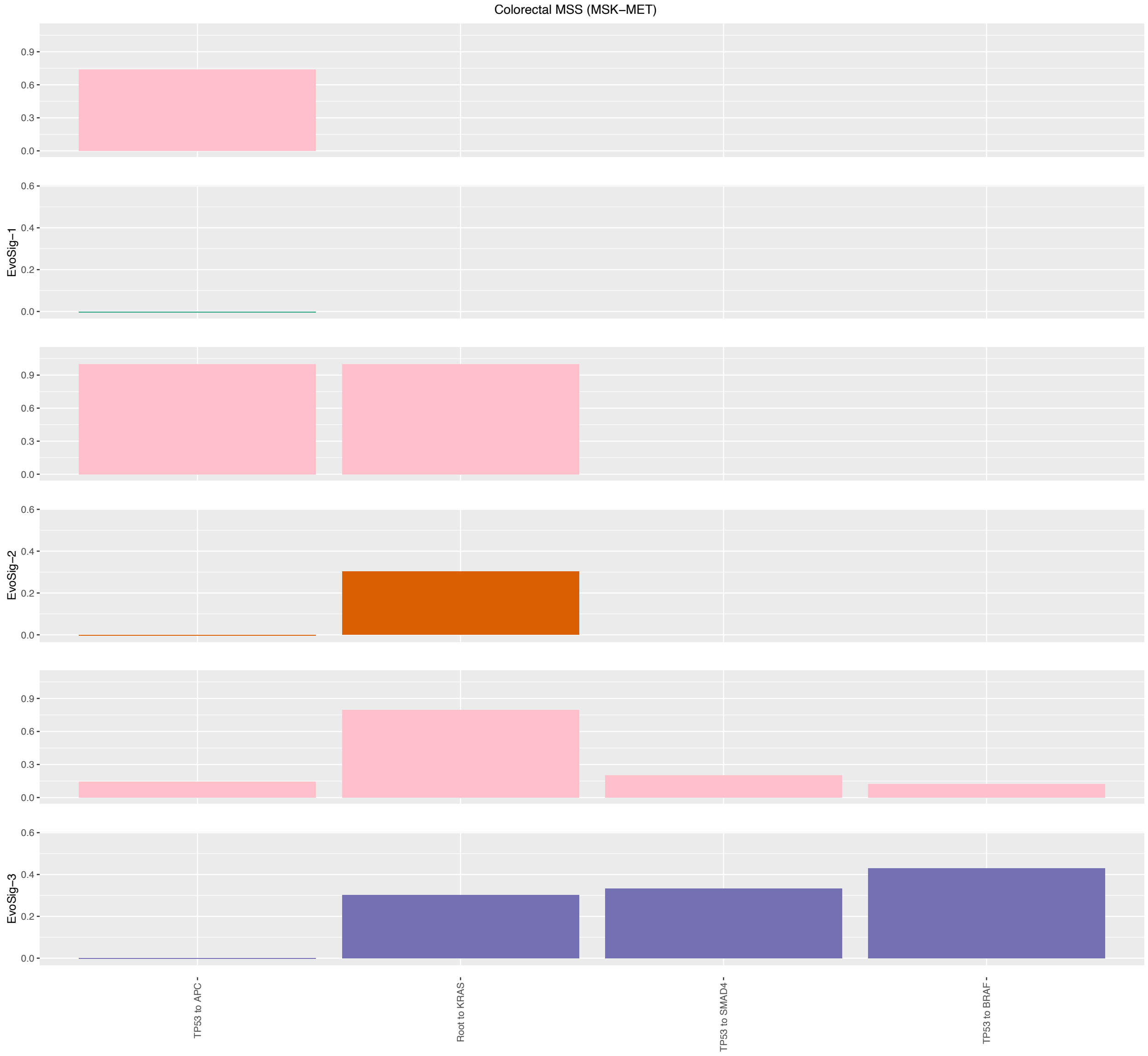

Figure 50: Evolutionary Signatures for Colorectal (MSS, MSK-MET) [21].

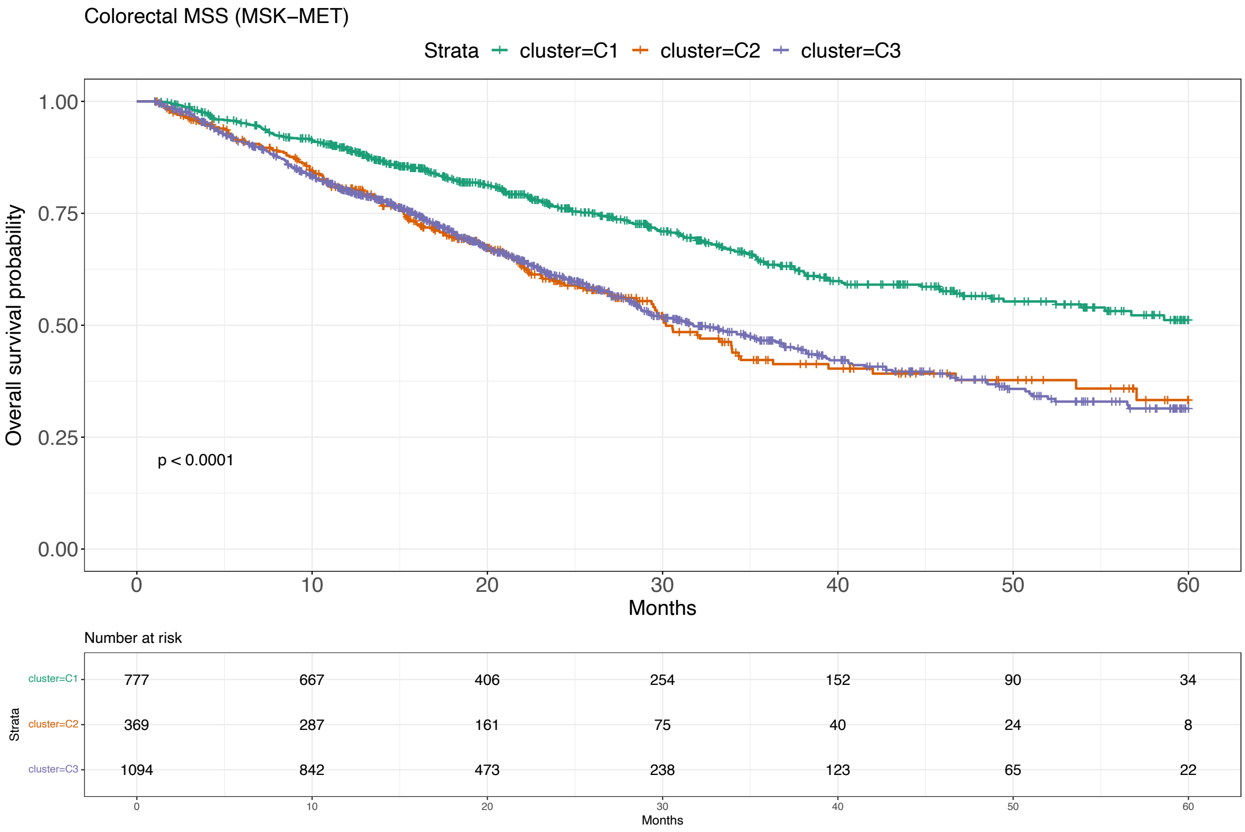

Figure 51: Survival analysis for Colorectal (MSS, MSK-MET) [21].

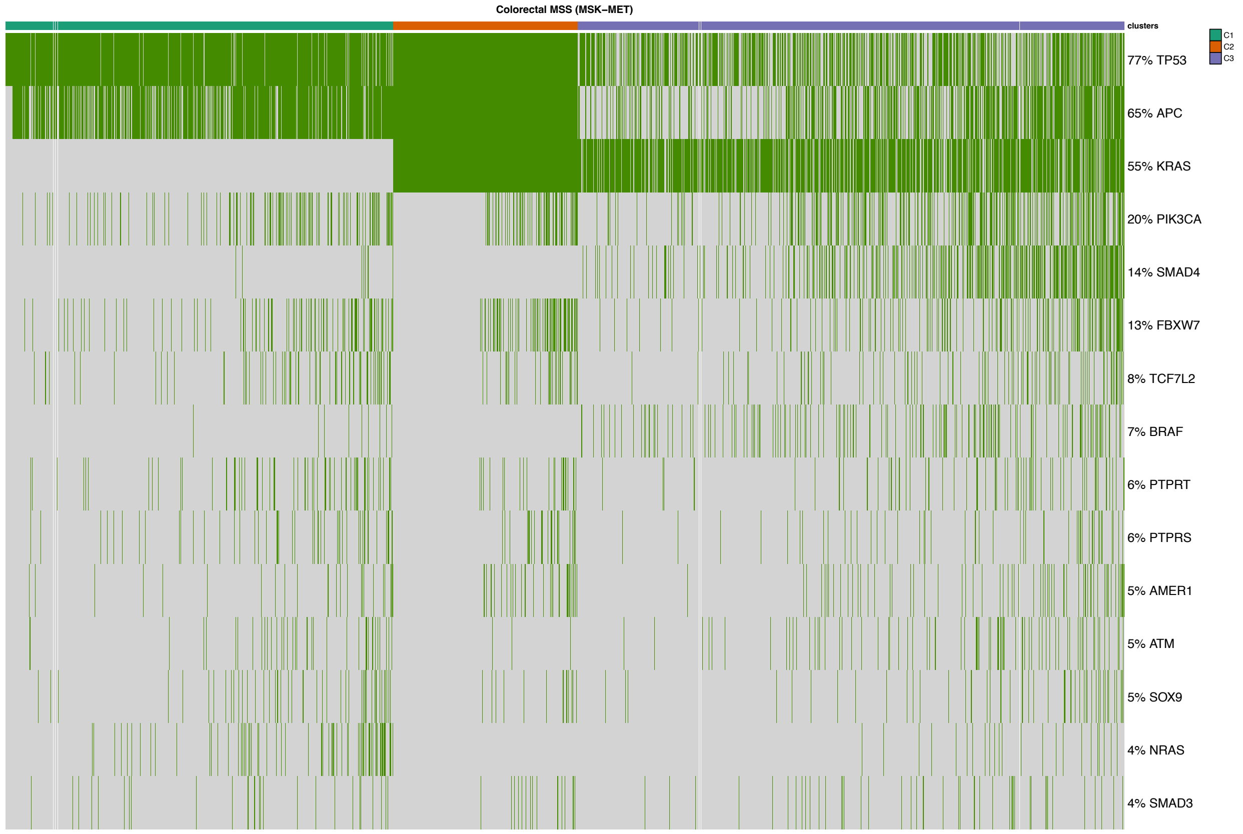

Figure 52: Mutational profile for Colorectal (MSS, MSK-MET) [21].

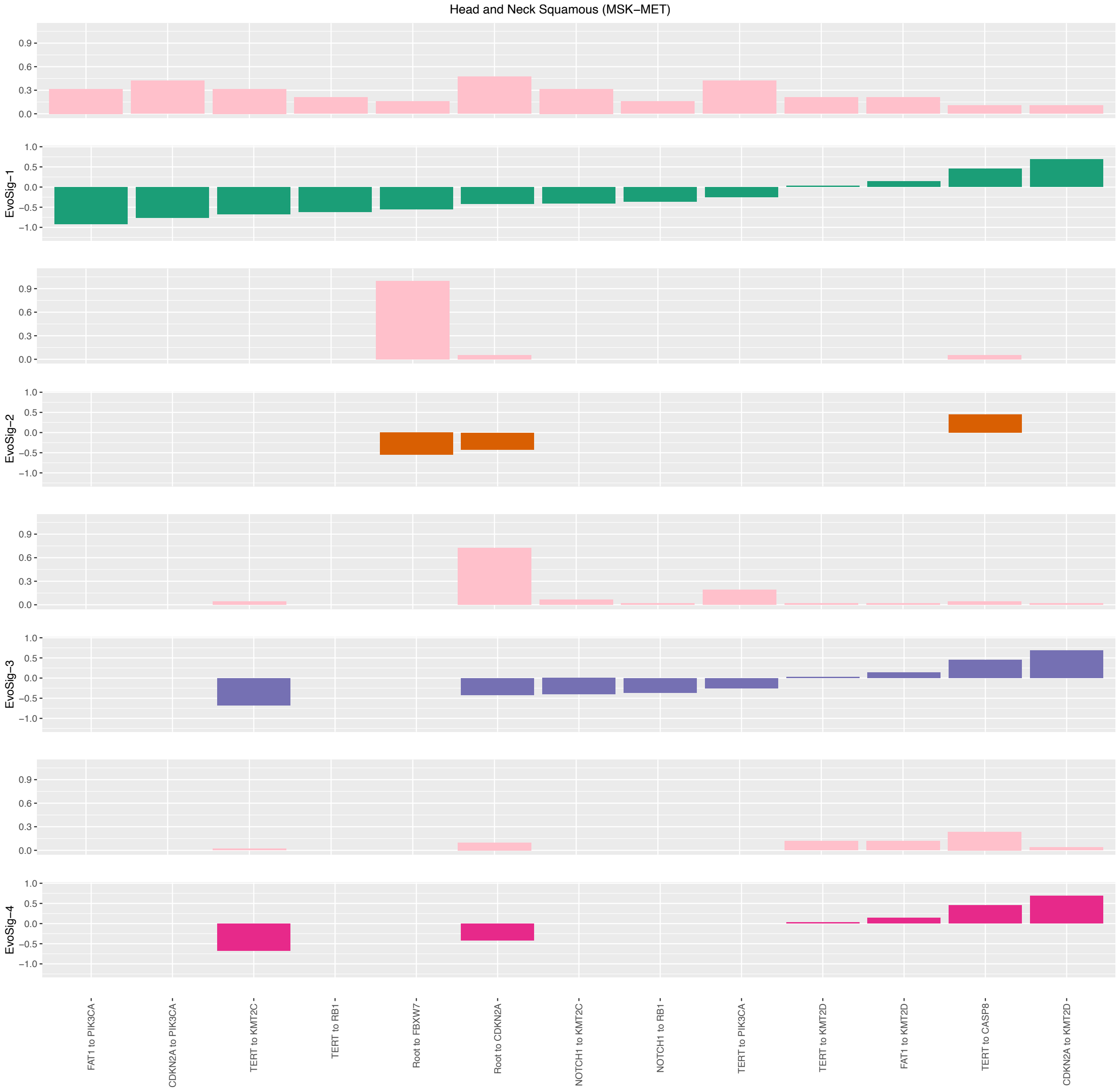

Figure 53: Evolutionary Signatures for Head and Neck Squamous (MSK-MET) [21].

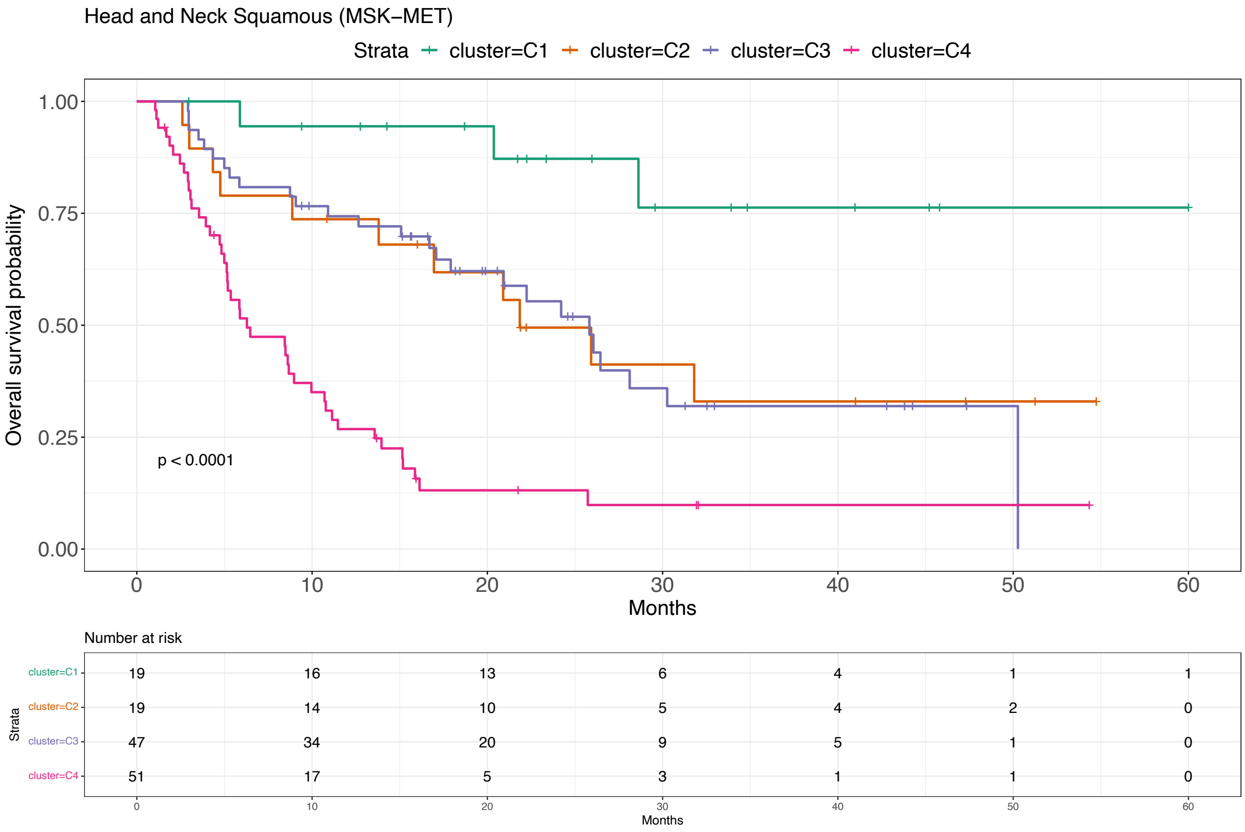

Figure 54: Survival analysis for Head and Neck Squamous (MSK-MET) [21].

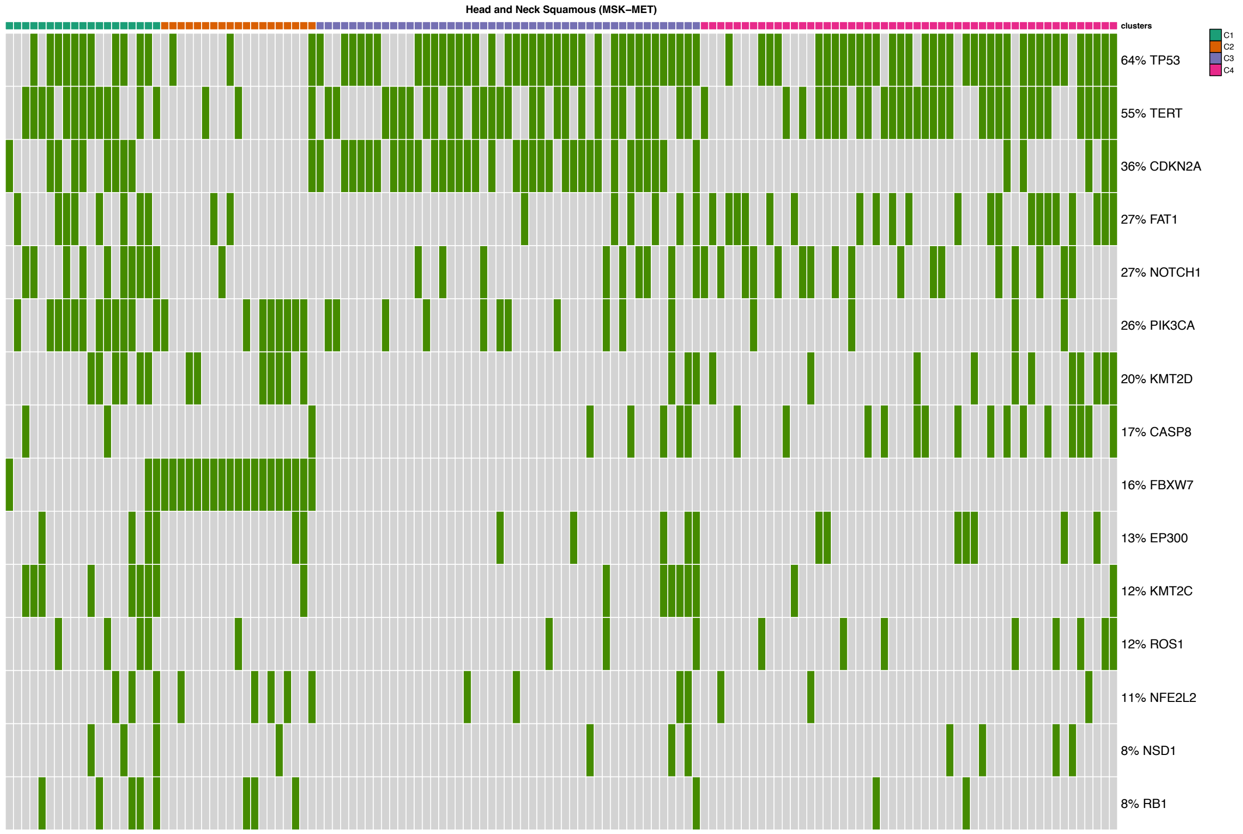

Figure 55: Mutational profile for Head and Neck Squamous (MSK-MET) [21].

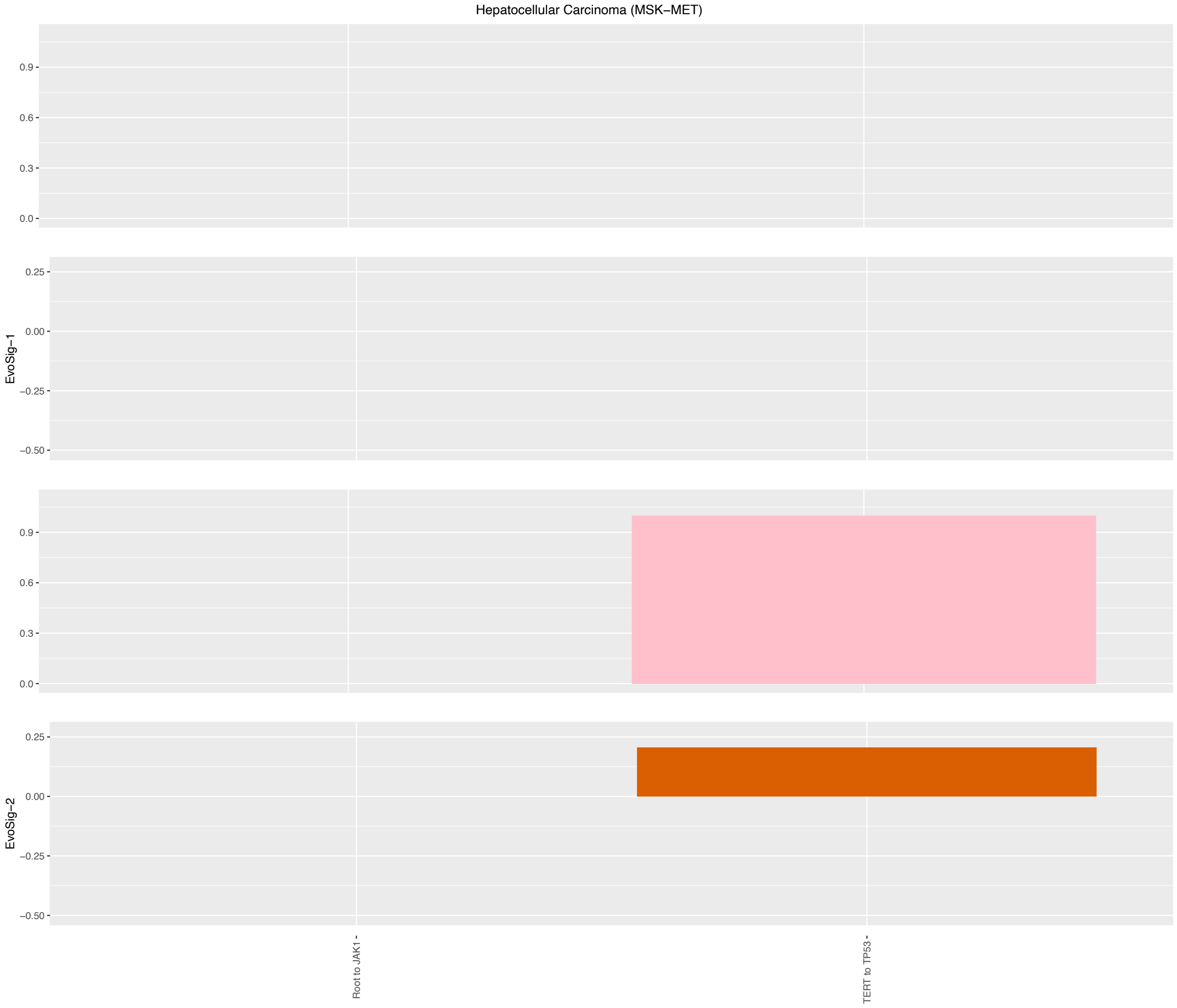

Figure 56: Evolutionary Signatures for Hepatocellular Carcinoma (MSK-MET) [21].

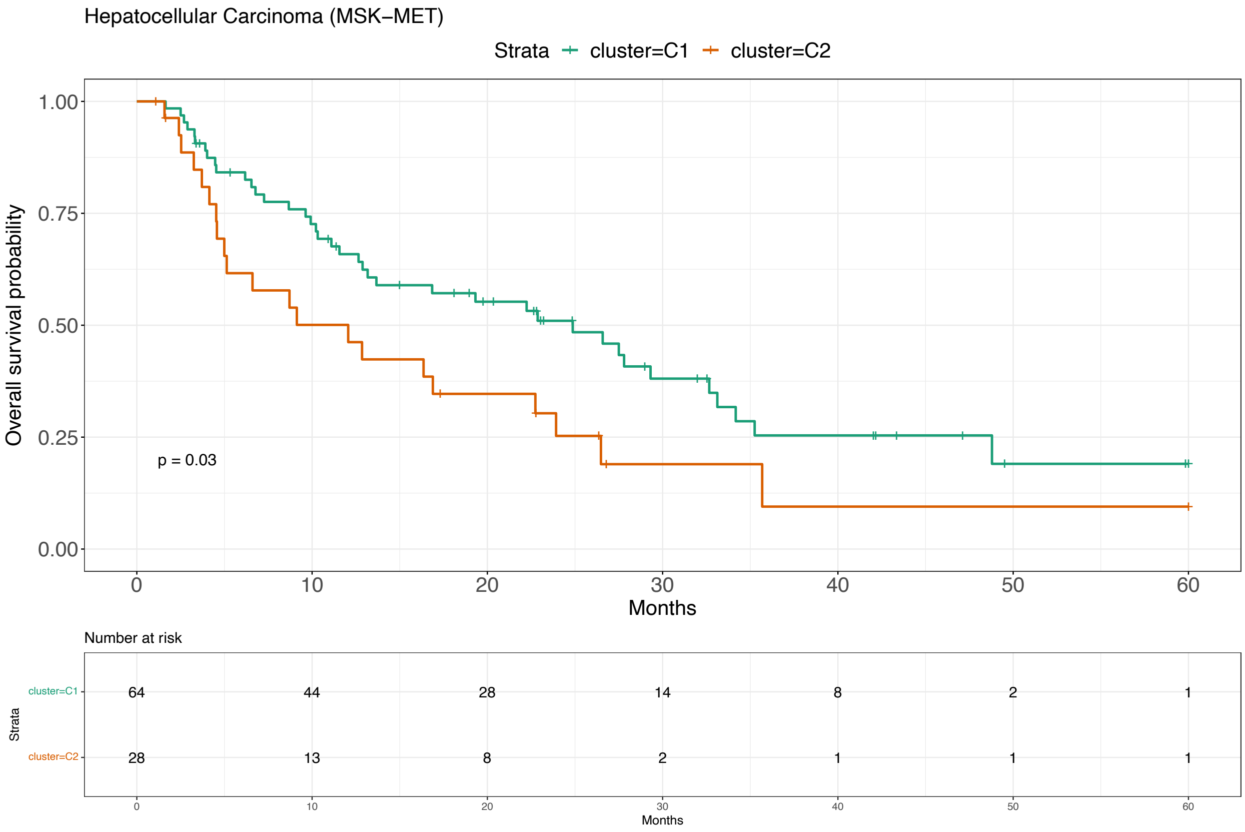

Figure 57: Survival analysis for Hepatocellular Carcinoma (MSK-MET) [21].

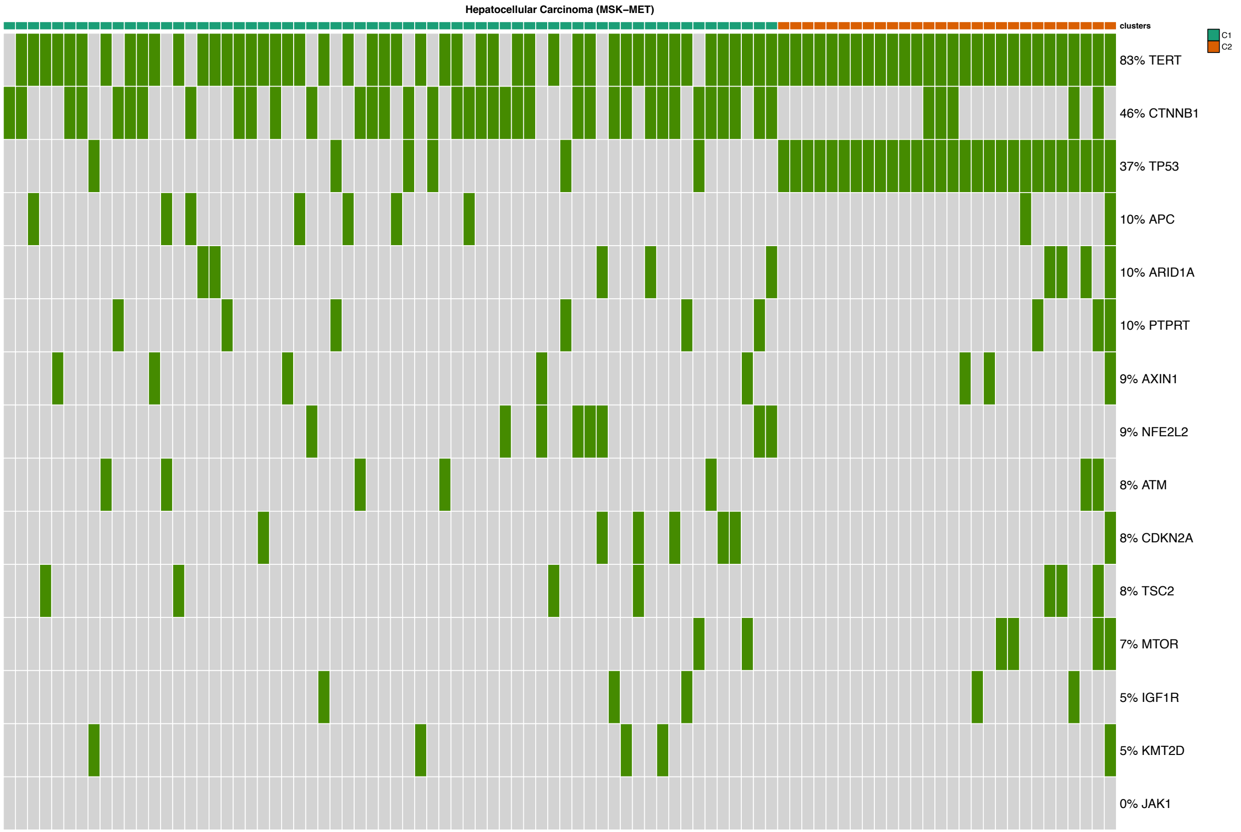

Figure 58: Mutational profile for Hepatocellular Carcinoma (MSK-MET) [21].

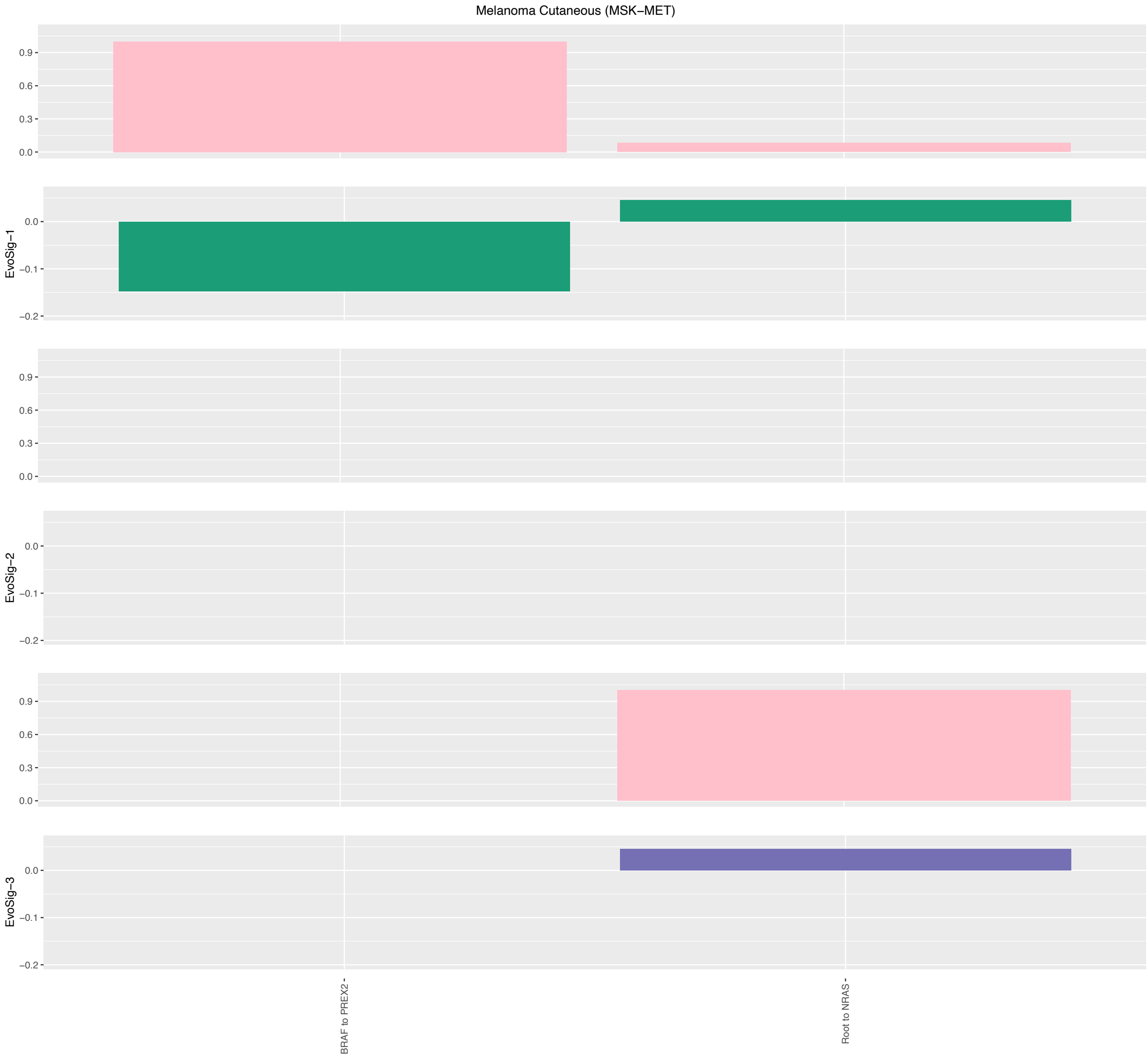

Figure 59: Evolutionary Signatures for Melanoma Cutaneous (MSK-MET) [21].

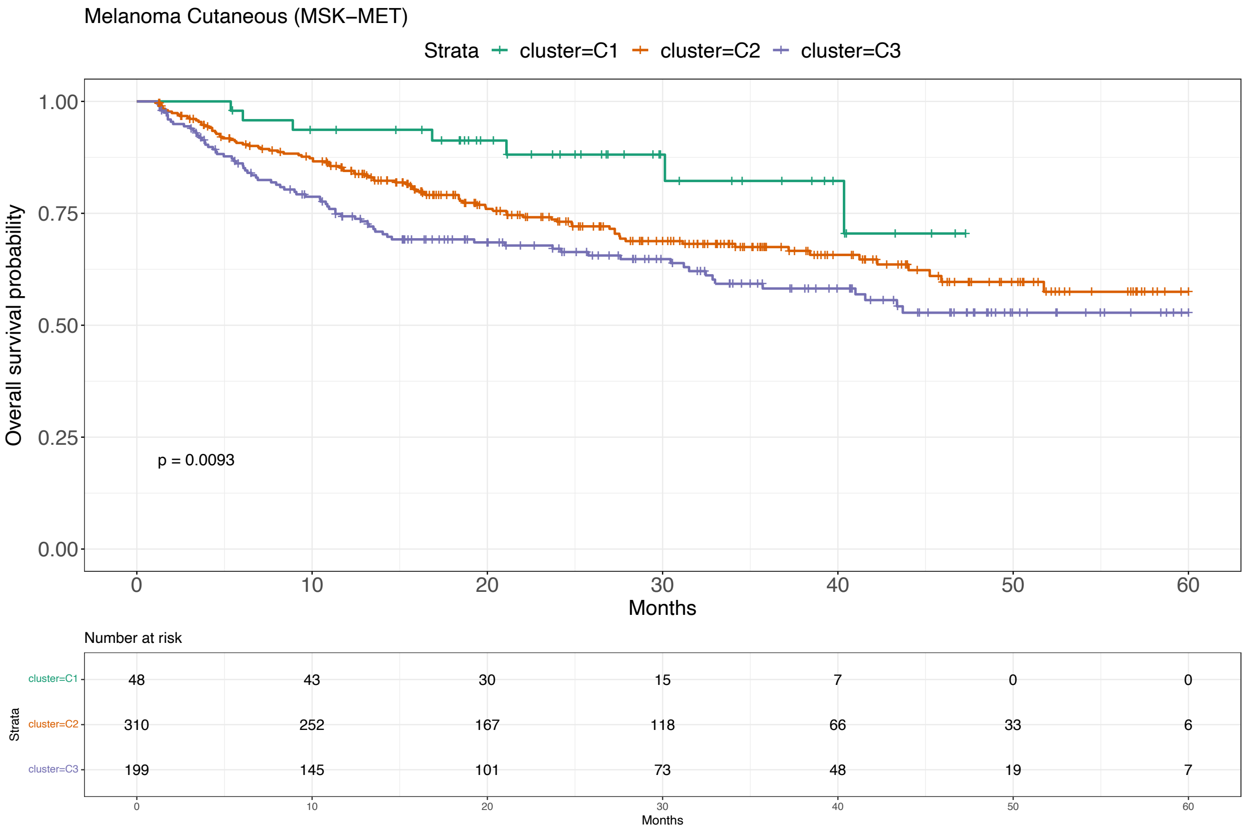

Figure 60: Survival analysis for Melanoma Cutaneous (MSK-MET) [21].

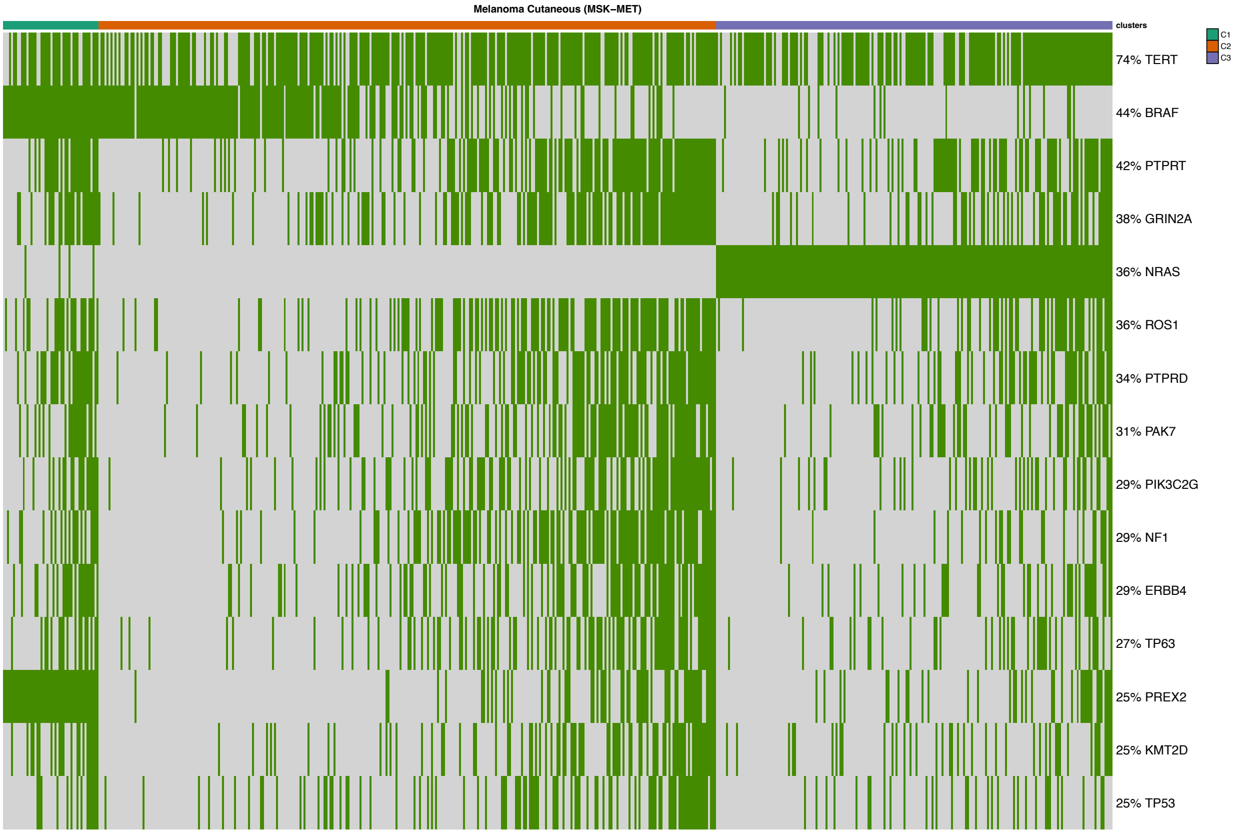

Figure 61: Mutational profile for Melanoma Cutaneous (MSK-MET) [21].

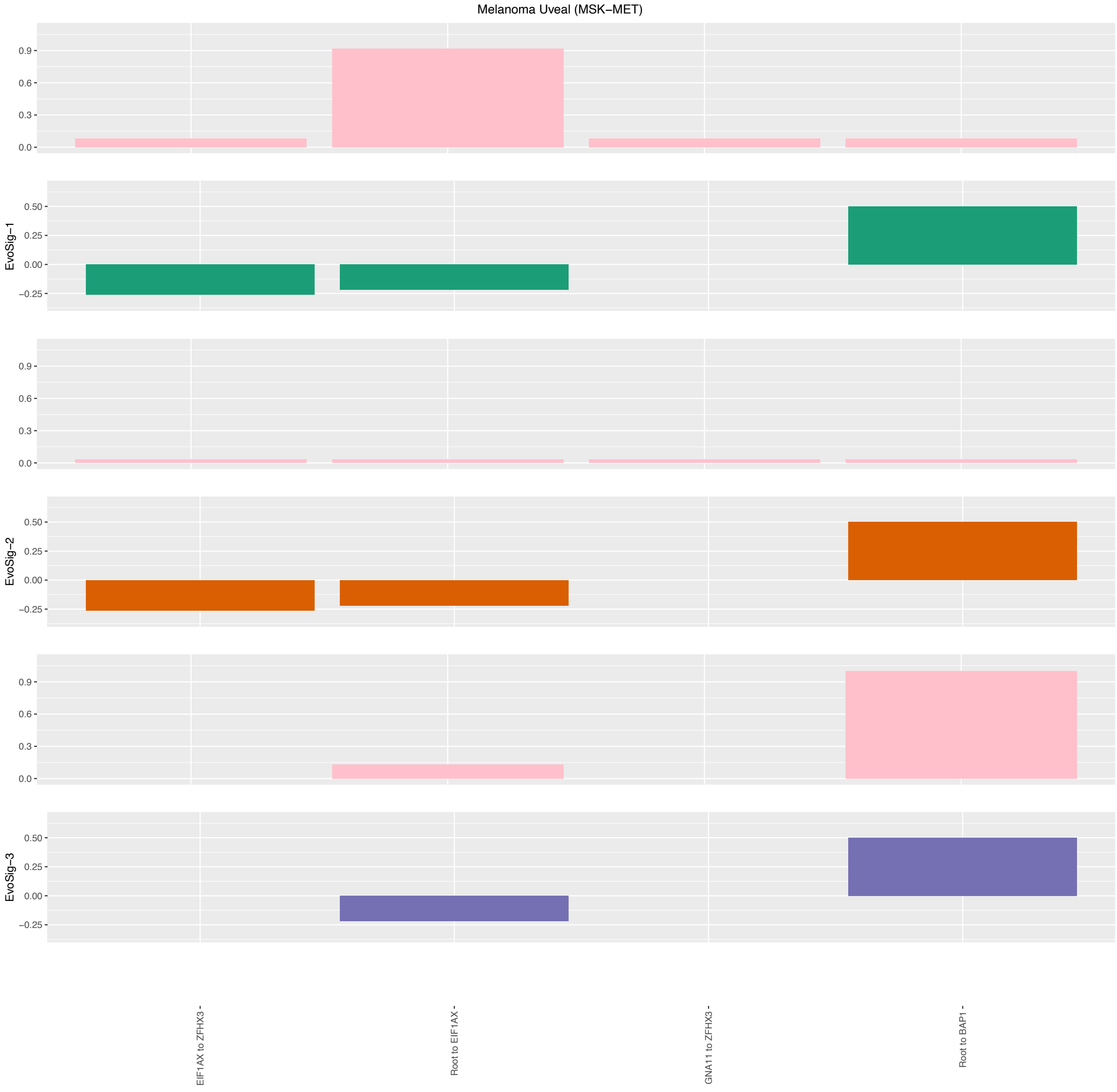

Figure 62: Evolutionary Signatures for Melanoma Uveal (MSK-MET) [21].

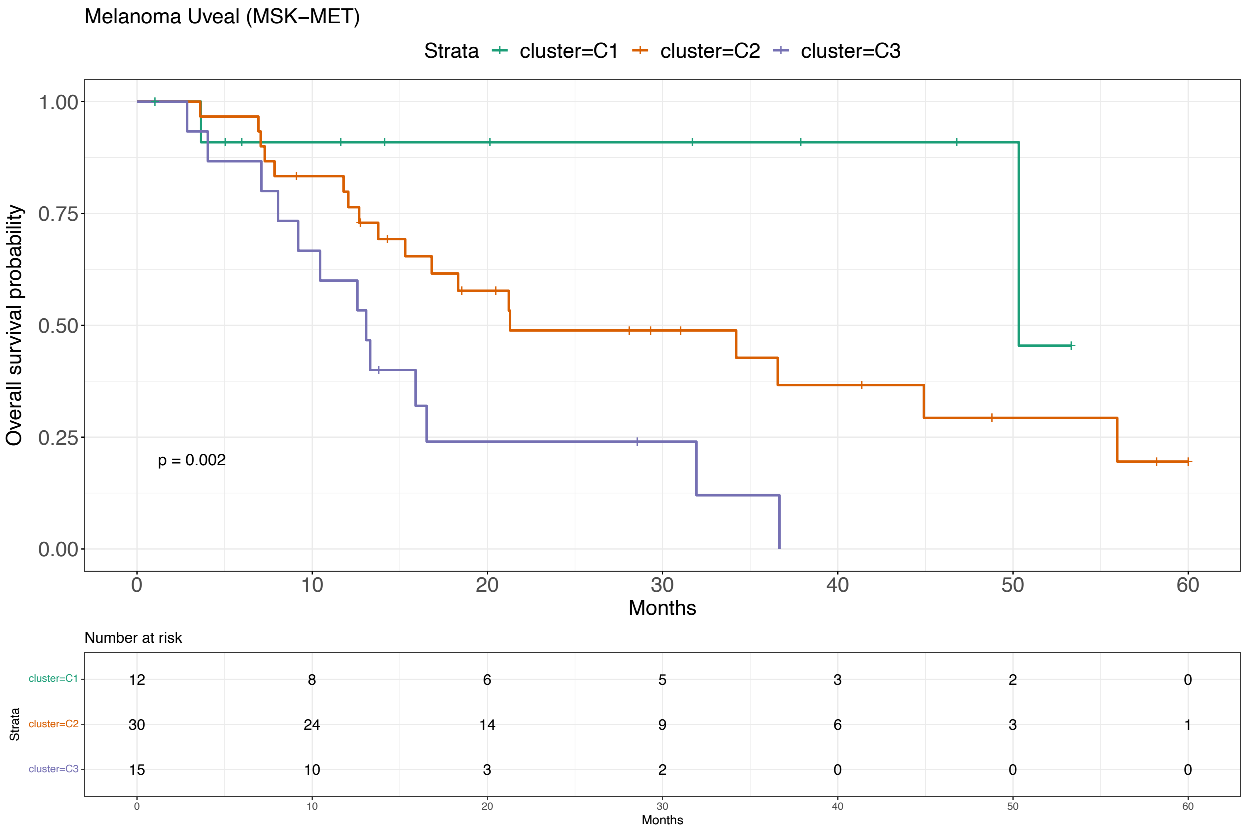

Figure 63: Survival analysis for Melanoma Uveal (MSK-MET) [21].

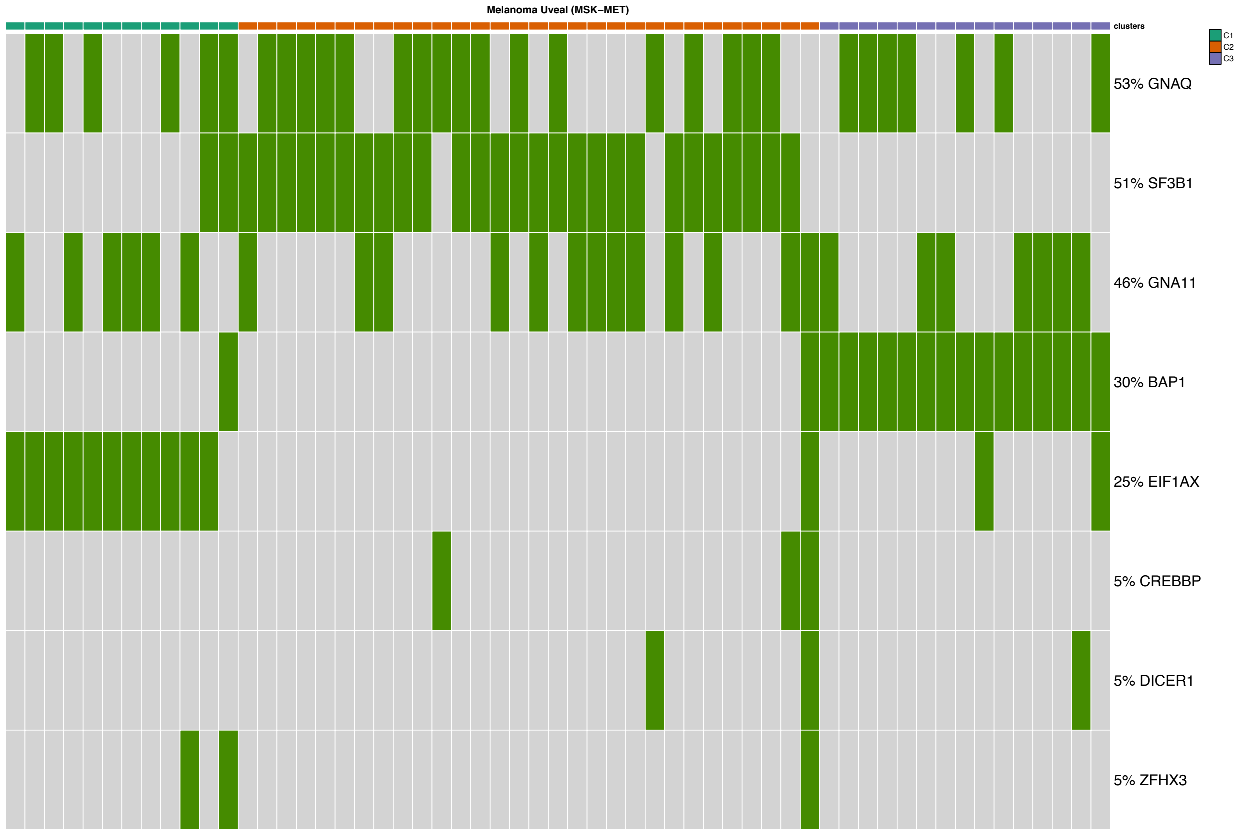

Figure 64: Mutational profile for Melanoma Uveal (MSK-MET) [21].

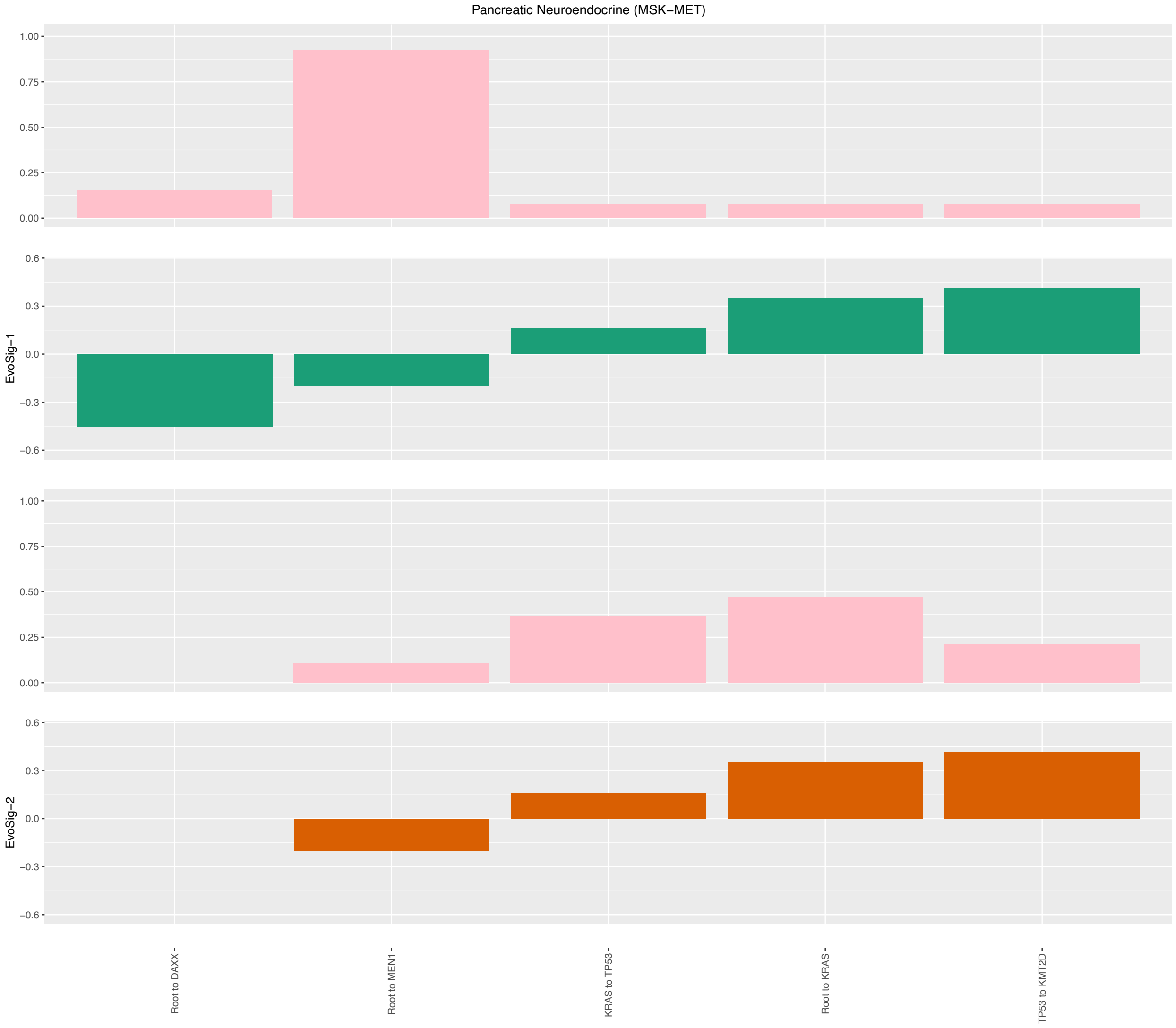

Figure 65: Evolutionary Signatures for Pancreatic Neuroendocrine (MSK-MET) [21].

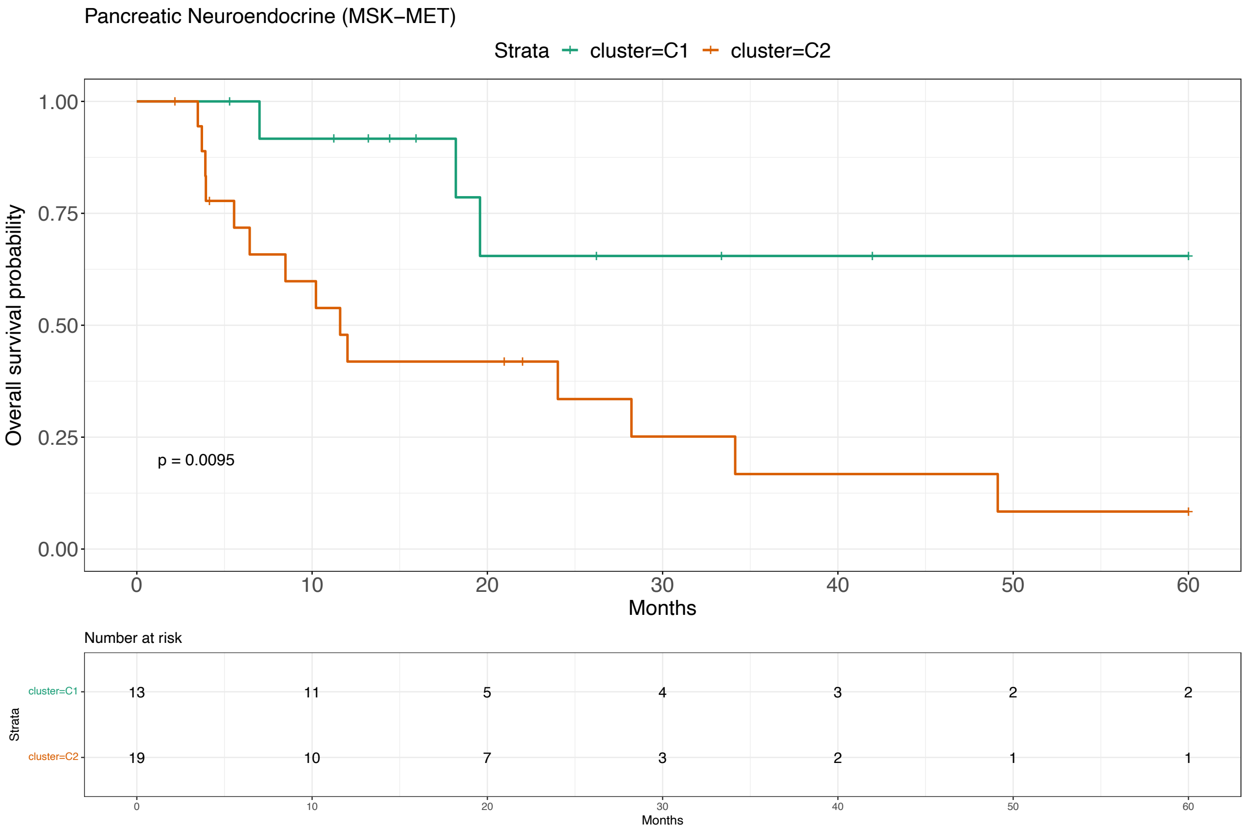

Figure 66: Survival analysis for Pancreatic Neuroendocrine (MSK-MET) [21].

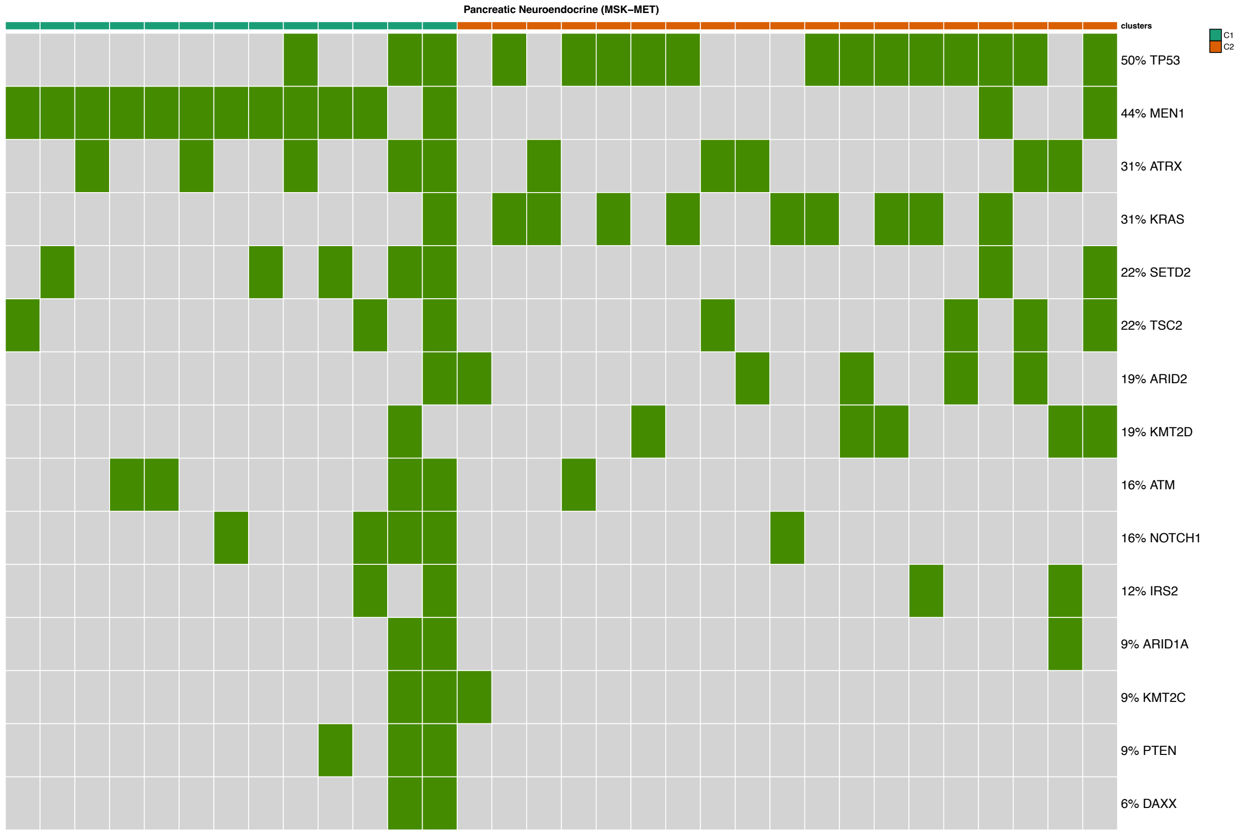

Figure 67: Mutational profile for Pancreatic Neuroendocrine (MSK-MET) [21].

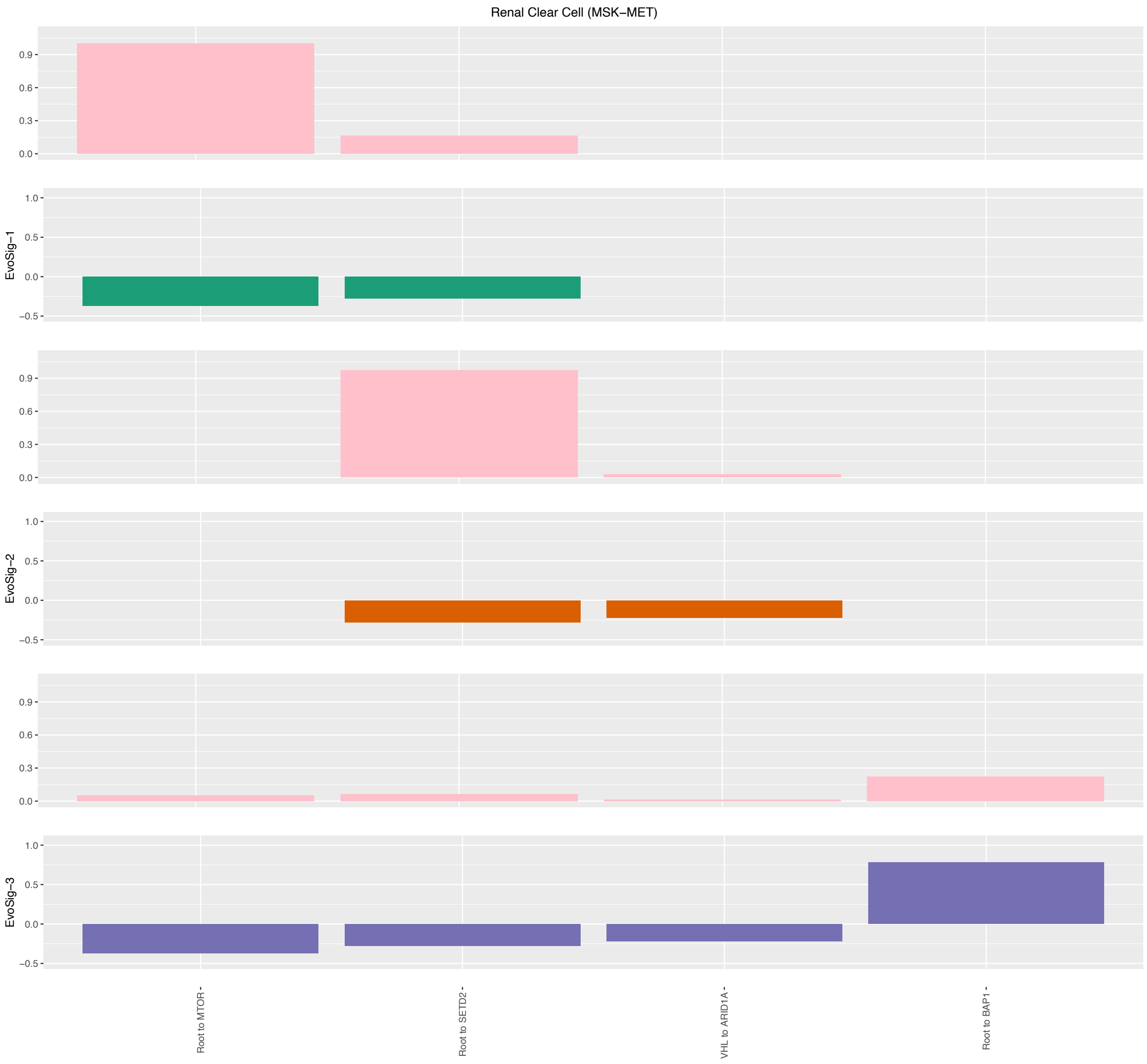

Figure 68: Evolutionary Signatures for Renal Clear Cell (MSK-MET) [21].

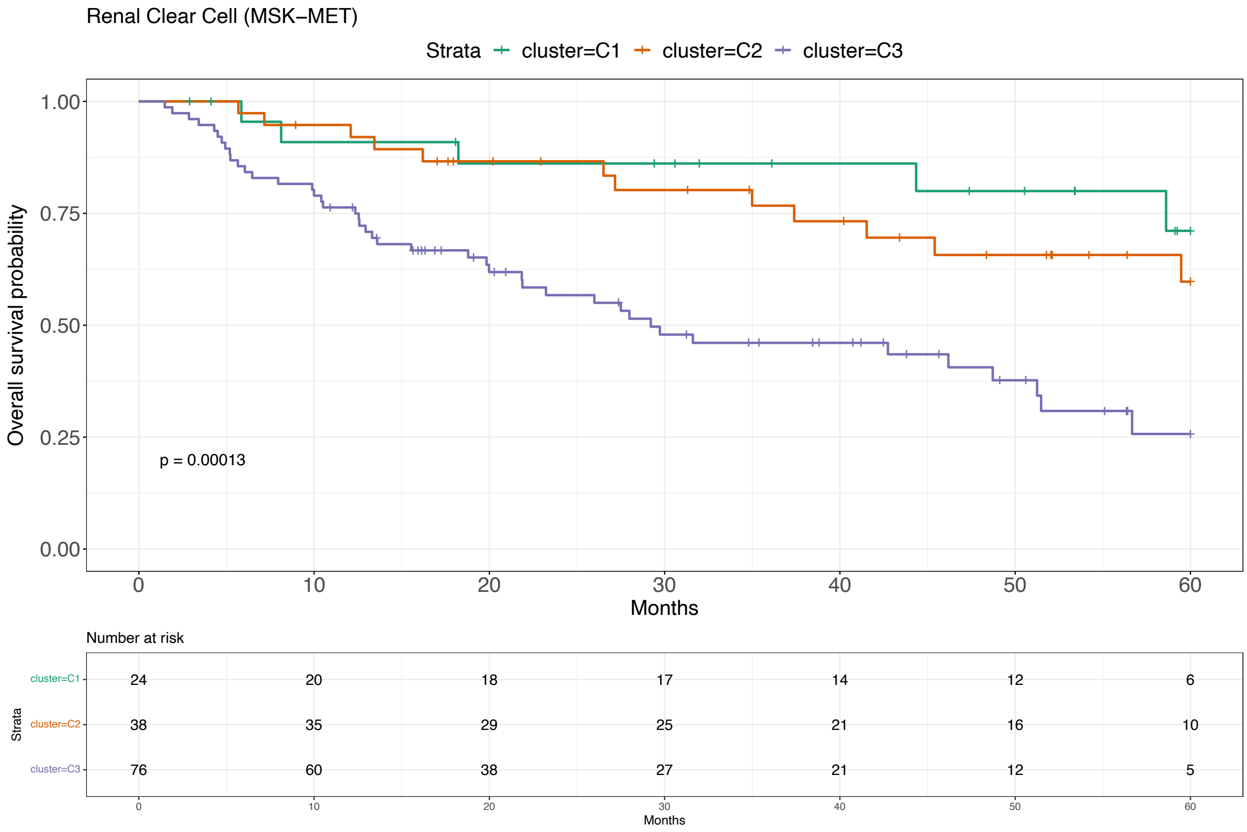

Figure 69: Survival analysis for Renal Clear Cell (MSK-MET) [21].

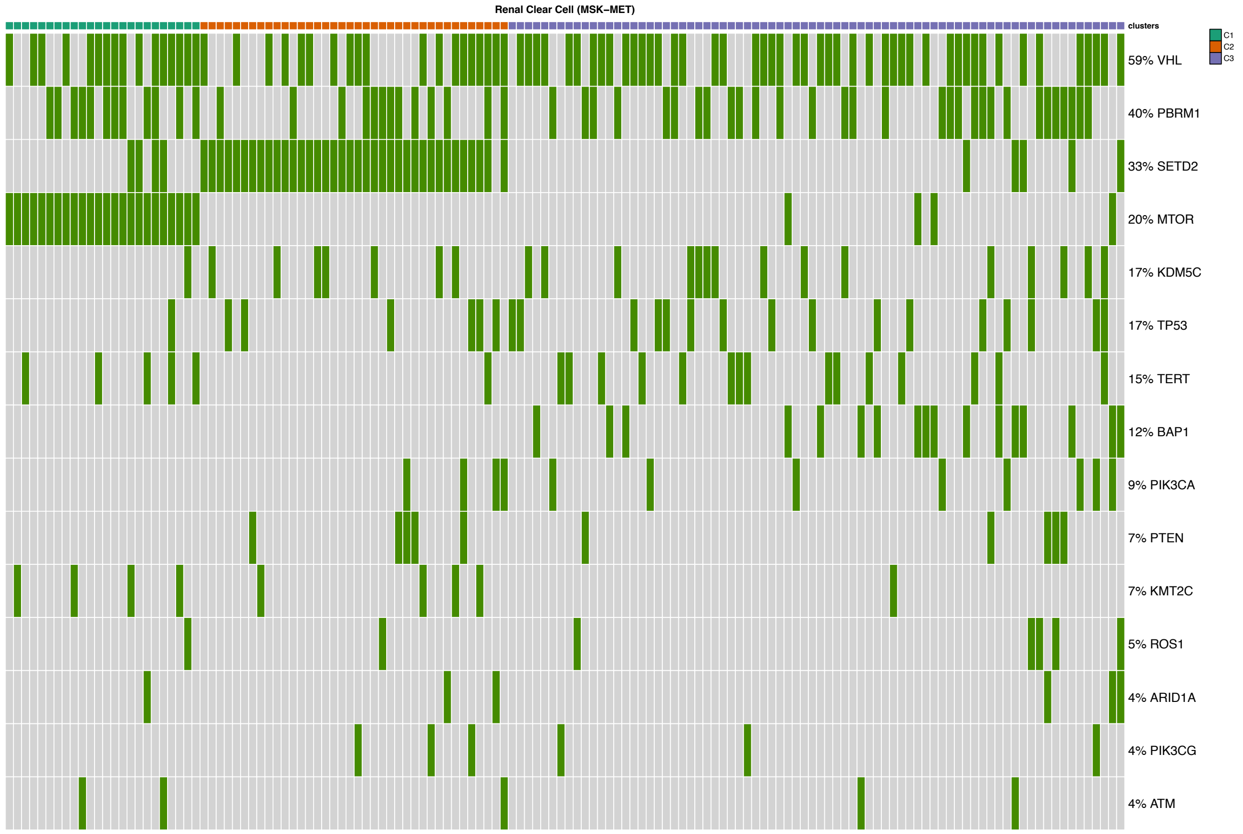

Figure 70: Mutational profile for Renal Clear Cell (MSK-MET) [21].

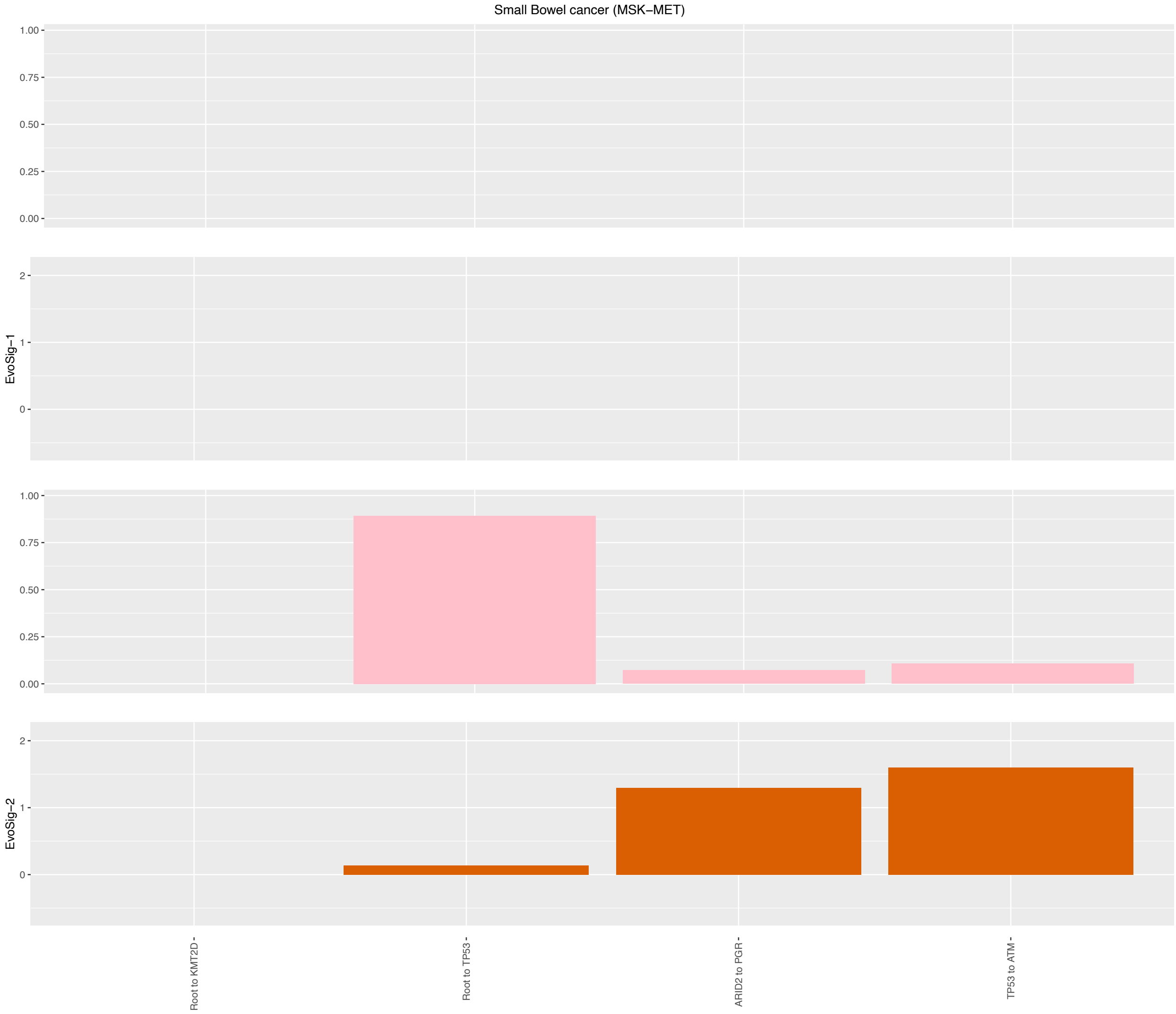

Figure 71: Evolutionary Signatures for Small Bowel Cancer (MSK-MET) [21].

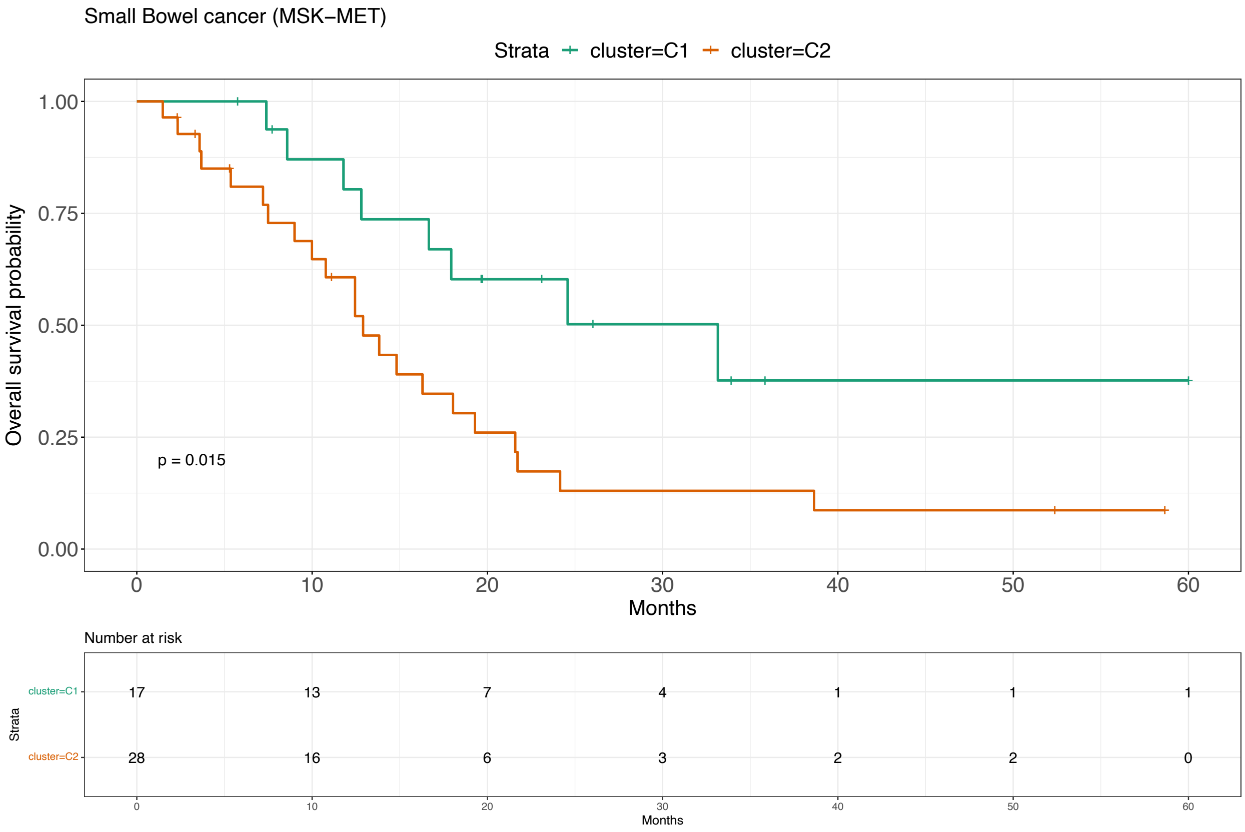

Figure 72: Survival analysis for Small Bowel Cancer (MSK-MET) [21].

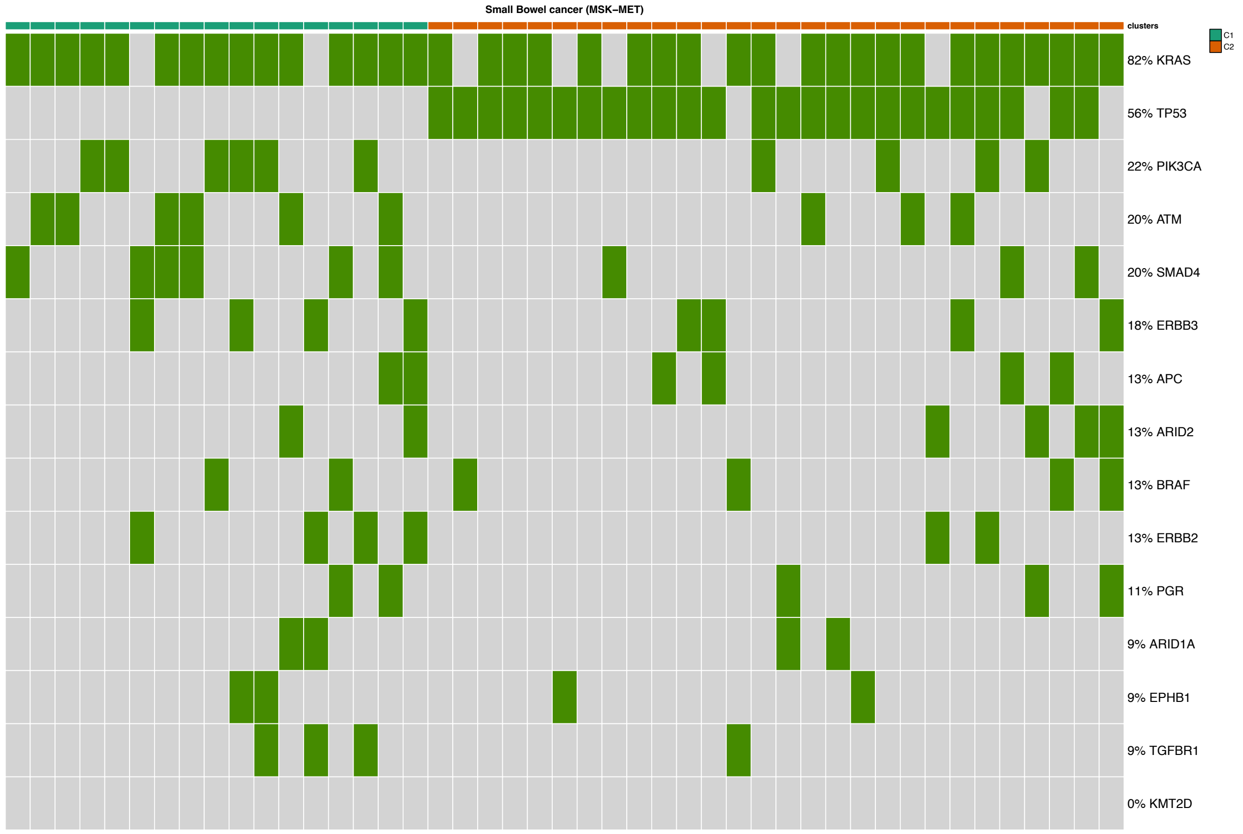

Figure 73: Mutational profile for Small Bowel Cancer (MSK-MET) [21].

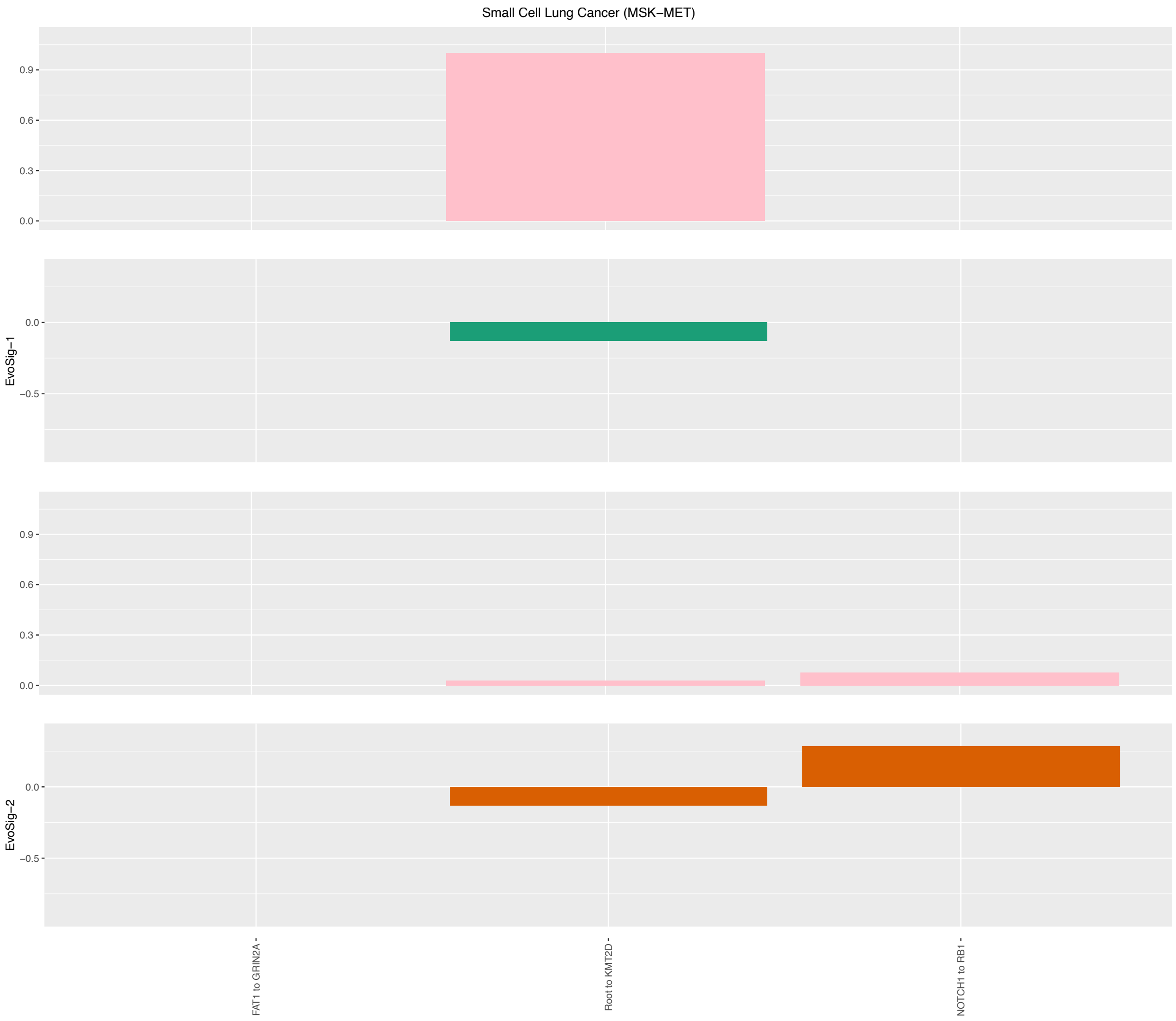

Figure 74: Evolutionary Signatures for Small-Cell Lung Cancer (MSK-MET) [21].

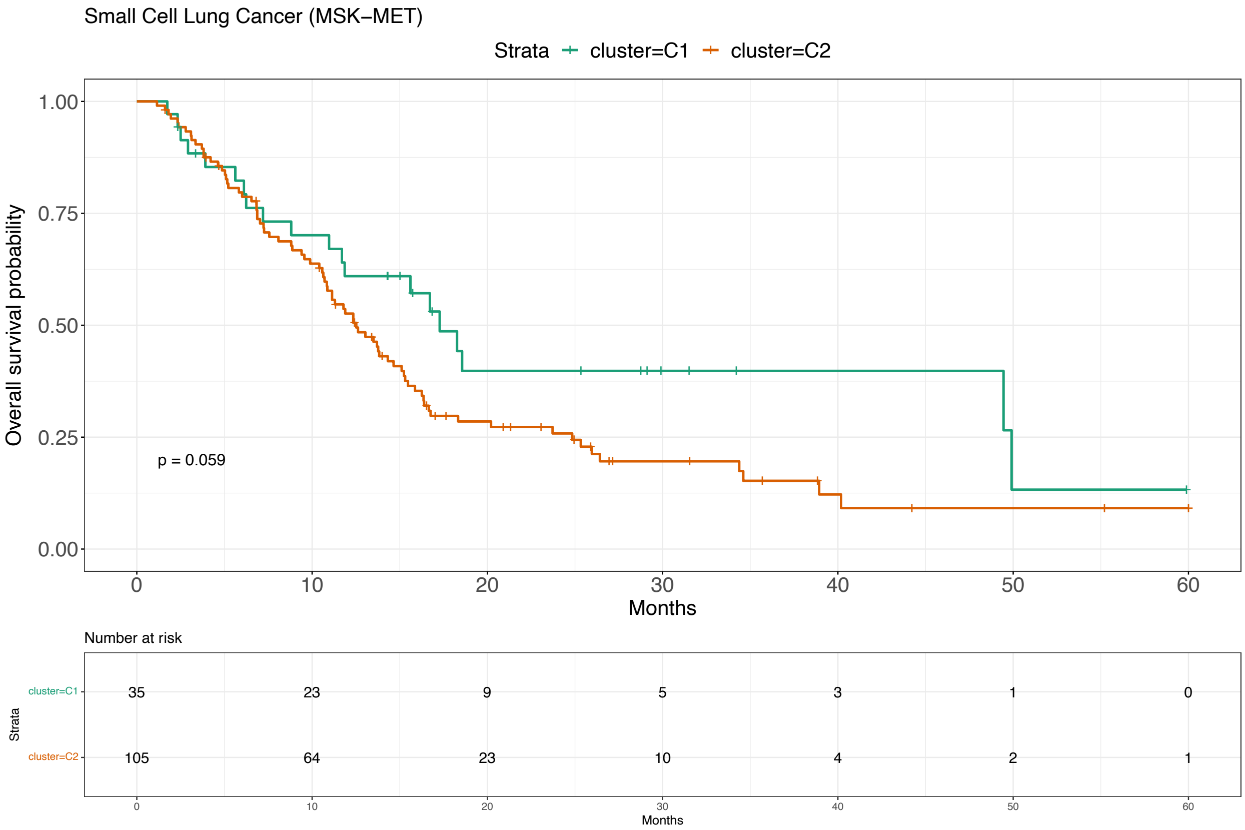

Figure 75: Survival analysis for Small-Cell Lung Cancer (MSK-MET) [21].

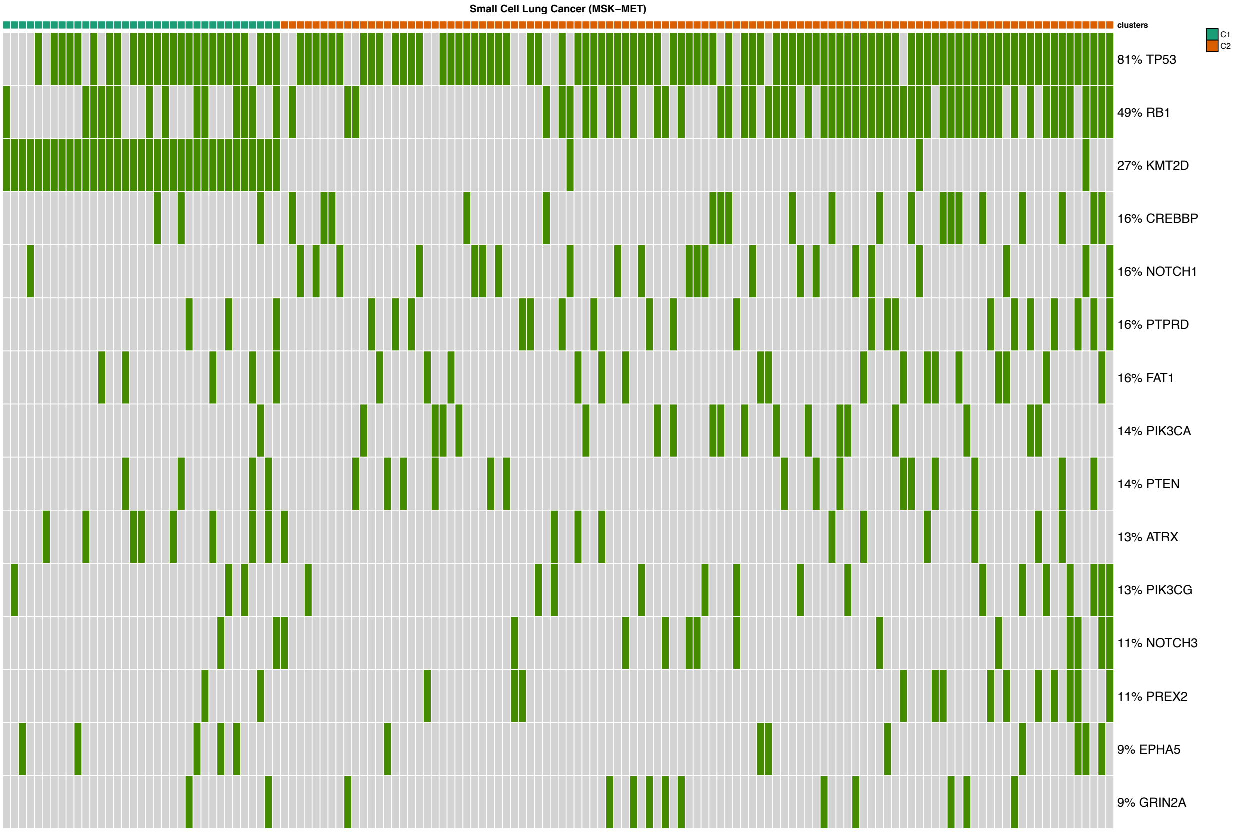

Figure 76: Mutational profile for Small-Cell Lung Cancer (MSK-MET) [21].

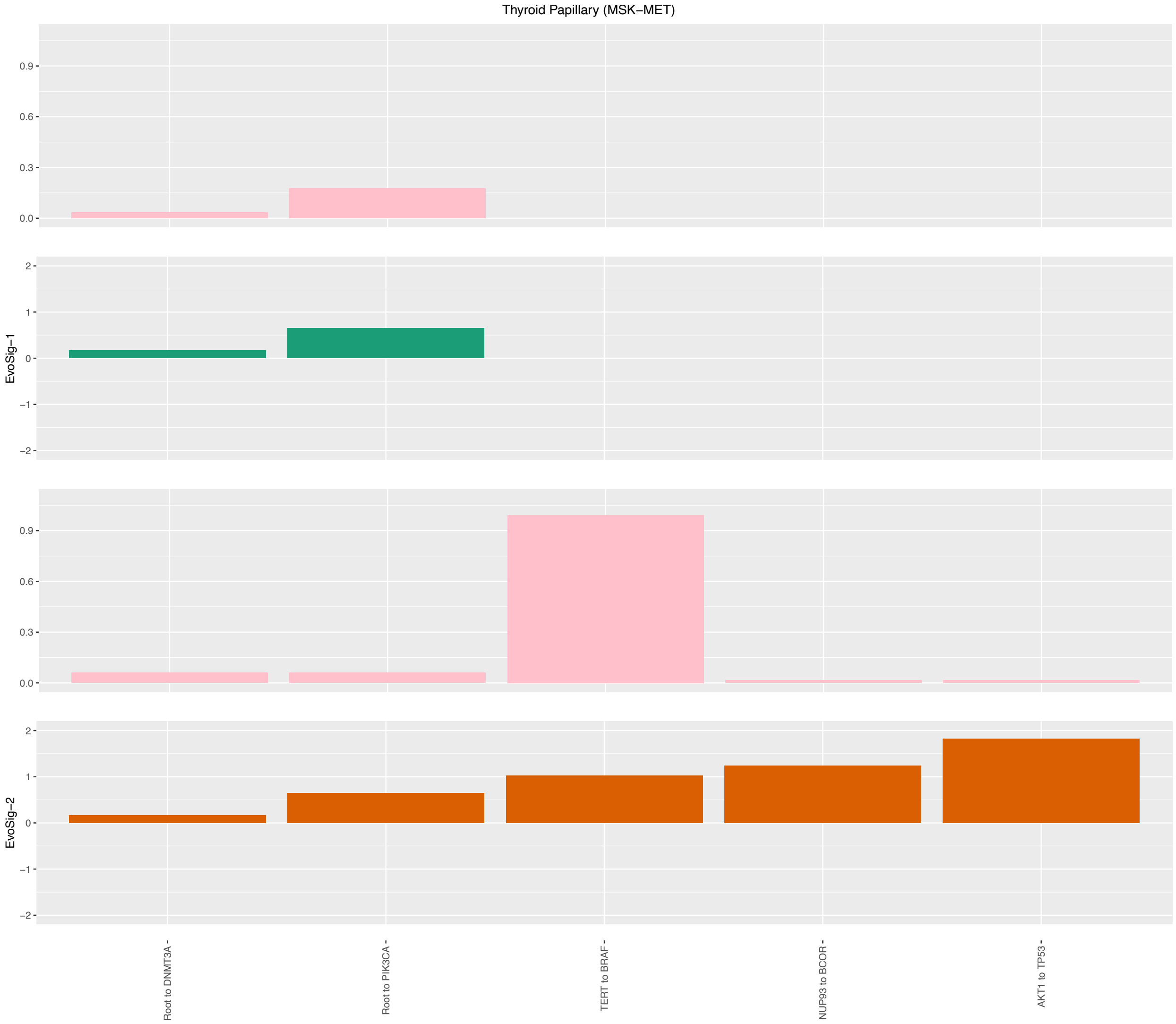

Figure 77: Evolutionary Signatures for Thyroid Papillary (MSK-MET) [21].

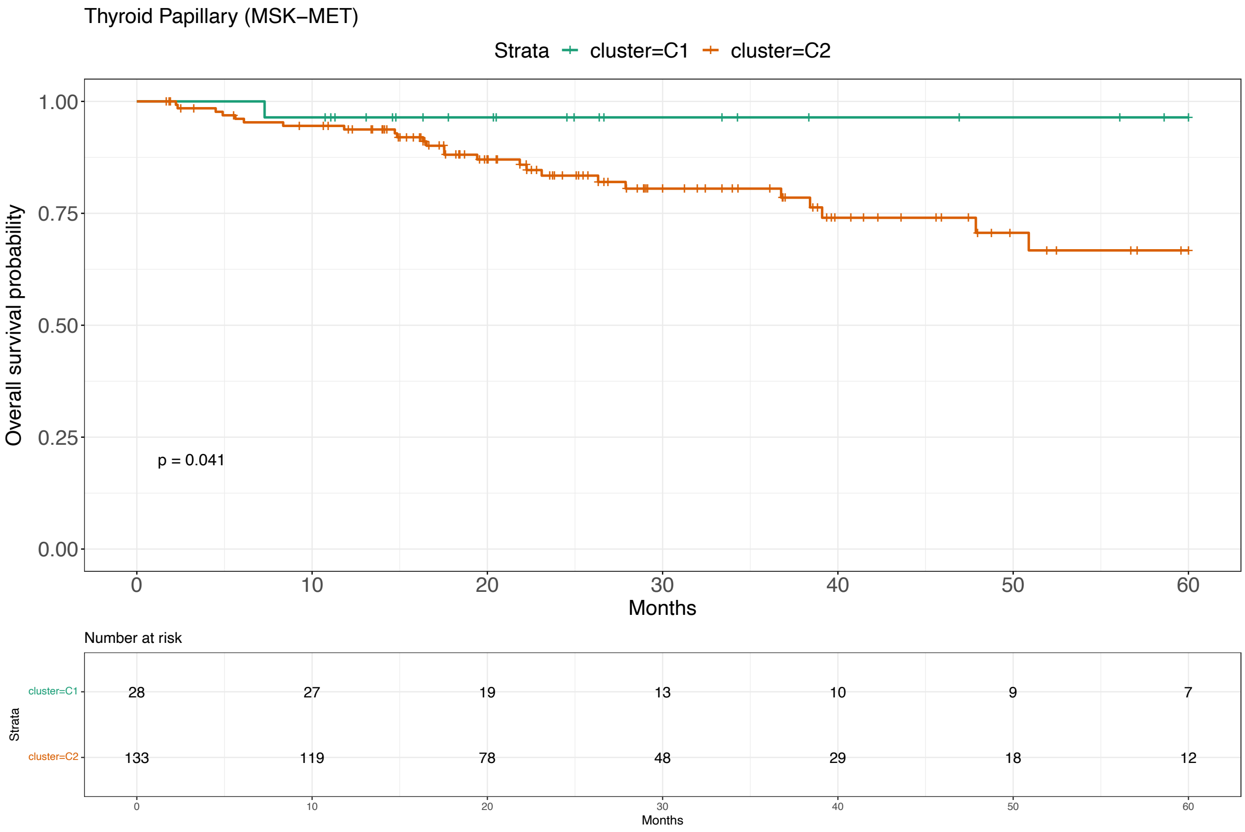

Figure 78: Survival analysis for Thyroid Papillary (MSK-MET) [21].

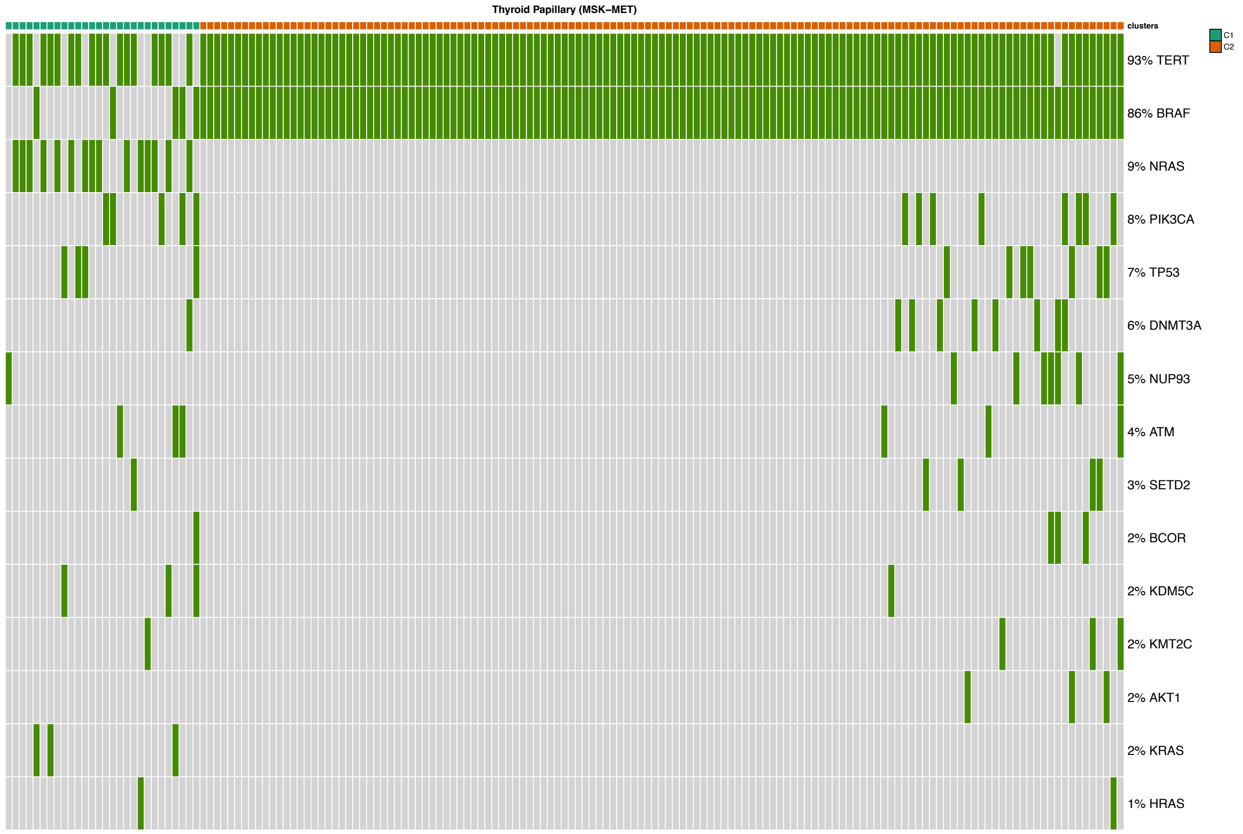

Figure 79: Mutational profile for Thyroid Papillary (MSK-MET) [21].

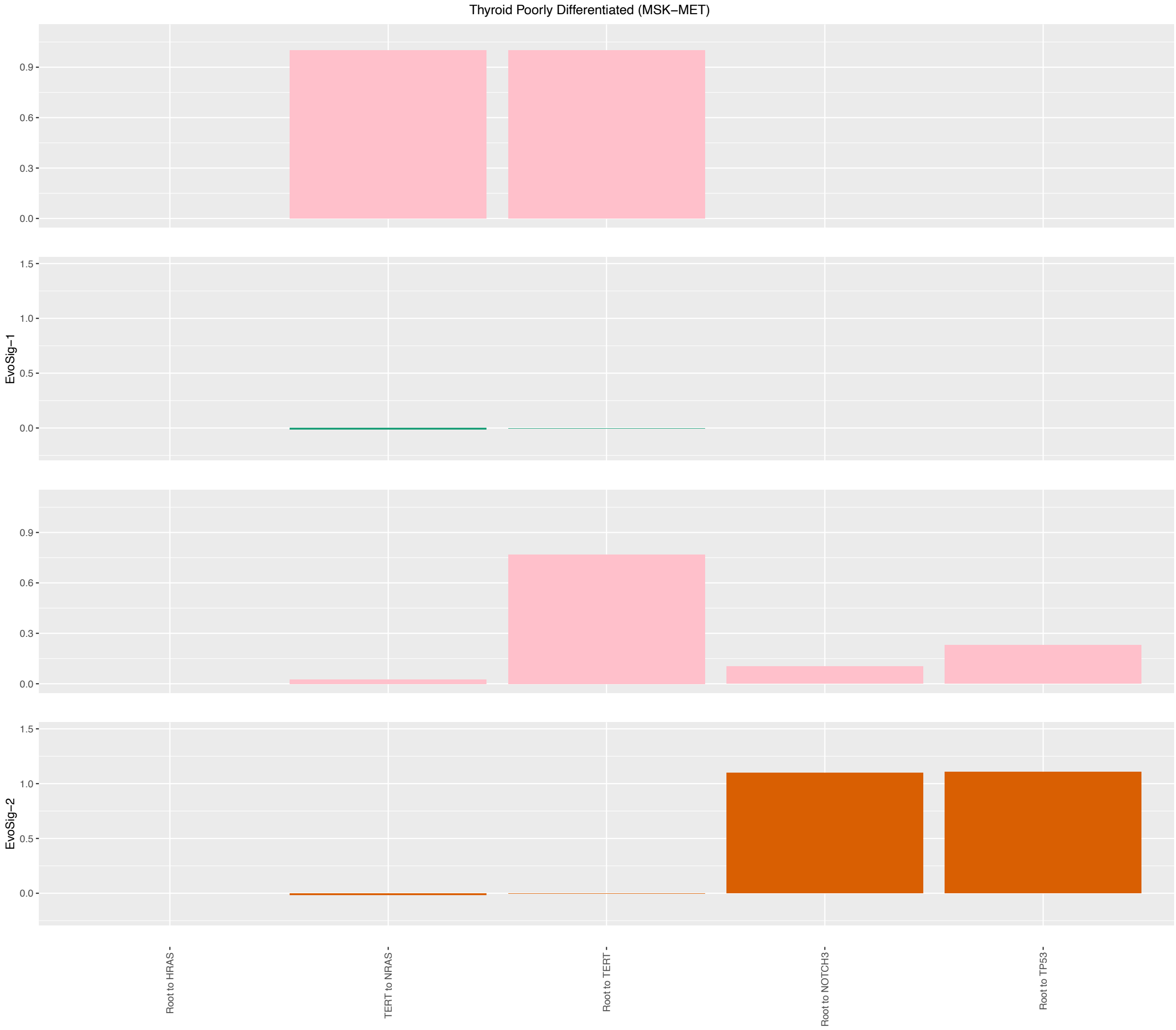

Figure 80: Evolutionary Signatures for Thyroid Poorly Differentiated (MSK-MET) [21].

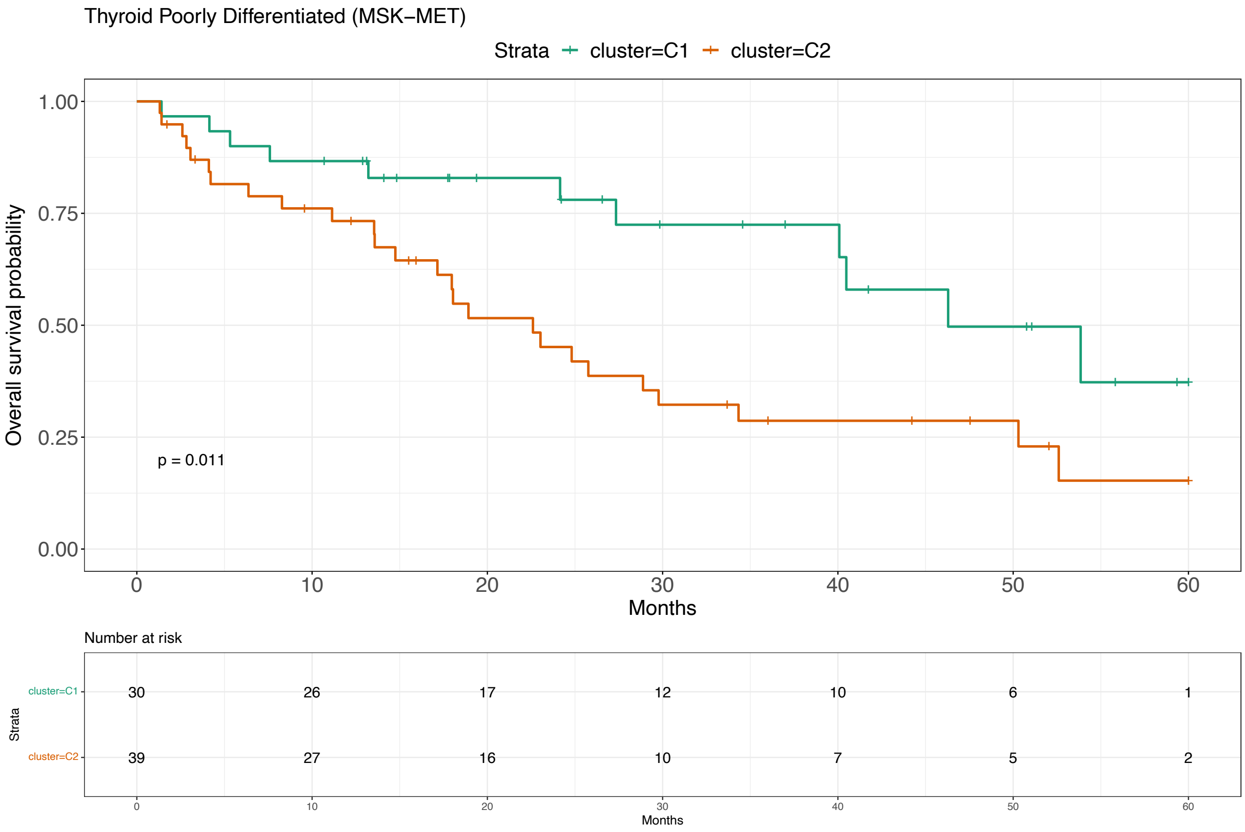

Figure 81: Survival analysis for Thyroid Poorly Differentiated (MSK-MET) [21].

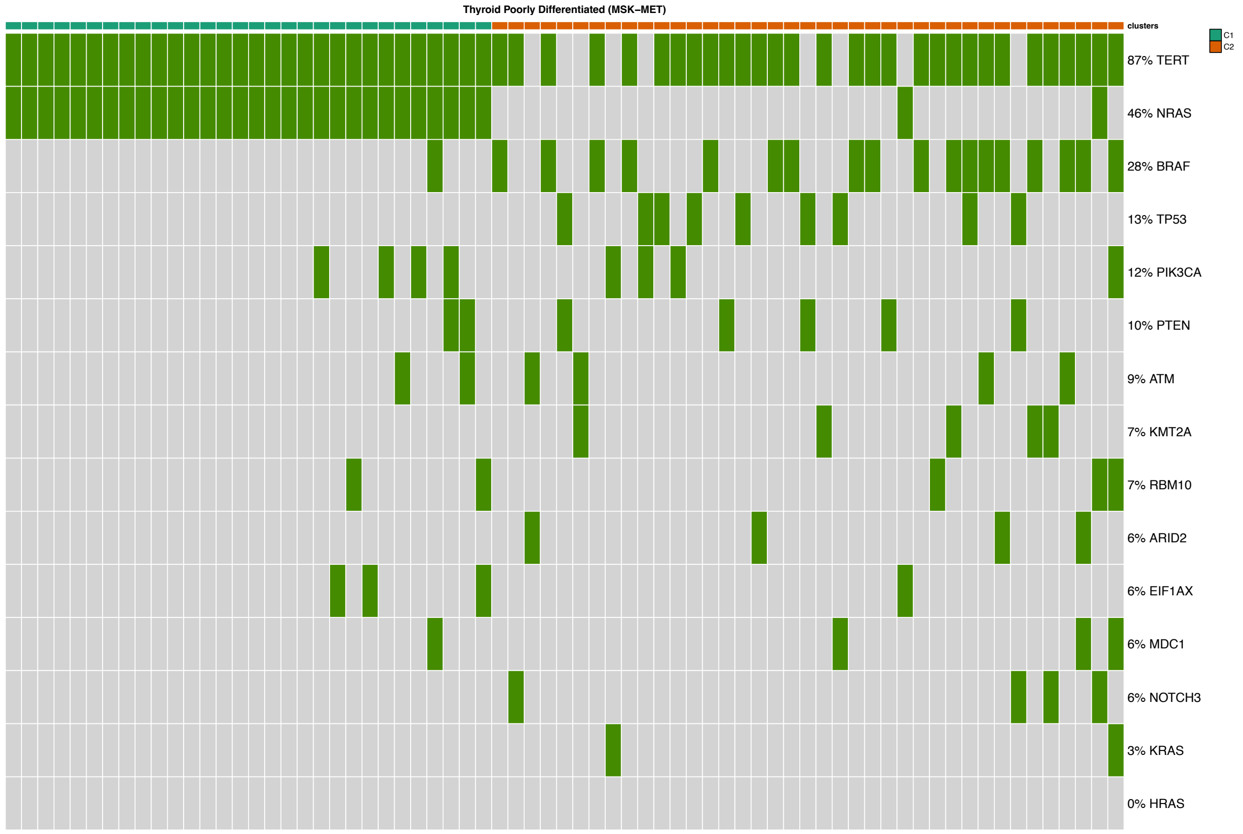

Figure 82: Mutational profile for Thyroid Poorly Differentiated (MSK-MET) [21].

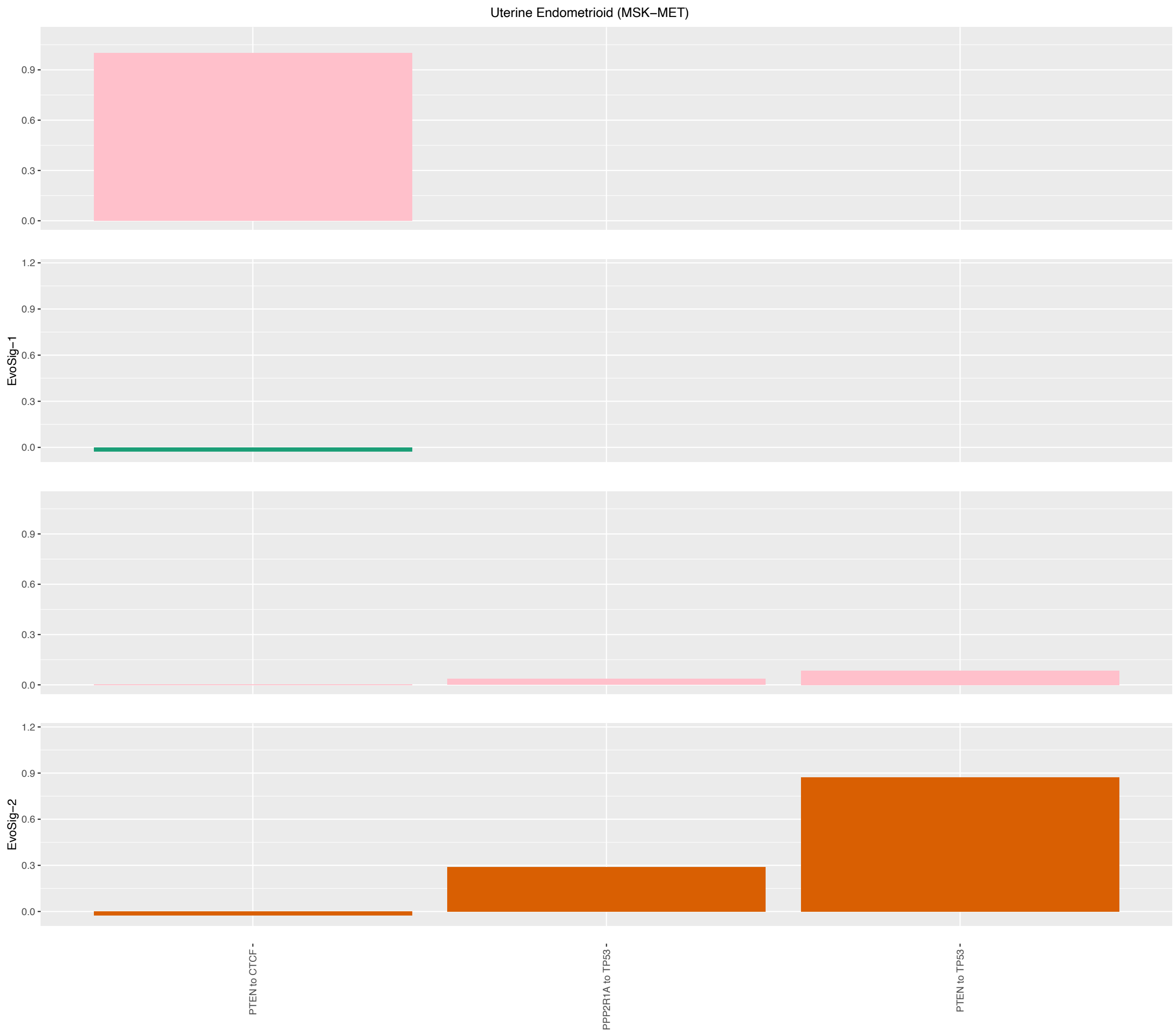

Figure 83: Evolutionary Signatures for Uterine Endometrioid (MSK-MET) [21].

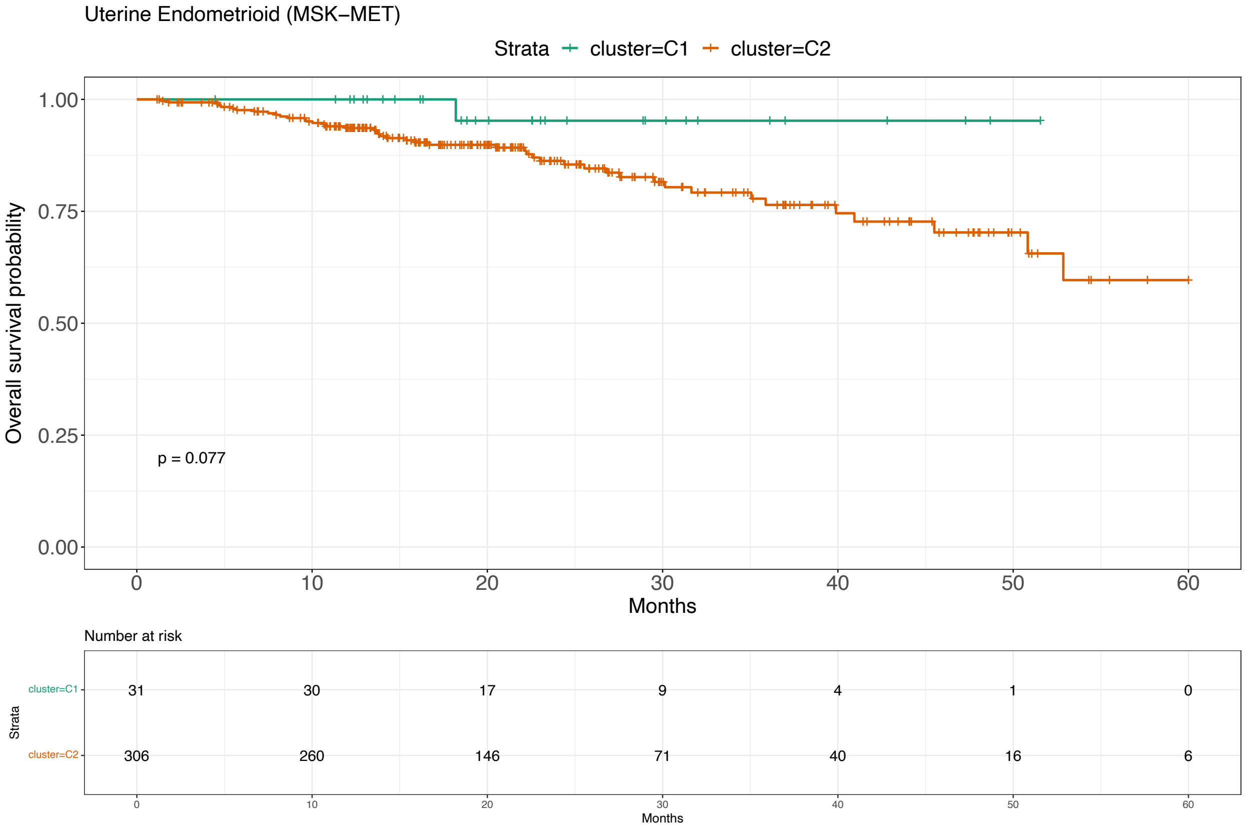

Figure 84: Survival analysis for Uterine Endometrioid (MSK-MET) [21].

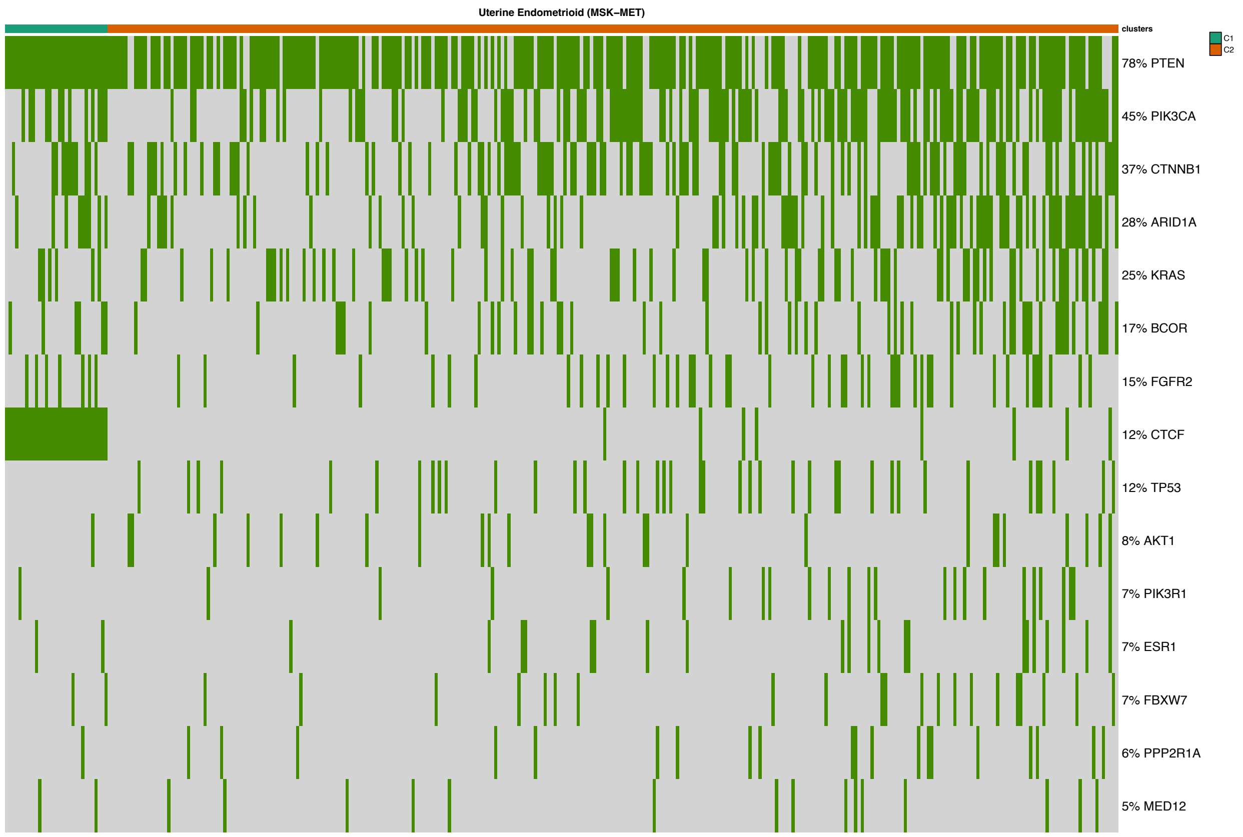

Figure 85: Mutational profile for Uterine Endometrioid (MSK-MET) [21].

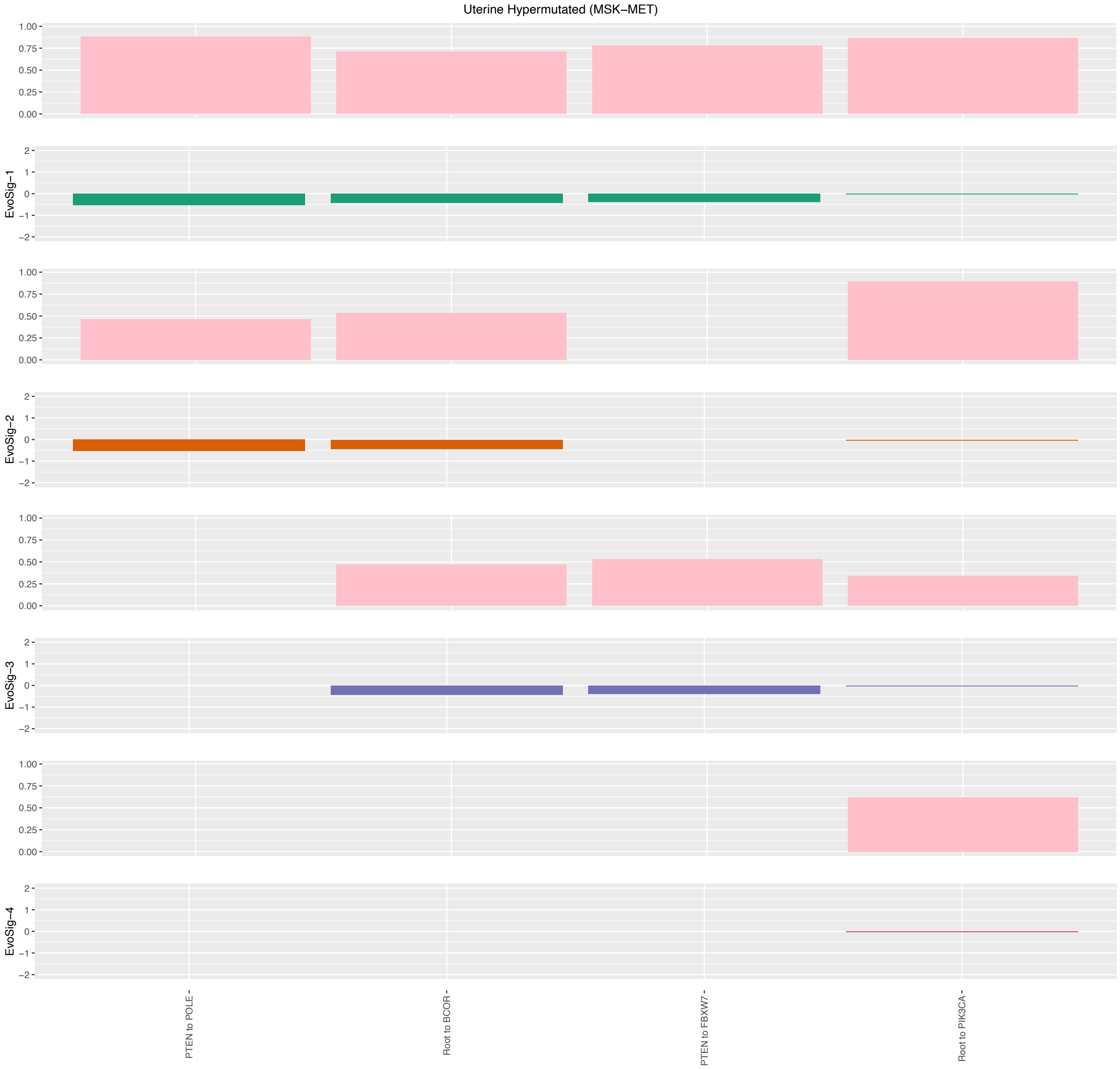

Figure 86: Evolutionary Signatures for Uterine Hypermutated (MSK-MET) [21].

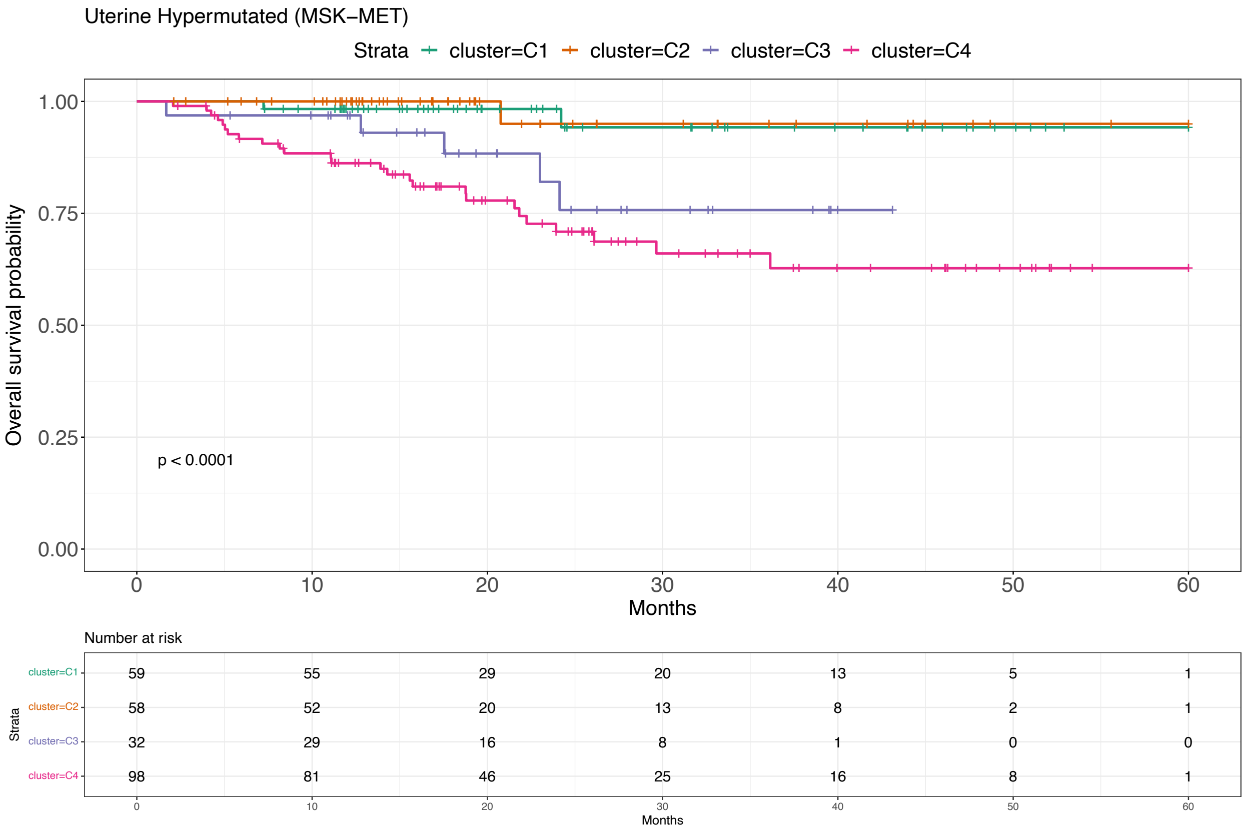

Figure 87: Survival analysis for Uterine Hypermutated (MSK-MET) [21].

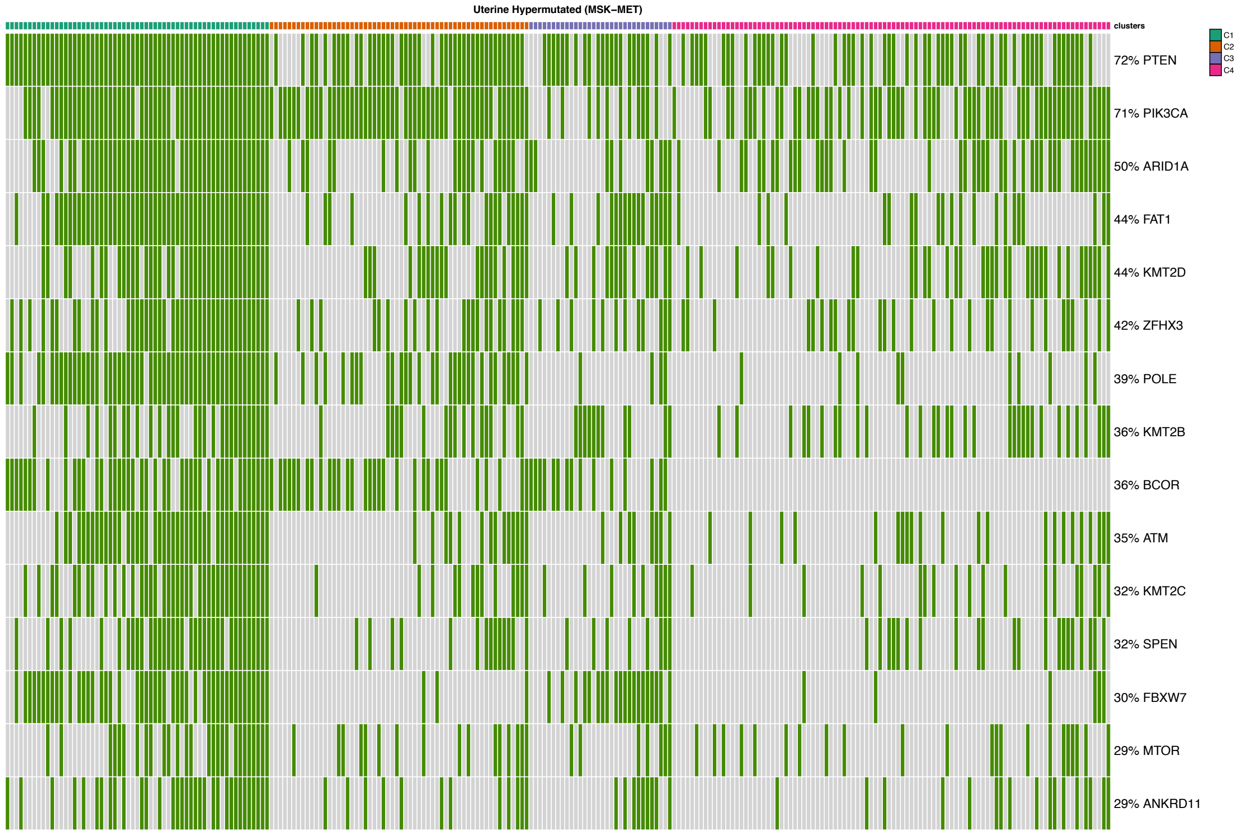

Figure 88: Mutational profile for Uterine Hypermutated (MSK-MET) [21].

### 5 **ASCETIC** models of Myeloid Malignancies and Early-stage Non-small cell lung cancer

Myeloid Malignancies ([24])

| | | |
|---|---|---|
| *CALR* | *ASXL1* | 0.93 |
| *CALR* | *ATM* | 0.65 |
| *SF3B1* | *CHEK2* | 0.54 |
| *NPM1* | *FLT3* | 0.68 |
| *DNMT3A* | *NRAS* | 0.96 |
| *JAK2* | *NRAS* | 0.74 |
| *JAK2* | *RUNX1* | 0.50 |
| *IDH1* | *TET2* | 0.97 |
| *IDH2* | *TET2* | 0.66 |
| *KIT* | *WT1* | 0.69 |

Early-stage Non-small cell lung cancer ([20])

| | | |
|---|---|---|
| *CCND1* | *ARID1B* | 0.88 |
| *SPEN* | *ARID1B* | 0.57 |
| *KEAP1* | *ARID2* | 0.96 |
| *CDKN2A* | *CYLD* | 0.83 |
| *ASXL1* | *DNM2* | 0.57 |
| *EGFR* | *EP300* | 0.62 |
| *FBXW7* | *EP300* | 0.86 |
| *TP53* | *FAT1* | 0.94 |
| *TP53* | *KRAS* | 0.62 |
| *SPEN* | *LATS1* | 0.71 |
| *KEAP1* | *NCOR1* | 0.64 |
| *EP300* | *NF1* | 0.61 |
| *PIK3CA* | *NFE2L2* | 0.68 |
| *SOX2* | *NFE2L2* | 0.52 |
| *DICER1* | *PLXNB2* | 0.77 |
| *ATM* | *POLE* | 0.60 |
| *DICER1* | *PTPRC* | 0.51 |
| *KMT2D* | *PTPRC* | 0.54 |
| *BRAF* | *TERT* | 0.73 |
| *KMT2D* | *UBR5* | 0.61 |
| *SOX2* | *UBR5* | 0.79 |
| *KEAP1* | *WRN* | 0.68 |

Table 1: Results by ASCETIC for (top) single-cell data for a set of myeloid malignancies [24], including clonal haematopoiesis, myeloproliferative neoplasms, and acute myeloid leukemia, and (bottom) multi-region sequencing data from patients with early-stage non-small cell lung cancer obtained from the TRACERx project [20]. First and second columns report parent and child gene. Third column reports the cross validation score for the relative relation.

### 6 Validation of ASCETIC models on unseen datasets

| Parent (P) | Child (C) | P → C | C → P | No relation |
|---|---|---|---|---|
| *IDH1* | *DNMT3A* | 2 | 0 | 2 |
| *JAK2* | *DNMT3A* | 0 | 0 | 0 |
| *IDH1* | *EZH2* | 0 | 0 | 0 |
| *NPM1* | *FLT3* | 5 | 2 | 0 |
| *U2AF1* | *FLT3* | 4 | 0 | 0 |
| *JAK2* | *KRAS* | 1 | 0 | 0 |
| *ASXL1* | *NRAS* | 0 | 0 | 0 |
| *DNMT3A* | *NRAS* | 0 | 2 | 0 |
| *JAK2* | *NRAS* | 0 | 0 | 0 |
| *ASXL1* | *PTPN11* | 0 | 0 | 1 |
| *IDH1* | *PTPN11* | 11 | 2 | 1 |
| *JAK2* | *RUNX1* | 0 | 0 | 0 |
| *DNMT3A* | *SETBP1* | 0 | 0 | 0 |
| *IDH1* | *TET2* | 1 | 0 | 0 |
| *IDH2* | *TET2* | 0 | 0 | 0 |
| *DNMT3A* | *WT1* | 0 | 2 | 1 |
| *KIT* | *WT1* | 1 | 0 | 0 |

Table 2: Validation on unseen data from Morita and colleagues [25]. First and second columns report parent and child gene. Third and fourth columns report number of samples were the phylogenetic trees by Morita and colleagues [25] show respectively consistent and inconsistent evolution dynamics with respect to the arcs inferred by ASCETIC. Fifth columns report the number of samples where the phylogenetic analysis provides inconclusive evidence regarding the ordering of accumulation between the considered pair of genes.

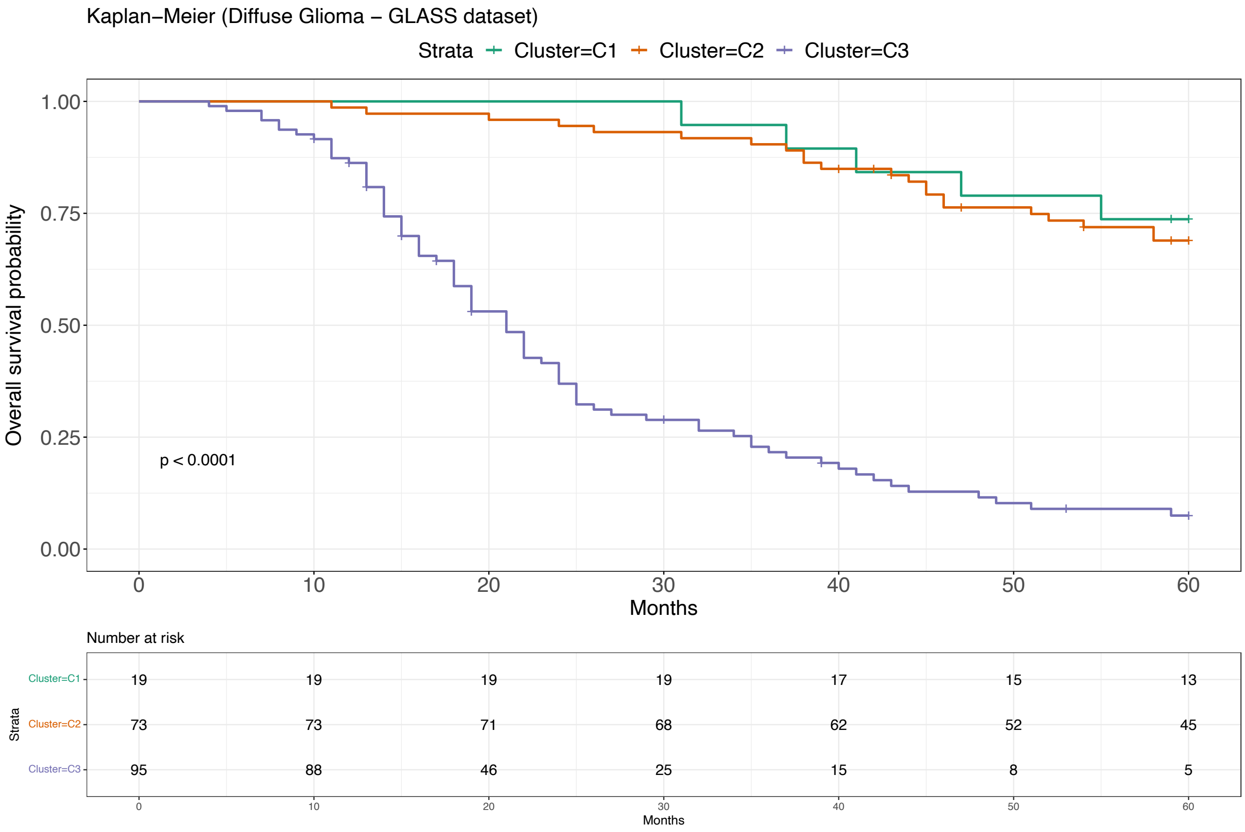

Figure 89: Survival analysis for Diffuse Glioma - Subtypes Validation (GLASS) [26].

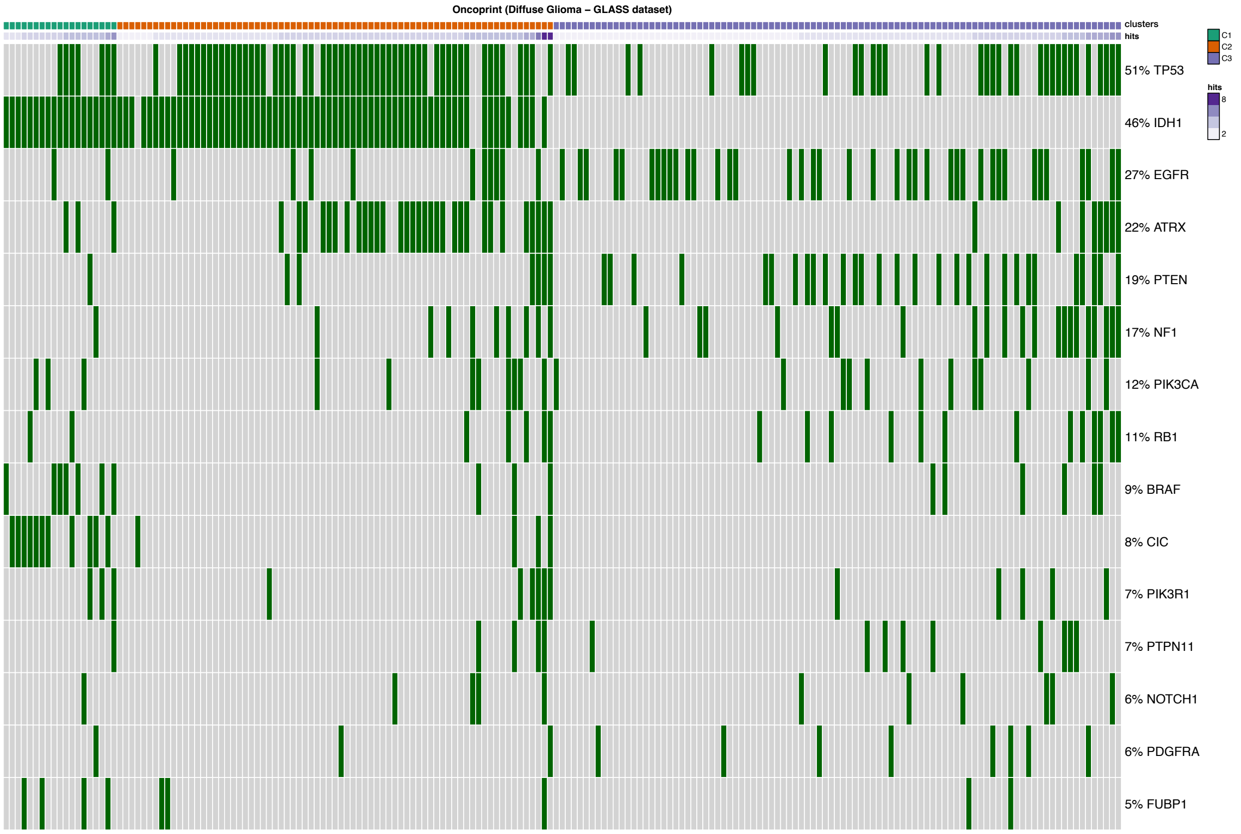

Figure 90: Oncoprint for Diffuse Glioma - Subtypes Validation (GLASS) [26].

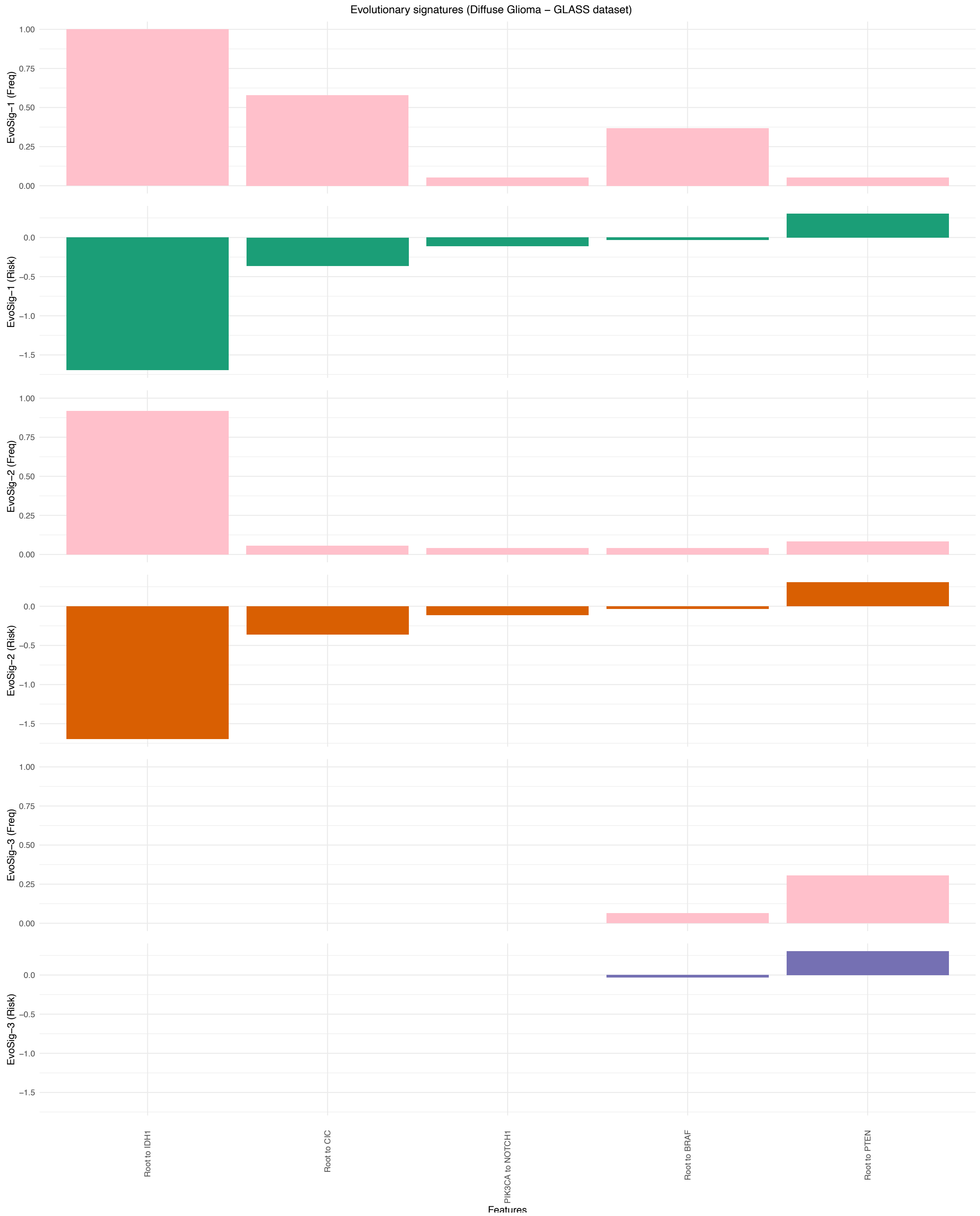

Figure 91: Evolutionary Signatures for Diffuse Glioma - Subtypes Validation (GLASS) [26].

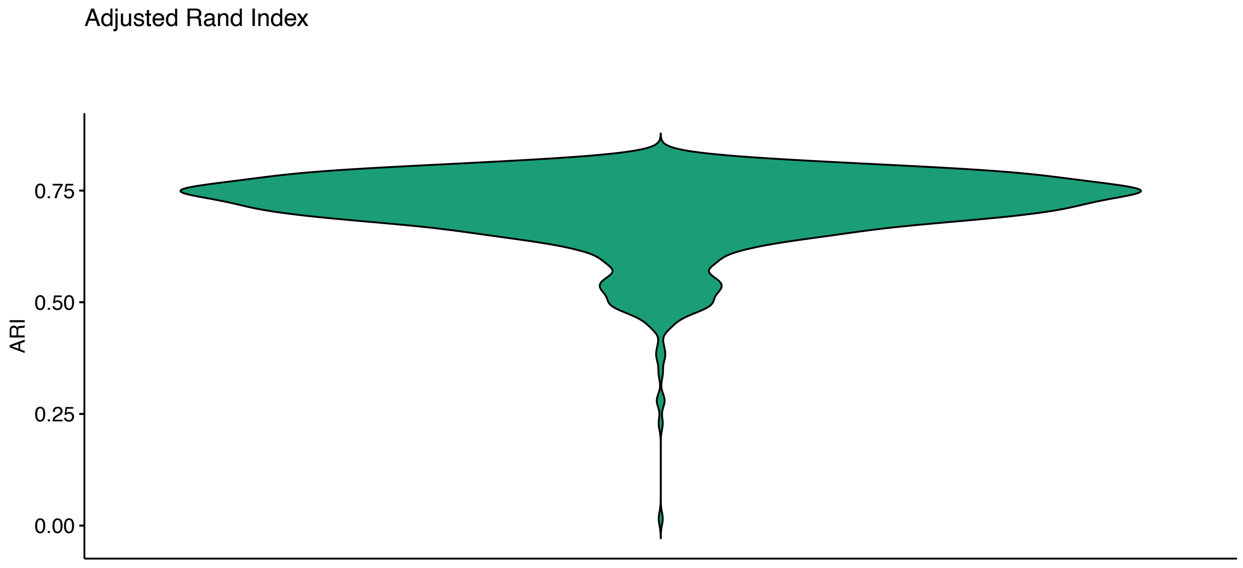

Figure 92: Results of simulations to evaluate the robustness of ASCETIC's stratification considering the Diffuse Glioma dataset from the GLASS study [26].

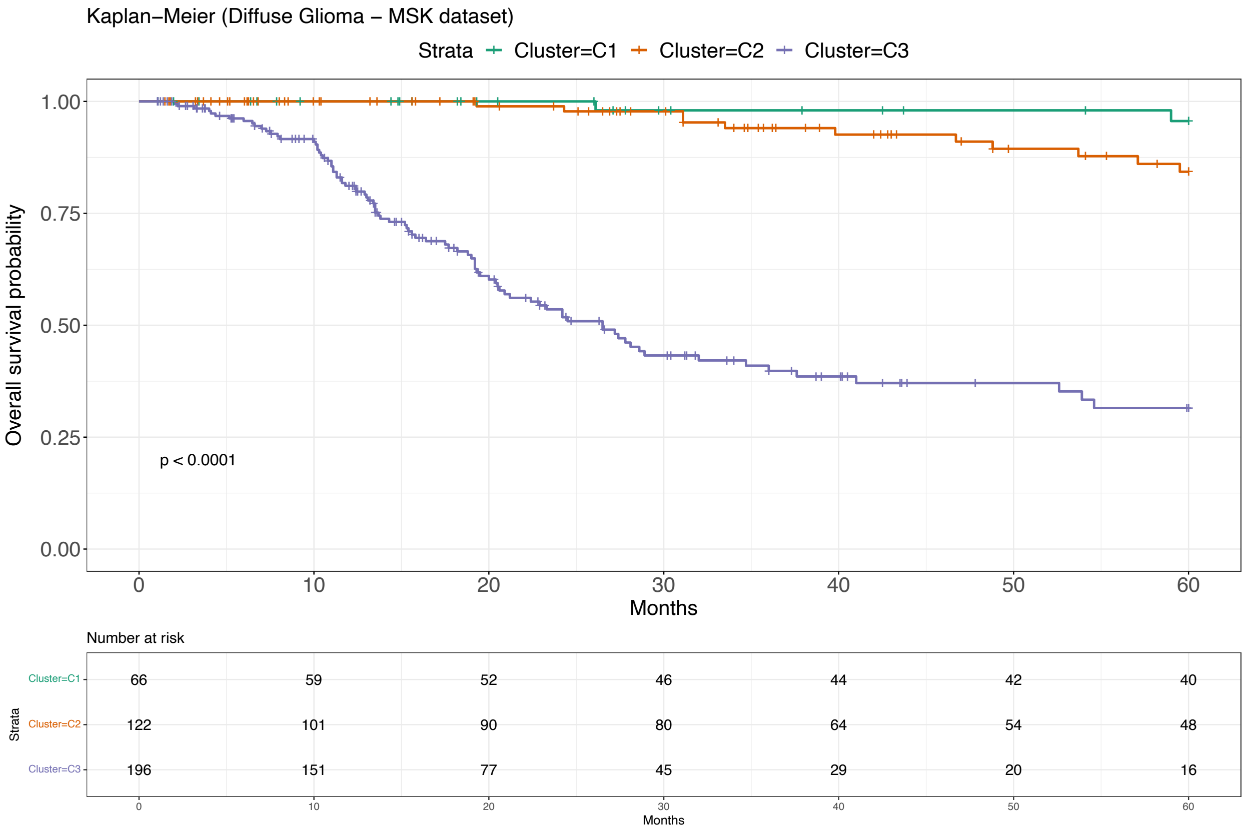

Figure 93: Survival analysis for Diffuse Glioma - Subtypes Validation (MSK) [27].

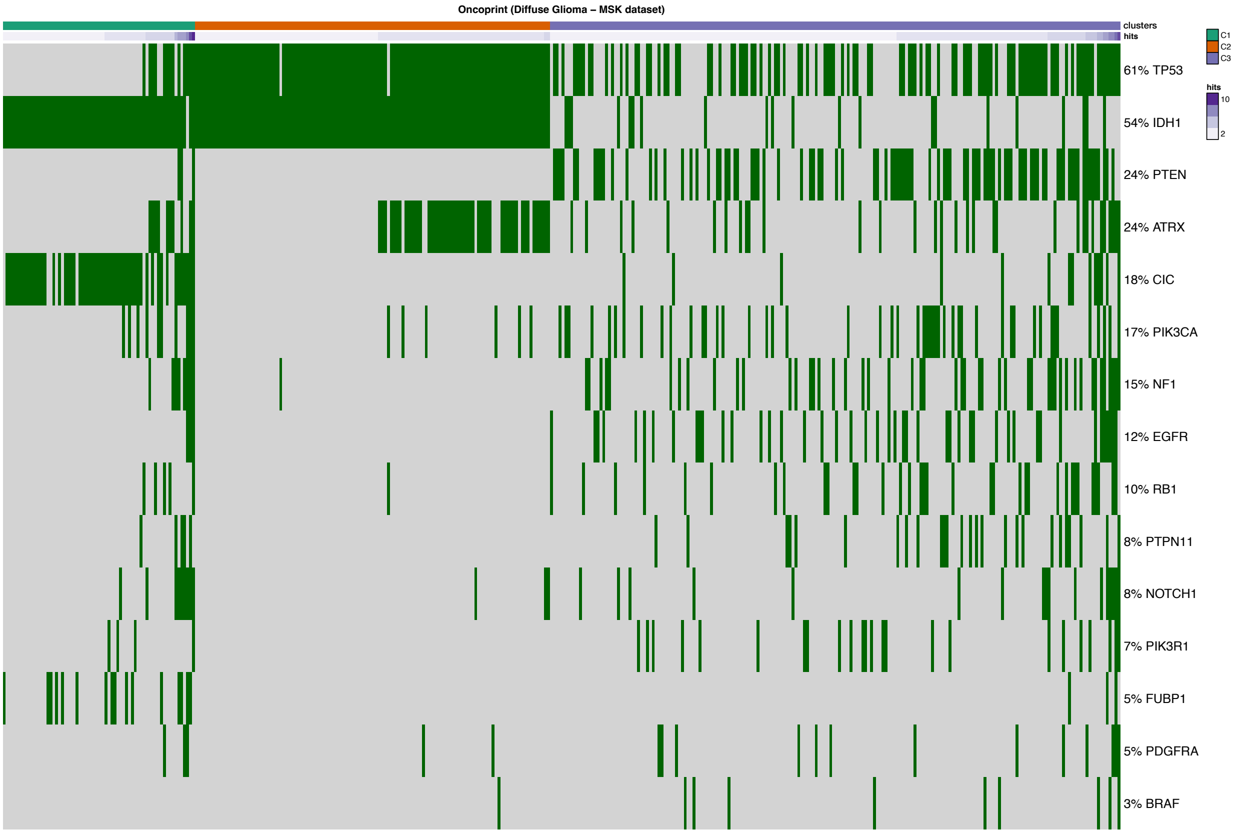

Figure 94: Oncoprint for Diffuse Glioma - Subtypes Validation (MSK) [27].

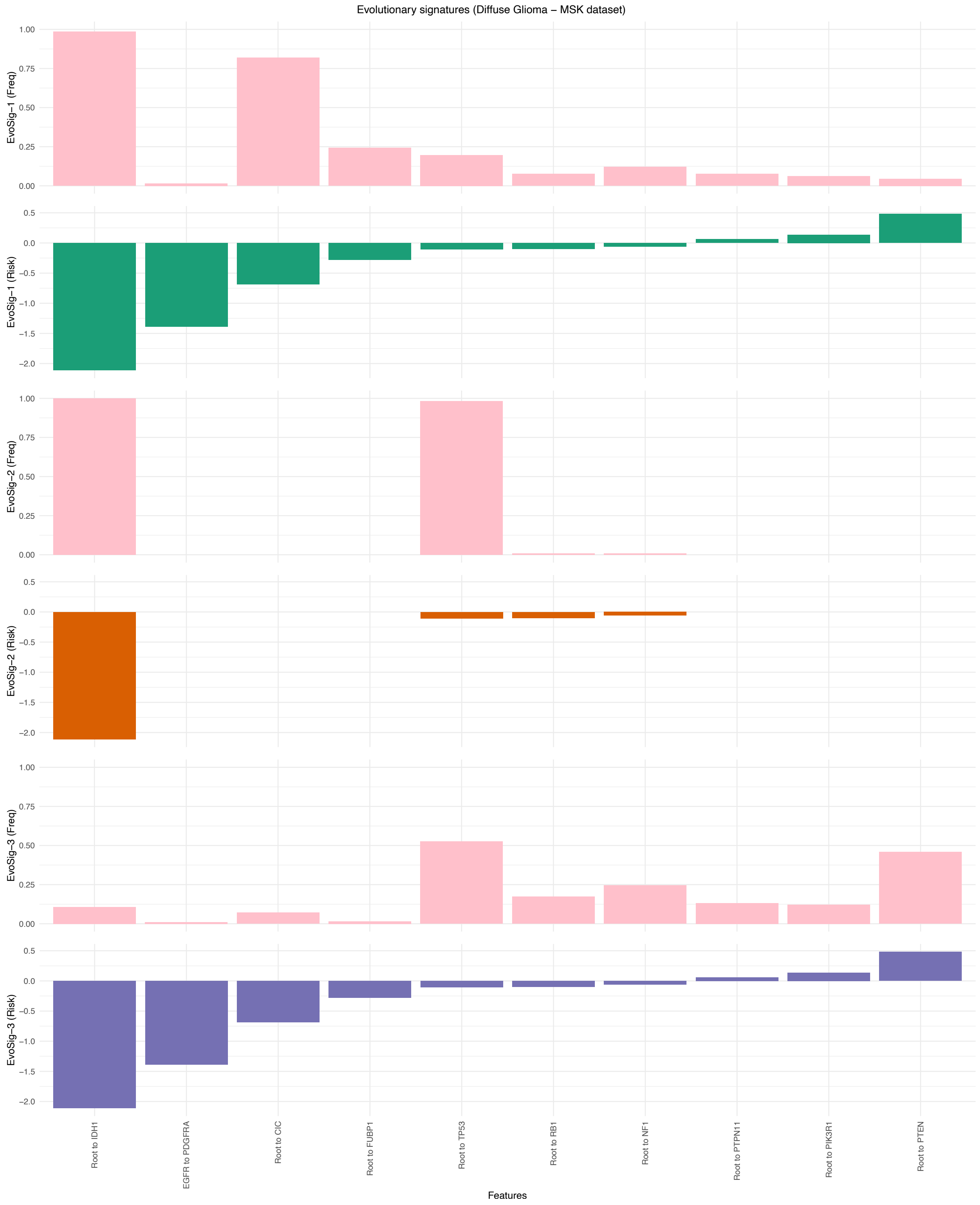

Figure 95: Evolutionary Signatures for Diffuse Glioma - Subtypes Validation (MSK) [27].

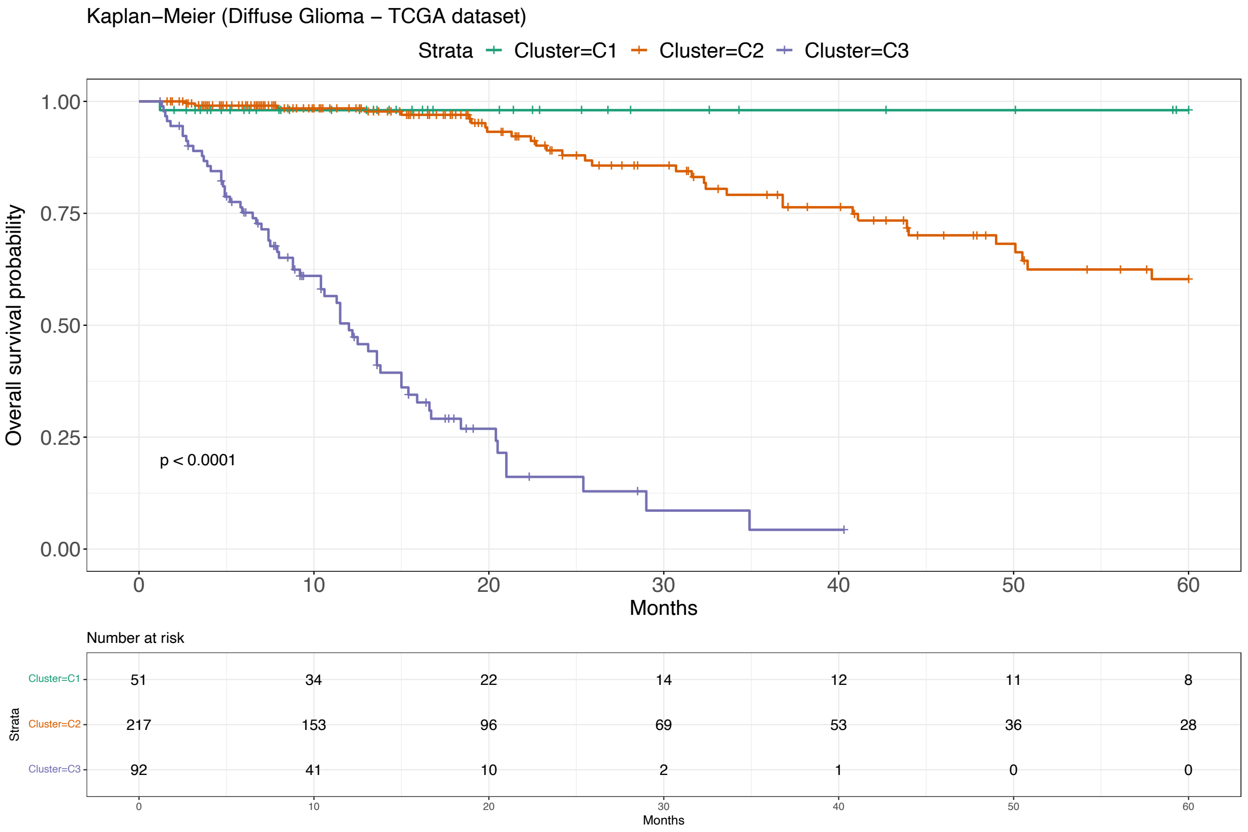

Figure 96: Survival analysis for Diffuse Glioma - Subtypes Validation (TCGA) [28].

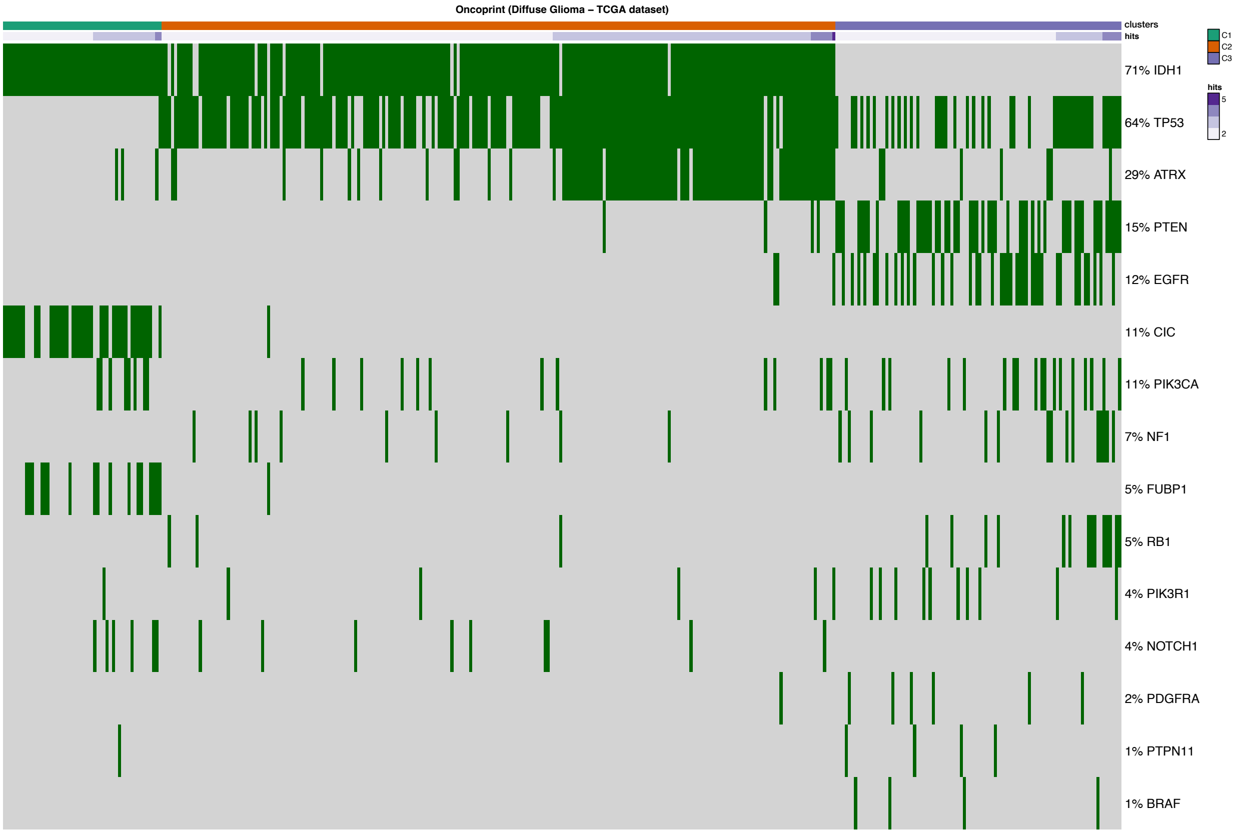

Figure 97: Oncoprint for Diffuse Glioma - Subtypes Validation (TCGA) [28].

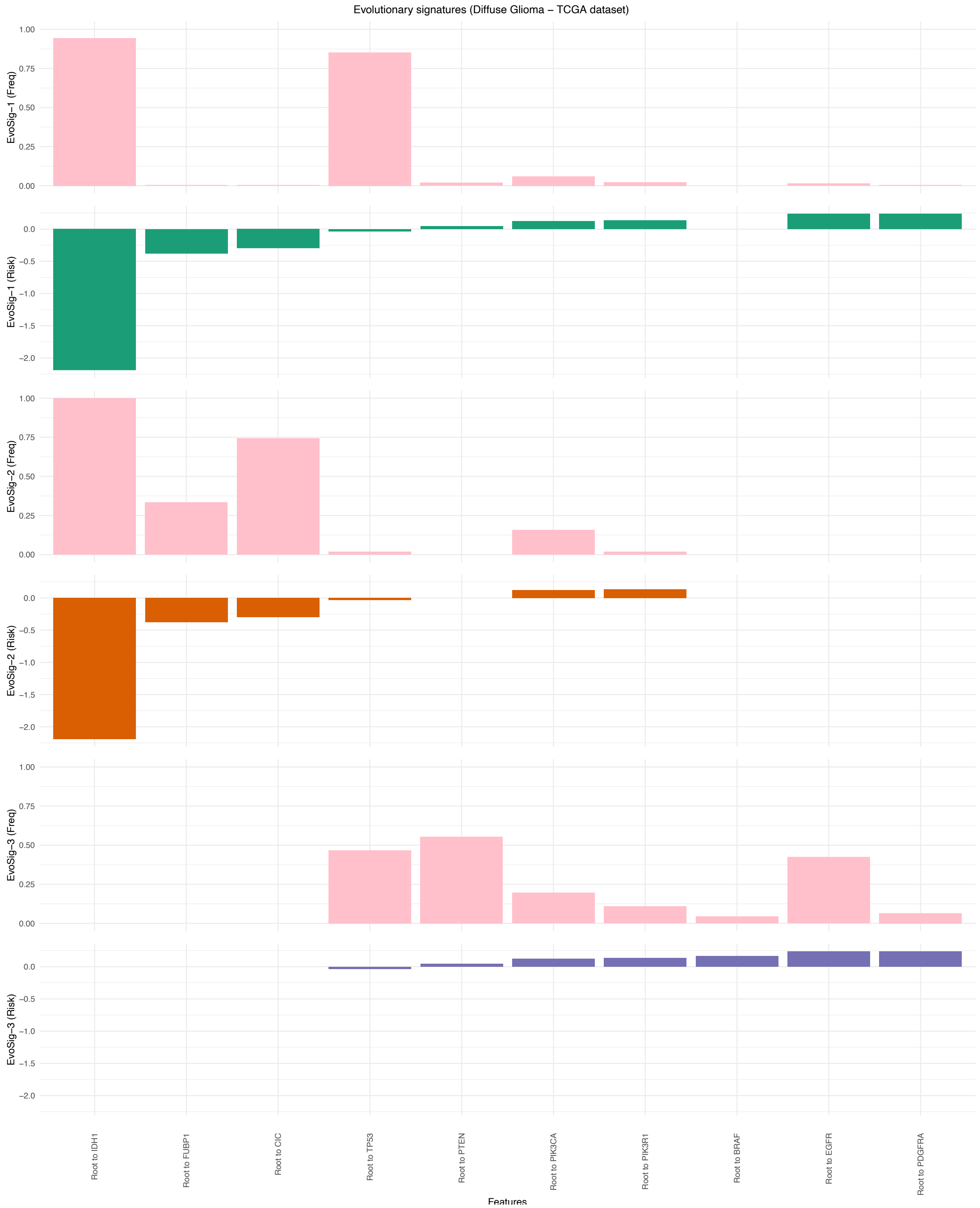

Figure 98: Evolutionary Signatures for Diffuse Glioma - Subtypes Validation (TCGA) [27].

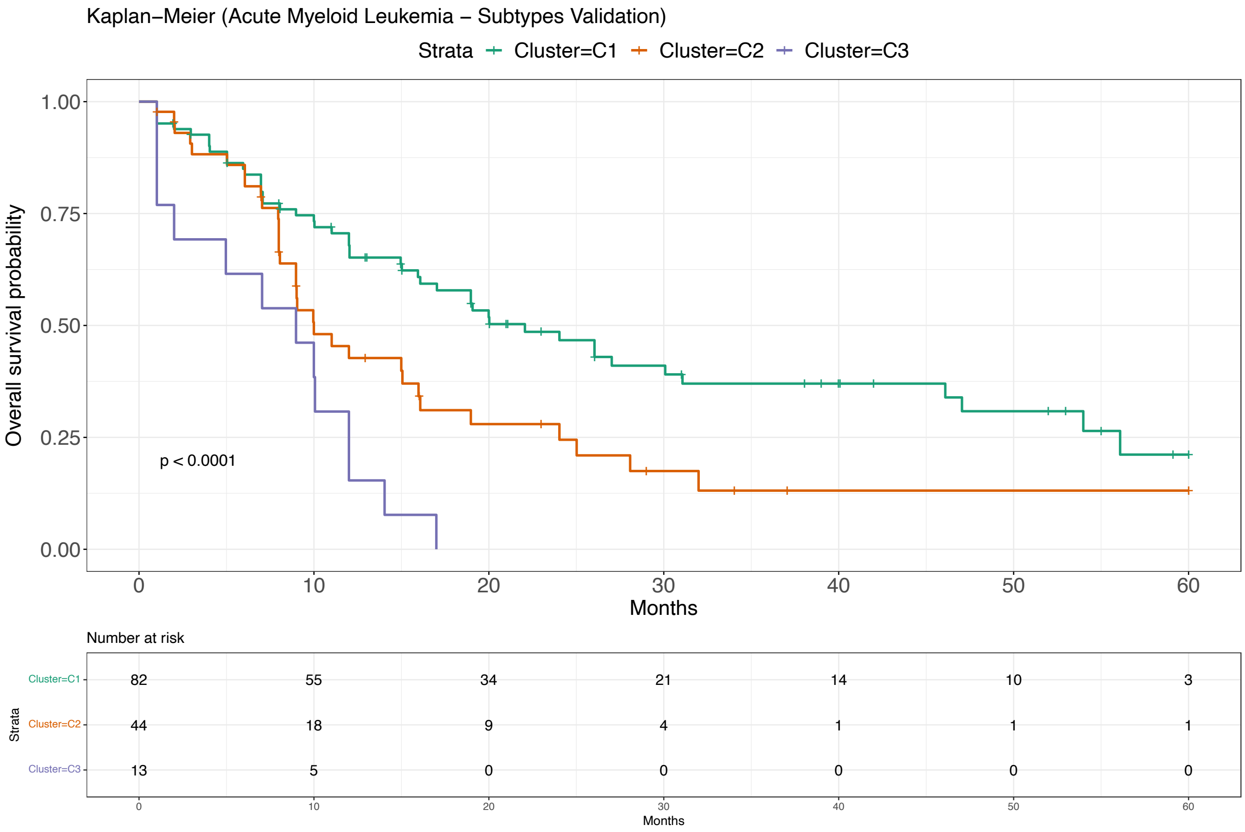

Figure 99: Survival analysis for Acute Myeloid Leukemia - Subtypes Validation (Pan-Cancer Atlas) [23].

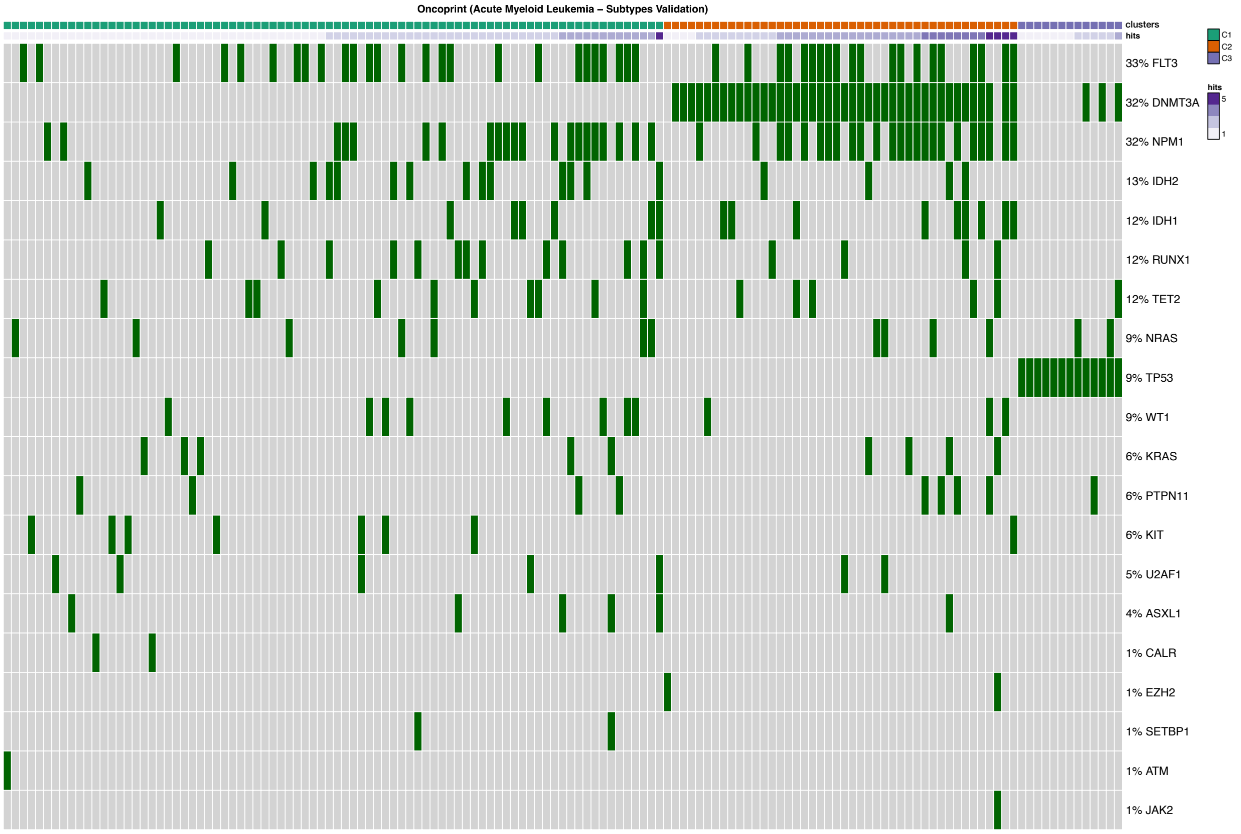

Figure 100: Oncoprint for Acute Myeloid Leukemia - Subtypes Validation (Pan-Cancer Atlas) [23].

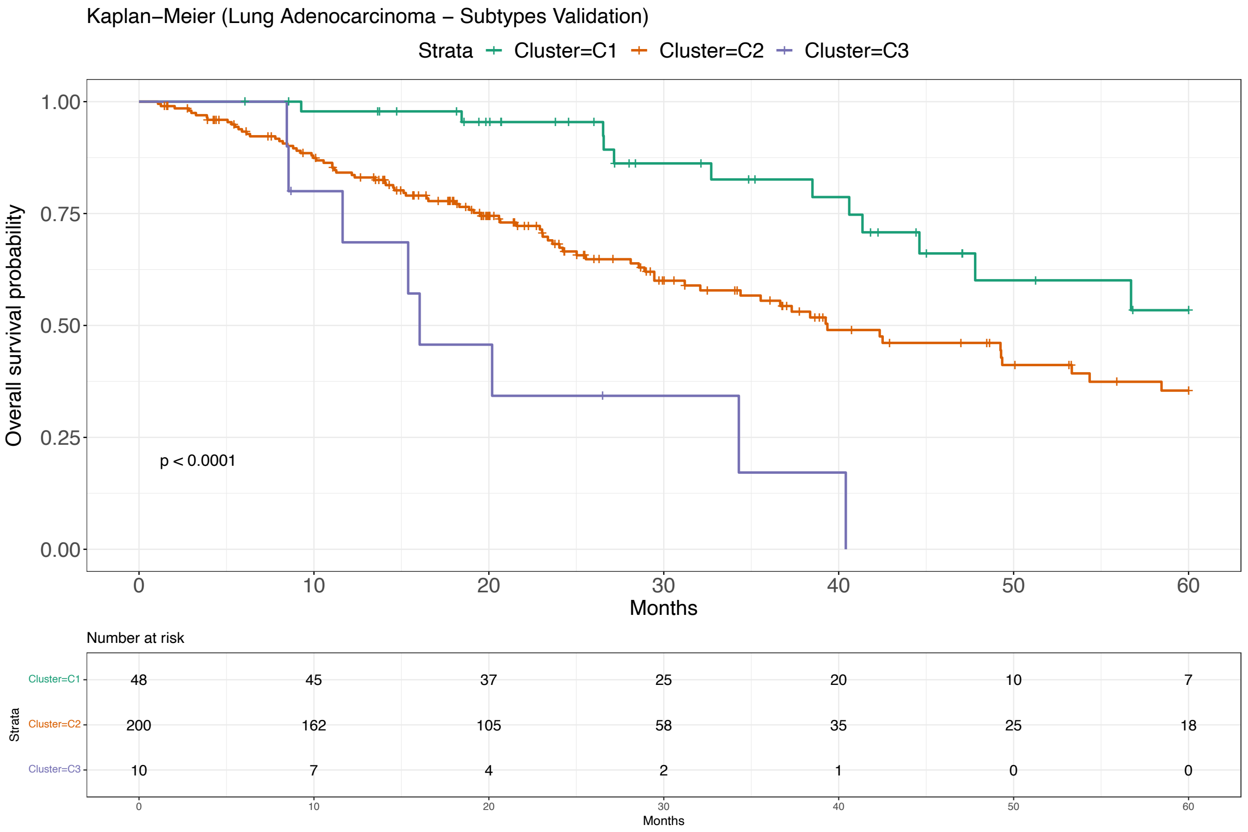

Figure 101: Survival analysis for Lung Adenocarcinoma - Subtypes Validation (Pan-Cancer Atlas) [23].

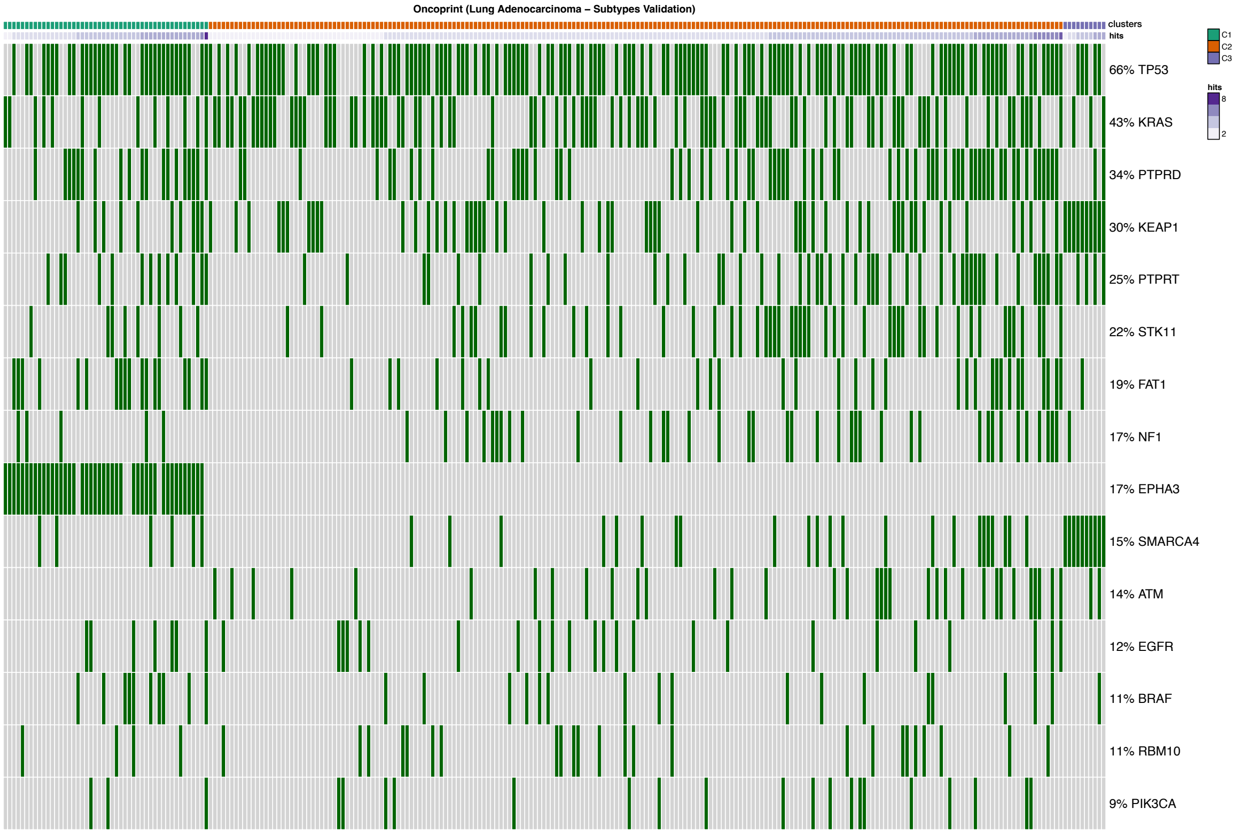

Figure 102: Oncoprint for Lung Adenocarcinoma - Subtypes Validation (Pan-Cancer Atlas) [23].

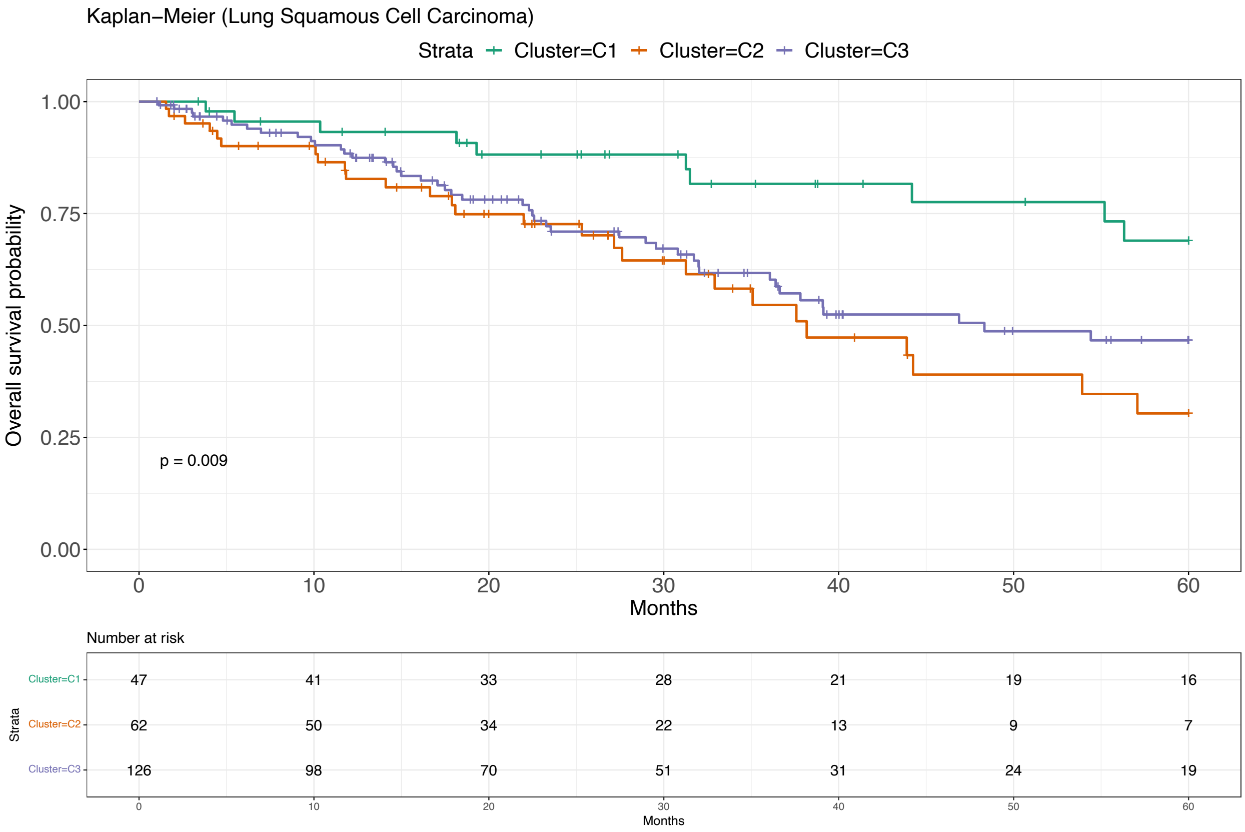

Figure 103: Survival analysis for Lung Squamous Cell Carcinoma - Subtypes Validation (Pan-Cancer Atlas) [23].

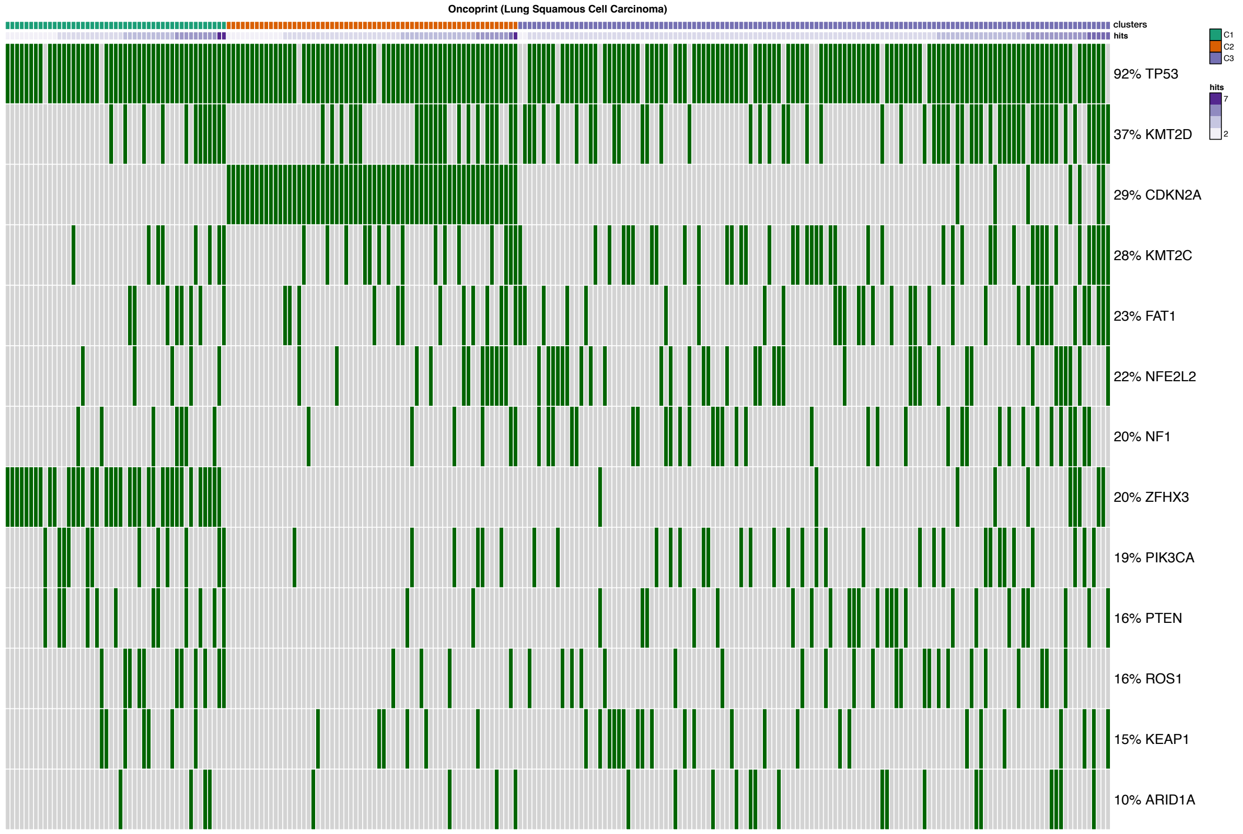

Figure 104: Oncoprint for Lung Squamous Cell Carcinoma - Subtypes Validation (Pan-Cancer Atlas) [23].

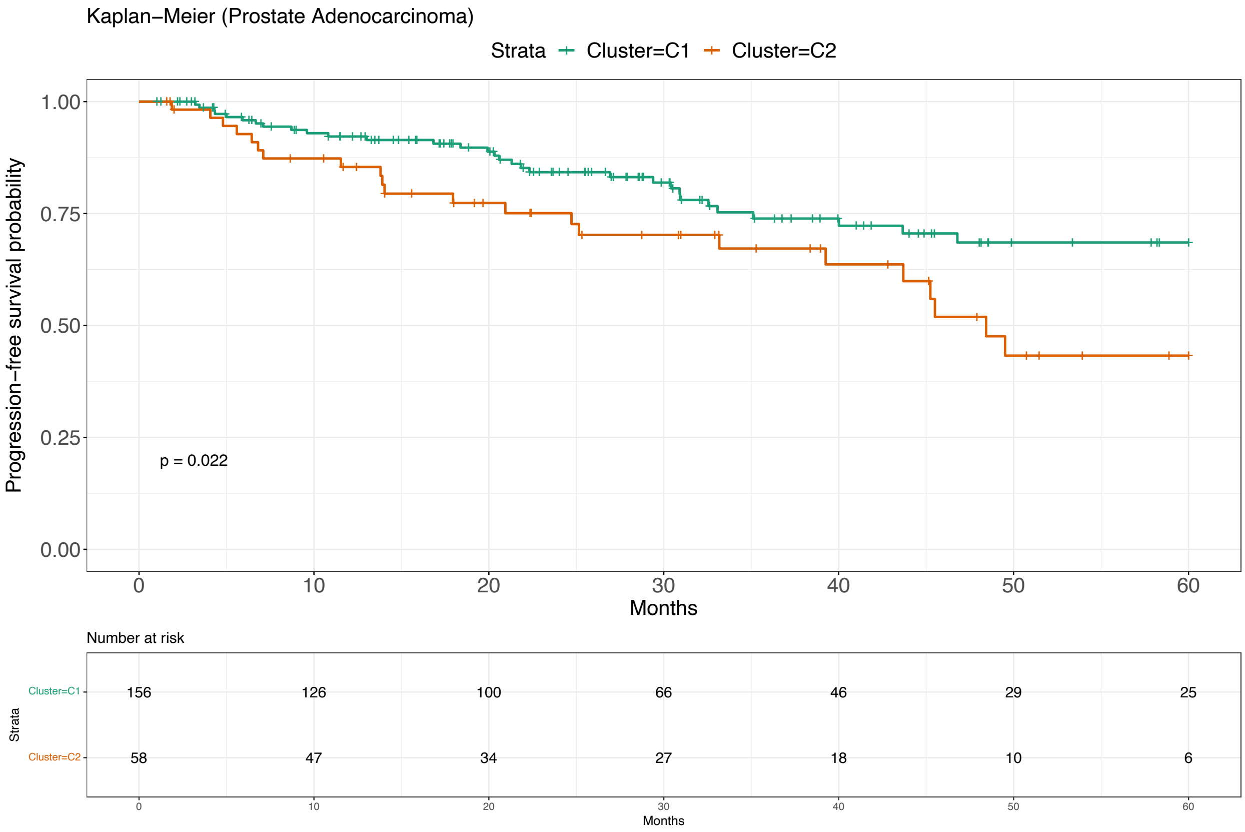

Figure 105: Survival analysis for Prostate Adenocarcinoma - Subtypes Validation (Pan-Cancer Atlas) [23].

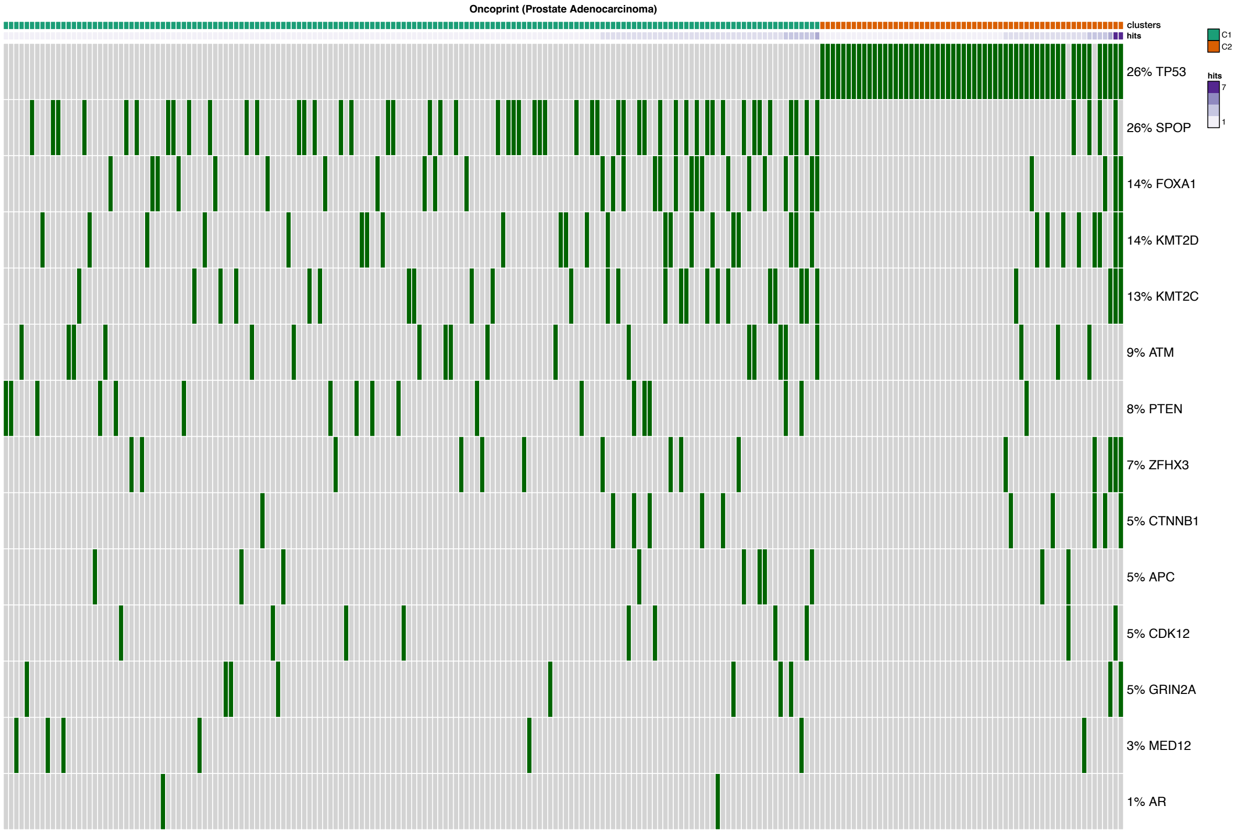

Figure 106: Oncoprint for Prostate Adenocarcinoma - Subtypes Validation (Pan-Cancer Atlas) [23].

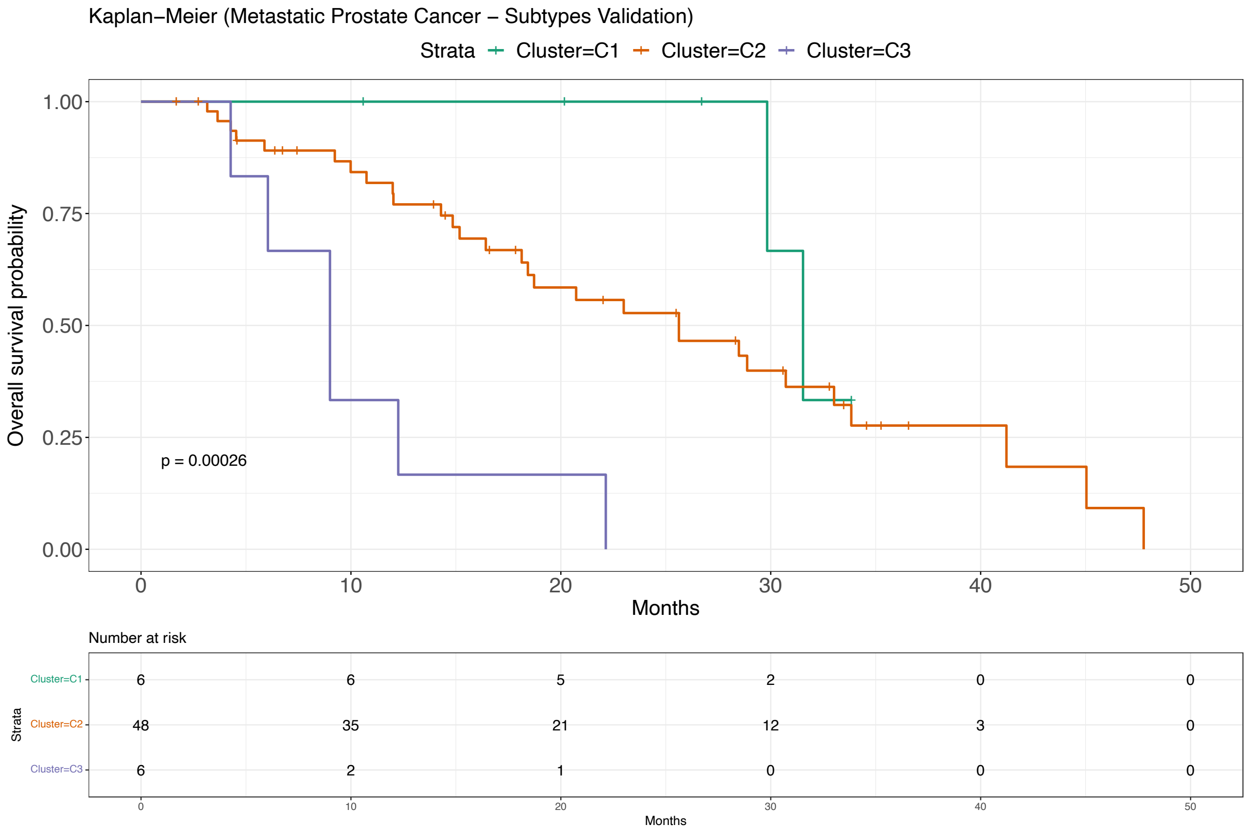

Figure 107: Survival analysis for Metastatic Prostate Cancer - Subtypes Validation (SU2C) [29].

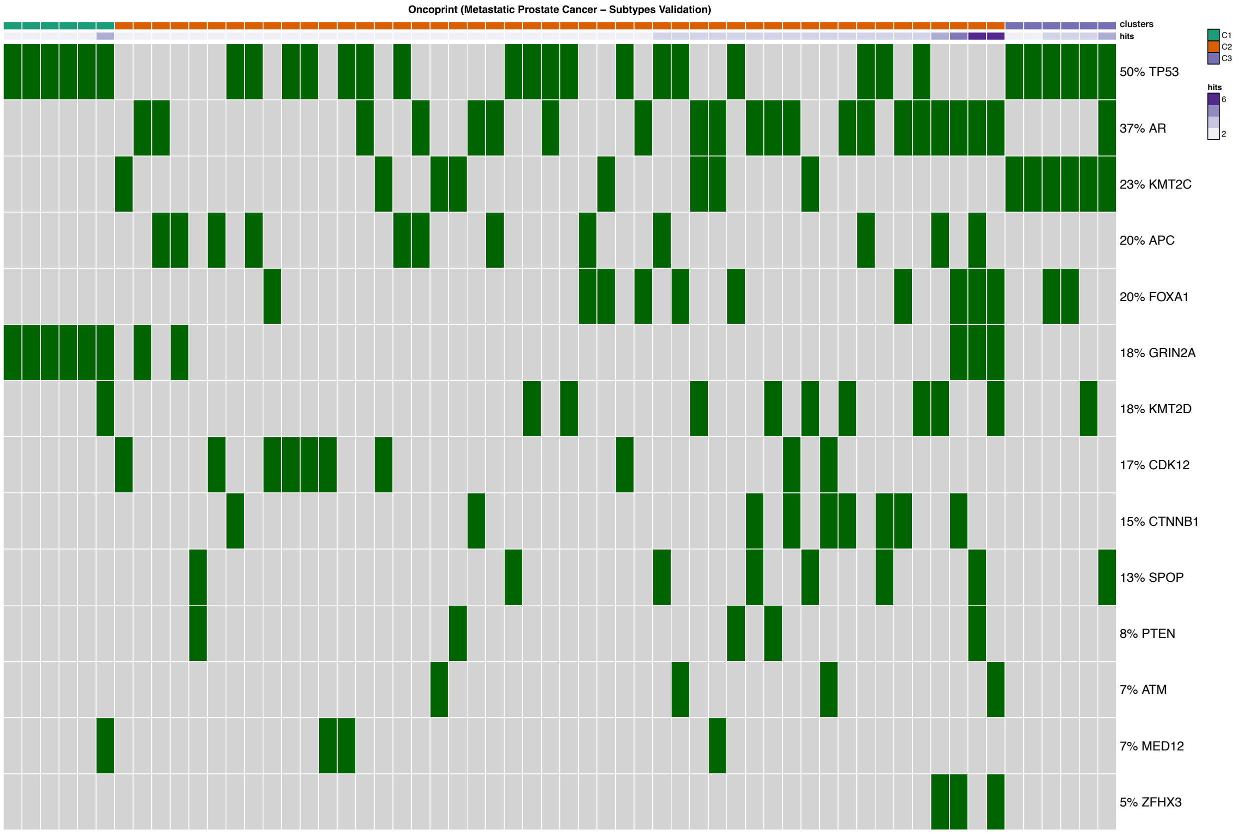

Figure 108: Oncoprint for Metastatic Prostate Cancer - Subtypes Validation (SU2C) [29].

[12] Jrgen Bang-Jensen and Gregory Gutin. Section 2.3. 4: The bellman-ford-moore algorithm. *Digraphs: Theory, Algorithms and Applications*, 2000.

[13] Nikolaj Tatti. Hierarchies in directed networks. In *2015 IEEE International Conference on Data Mining*, pages 991–996. IEEE, 2015.

[14] Patrick Suppes. A probabilistic theory of causality, 1970.
